# Multitrait genetic-phenotype associations to connect disease variants and biological mechanisms

**DOI:** 10.1101/2020.06.26.172999

**Authors:** Hanna Julienne, Vincent Laville, Zachary R. McCaw, Zihuai He, Vincent Guillemot, Carla Lasry, Andrey Ziyatdinov, Amaury Vaysse, Pierre Lechat, Hervé Ménager, Wilfried Le Goff, Marie-Pierre Dube, Peter Kraft, Iuliana Ionita-Laza, Bjarni J. Vilhjálmsson, Hugues Aschard

## Abstract

**Background:** Genome-wide association studies (GWAS) uncovered a wealth of associations between common variants and human phenotypes. These results, widely shared across the scientific community as summary statistics, fostered a flurry of secondary analysis: heritability and genetic correlation assessment, pleiotropy characterization and multitrait association test. Amongst these secondary analyses, a rising new field is the decomposition of multitrait genetic effects into distinct profiles of pleiotropy.

**Results:** We conducted an integrative analysis of GWAS summary statistics from 36 phenotypes to decipher multitrait genetic architecture and its link to biological mechanisms. We started by benchmarking multitrait association tests on a large panel of phenotype sets and established the *Omnibus* test as the most powerful in practice. We detected 322 new associations that were not previously reported by univariate screening. Using independent significant associations, we investigated the breakdown of genetic association into clusters of variants harboring similar multitrait association profile. Focusing on two subsets of immunity and metabolism phenotypes, we then demonstrate how SNPs within clusters can be mapped to biological pathways and disease mechanisms, providing a putative insight for numerous SNPs with unknown biological function. Finally, for the metabolism set, we investigate the link between gene cluster assignment and success of drug targets in random control trials. We report additional uninvestigated drug targets classified by clusters.

**Conclusions:** Multitrait genetic signals can be decomposed into distinct pleiotropy profiles that reveal consistent with pathways databases and random control trials. We propose this method for the mapping of unannotated SNPs to putative pathways.

## Main

Genome-wide association studies (GWAS) have identified thousands of significant genetic associations for multiple traits and diseases^1^. Publicly available summary statistics from these GWAS have proven invaluable in human genetic studies, enabling a range of secondary analyses without requiring individual-level genotype data and thus, averting major practical and ethical issues^2^. Among others, the estimation of phenotype heritability^3^, the derivation of polygenic risk score^4^, and the assessment of causal relations between phenotypes^5^ are paragons of their critical utility. GWAS summary statistics have also been extremely useful to investigate pleiotropy and the genetic relationship between human phenotypes. For example, recent works assessed whether significant loci for a given phenotype are also associated with other traits^6,7^ while others estimated genome-wide^8,9^ and regional^10^ genetic correlations among phenotypes. The joint analysis of multiple traits is also an efficient way to detect variants missed by univariate screening^11–23^, especially variants with association patterns that deviate from the observed phenotypic correlation^24–26^. Nevertheless, while simulation studies and examples from real data applications in best case scenarios have confirmed the relevance of multitrait association tests, there have seldom been applied to large-scale dataset.

Here, we argue that, besides the detection of new associated variants, multitrait GWAS summary statistics analysis offers a powerful framework to decipher the complex inter and intra-phenotype genetic architecture. We performed series of analyses on GWAS summary statistics from 36 phenotypes categorized into five clinically relevant sets (*Immunity, Anthropometry, Metabolism, Cardiovascular* and *Brain*) that demonstrate how such data can be used to reveal potential genetic pathways and their links to diseases. First, characterizing and comparing the relative performances of alternative multitrait association models, we found strong specificity of the signal identified by each approach, both in terms of association patterns and expressed tissue enrichment. We then used a Gaussian mixture model on the phenotypes by variants association matrix to identify potential clusters of variants displaying similar genetic multitrait association profiles. In-depth functional analysis of the resulting clusters demonstrates a connection between those profiles and tissue specific expression. This breakdown of multitrait association signal highlighted how the overall genetic correlation between phenotypes can be decomposed into likely distinct genetic pathways. Finally, we used the phenotypes from the *Immunity* and *Metabolism* sets as case studies to demonstrate the matching between the identified profile and known biological pathways. Noteworthy, mapping SNPs with unknown functions to pleiotropy profiles can indicate putative pathways. We conclude by investigating the potential clinical utility of the identified clusters for drug targeting.

## Results

### Multitrait genetic association signal

We analyzed the 36 GWAS studies of European ancestry (**Tables S1** to **S3**) using two approaches applied to seven phenotype sets: five medical-based sets (*Immunity, Anthropometry, Metabolism, Cardiovascular* and *Psychiatric*), a BMI related set including anthropometry traits and lipids (referred further as the Composite set), and finally all 36 phenotypes jointly (Fig. 1). Note that, we included Bone mineral density traits in the immunity set because an enrichment of BMD genome wide significant loci in immune pathways and immune cell regulatory regions has been previously reported^27,28^. The first step of our study consisted in maximizing the number of associated genetic variants by performing multitrait association tests using existing methods. In brief, we denote the single nucleotide polymorphisms (SNP) vectors of *Z*-scores, = (*z*_1_,…,*z*_*K*_), where *K* is the number of phenotypes (i.e. the number of GWAS analyzed jointly). The first model we used, which we refer to as *sumZ*, assumes that genetic effects across the phenotype analyzed follow a prior direction specified by a vector of weights w, to form a weighted sum of *Z*-scores. Here we considered four weighting schemes: i) uniform weighting (*sumZ*_*1*_); ii) weighting according to the first principal component of the phenotypic correlation matrix (*sumZ*_*r*_); iii) weighting according to the first principal component of the overall genetic correlation matrix (*sumZ*_*g*_); and iv) weighting according to the independent component analysis of the Z-scores matrix (*sumZ*_*ica*_). The second approach, which we refer to as *omnibus*, does not rely on prior specification on the direction and/or magnitude of the SNP effect across traits. In brief, it compares, for one SNP, the vector of genetic effects **z** with the expected multivariate normal distribution under the null. It is a standard omnibus test based on summary statistics that allows for one degree of freedom *per* outcome (here *per* phenotype). We performed in-depth validation of each approach using both simulation and real data from the UK Biobank cohort, characterizing their robustness (**Figs S1** to **S3**) and their link to methods based on individual-level data (**Figs S4** to **S6** and **Supplementary Note**). We also developed corrections for several critical real data issues related to model misspecification (**Figs S7** to **S12**) and missing data (**Fig.S13**).

**Figure 1.**
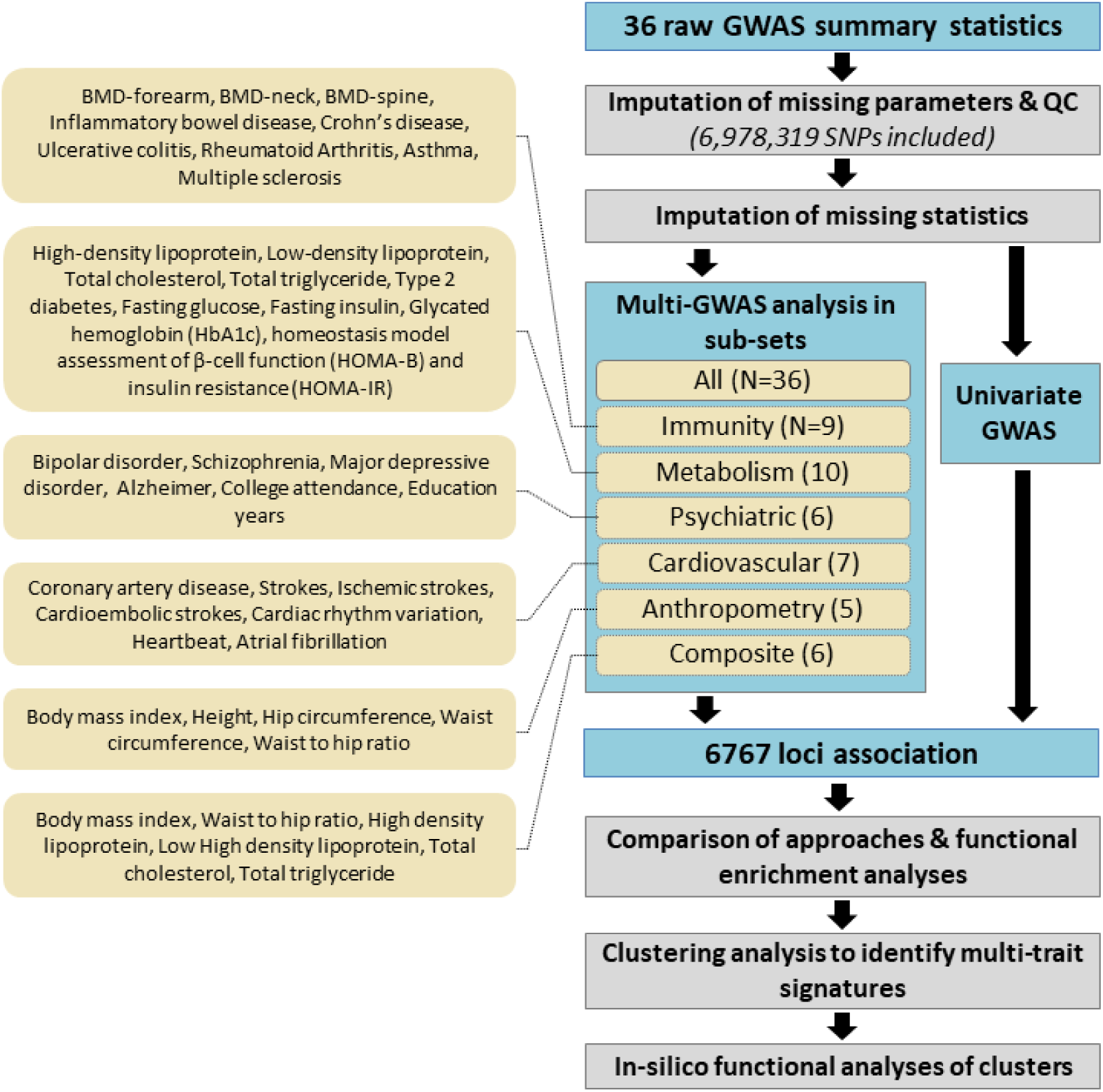
Analysis overview. The diagram below presents the overall analysis pipeline. A total of 36 GWAS were included covering several common diseases and quantitative traits. All GWAS summary statistics went through extensive pre-processing and quality control filtering, and missing single SNP statistics were imputed when possible. Multitrait approaches were then applied to all clean GWAS data and on each clinically based set (All, *Immunity, Metabolism, Brain, Cardiovascular, Anthropometry*, and *Composite*). After combining univariate and multivariate results, and merging SNPs within locus, a total of 6,767 associations were identified. After a comparison of results per approach, a clustering analysis was performed for variants within each set. Finally, we performed in-silico functional analysis of the clusters derived in the *Metabolism* set to assess their biological relevance.

To empirically determine the detection ability of each approach, we derived the overlap of significant loci of the multitrait tests per phenotypes set (**Figs S14** to **S20**), and after merging all analyses (Fig. 2A). Univariate phenotype association were included in the comparison using the minimum of univariate p-value across all outcomes (noted *P*_*univ*_). Across all phenotype sets, 391 associations were identified by the multitrait tests only, 392 were identified by univariate association tests only, and 1557 were significant for both univariate and multitrait tests (see Fig. 2A). The largest number of new associations were detected by the *Omnibus* test. The performances of the *sumZ* tests varied substantially depending on the phenotype set. For example, the weighting scheme based on phenotypic correlation (*SumZ*_*r*_), detects slightly more signals than other weights for the *Immunity* set (**Fig. S18**) but fewer associations in other phenotype sets (Fig. 2A). While the Omnibus detected the largest number of new associations, the substantial share of signals found by other models suggests that applying several multivariate tests, especially the combination *omnibus*, *sumZ*_*ica*_, *sumZ*_*g*_, could be an interesting solution to maximize detection. Finally, we checked the 392 associations identified by the multitrait test only in this data against previously reported associations from the GWAS catalogue^1^ for the same phenotypes. Altogether, we report a total of 322 new associations (**Tables S4** to **S10**).

**Figure 2.**
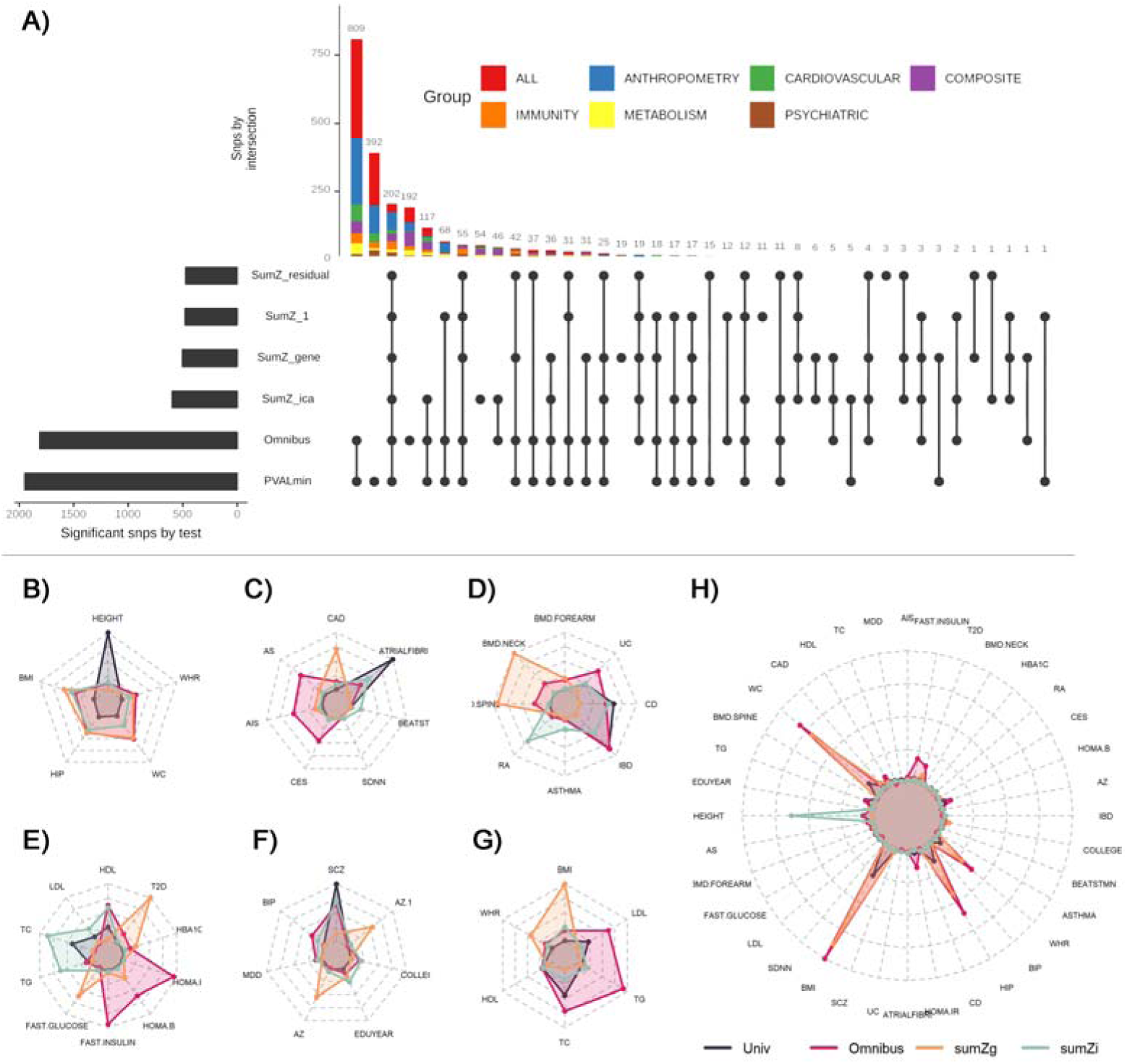
Multitrait approach comparison. Panel (**A**) shows independent variants detected across the six approaches: univariate test (*univ*), omnibus test (*omni*), weighted sum of Z-score with uniform weight (*sumZ*_1_), weight defined as the loading of the first principal component of the phenotypic correlation (*sumZ*_*r*_), the genetic correlation (*sumZ*_*g*_), or defined using the loadings of an independent component analysis (*sumZ*_*ica*_). Each line corresponds to a test and each column to a set of significant variants. For each set, the test for which variants are significant are represented with a black dot on the test line. The barplot at the left represents the total number of significant independent signals detected by each approach. The stack bar at the top represents the cardinality of the sets. The next panels show the link between strengths of univariate association signal and the relative performance (i.e. larger power) of the four most tests: univ, omni, *sumZ*_*g*_, and *sumZ*_*ica*_, for each phenotype set: *anthropometry* (**B**), *cardiovascular* (**C**), *immunity* (**D**), *metabolism* (**E**), *brain* (**F**), *composite* (**G**), and *all phenotypes* (**H**). Within each phenotype set, we split the top associated SNPs per region based on the most significant test, and derived the median chi-squared for each test. The radar plots show the derived median per test and illustrate the strong heterogeneity in patterns identified. For example, out of the 1605 SNPs from the *anthropometry* set, 1235 had stronger signal with univ as compared with other tests. The median chi-squares in that group were 49.1, 1.1, 2.0, 1.0, and 0.7 for height (Height), body mass index (BMI), hip circumference (HipC), waist circumference (WaistC), and waist to hip ratio (WHR). Comparatively, the 267 SNPs harboring a stronger signal with *omnibus*, had median of 6.8, 20.1, 15.9, 11.2, and 7.2 for the same phenotypes.

To understand further the relative performance of those three tests (*omnibus*, *sumZ*_*ica*_, *sumZ*_*g*_) along the univariate test, we explored which multitrait signal was associated with the largest increase in detection per test. For that aim, we listed all loci found associated with at least one of the four approaches, and assigned each locus to a test based on the lowest *p*-value. We then derived the median chi-squared z-score by phenotype across the loci assigned to each test. As showed in Fig. 2B-H, the median pattern varied substantially across tests and phenotype sets. Higher power for the univariate test was, as expected, observed for strong association signals for a single phenotype, and mostly reflected a very large sample size for that phenotype and/or a strong heritability (e.g. height in the anthropometry set, Fig. 2B, or atrial fibrillation in the cardiovascular set, Fig. 2C). Strong association signal for the *omnibus* test was linked to the inclusion of correlated phenotypes and sample overlap resulting in a high residual covariance (**Σ**_r_, **Table S2**). For example, median chi-squared were elevated for the any strokes (AS), any ischemic strokes (AIS) and cardioembolic strokes (CES) in the cardiovascular set. The pattern preferentially detected by the *sumZ*_*g*_ test are harder to interpret. Yet, we notice that *sumZ*_*g*_ displays strong signal for SNPs associated with physiologically related traits (e.g. T2D and fasting glucose in the metabolism set, Fig. 2E, or bone mineral density of neck and spine in the immunity set, Fig. 2D).

To confirm the relevance of association detected by multivariate tests, we also conducted a tissue enrichment analysis to significant variants identified by the multitrait approaches and by the univariate analyses separately (**Tables S11** and **S12**). Overall, univariate variants and with multitrait variants harbored a very similar functional enrichment landscape (**Fig. S21**). Most enriched tissues are already known to be involved in the phenotype in question, including for example liver, fat and pancreas for the *Metabolism* set, immune cell types and thymus for the *Immunity set*, and heart for the *Cardiovascular* set. Our enrichment study also confirmed less obvious observations, which have nevertheless been noted before: the involvement of immunity in brain-related traits (e.g. autisms and schizophrenia)^29,30^ and the over-representation of brain tissues in the *Metabolism* set^31,32^.

### Distinct genetic association profiles correspond to distinct genetic correlation

Our comparison of approaches highlights that associated genetic variants display a broad range of multitrait association profiles. We investigated how these profiles can be broken down into groups of homogeneous multivariate genetic effects. This is directly related to the principle of genetic correlation, which quantifies the concordance of genetic effects across traits (e.g. ^9^). The difference here, is that genetic correlation captures only the average over the whole genome, and as discussed in recent studies, more localized genetic structures likely exist for many pairs of traits^10^. To detect such structure, we implemented a multivariate Gaussian mixture model (MGMM)^33^ for the identification of clusters among SNP found associated with at least one approach. We applied MGMM assuming between 2 to 10 clusters and use the BIC and silhouette criteria to determine the most relevant number of clusters. We further bootstrapped the computation of the clustering criteria to ensure the robustness of the selection (**Supplementary Material**). The best suited number of clusters is 6, 8, 8, 9, 3, 2 and 5 for the *Metabolism, Immunity, Cardiovascular, Anthropometry, Psychiatric, Composite*, and *All sets*, respectively (**Fig. S22**). As illustrated for the *Metabolism* set in **Fig. S23**, adding significant SNPs from the multitrait tests on top of those identified by the univariate tests enabled us to detect more clusters.

The resulting clustering are presented in Fig. 3 for the *Metabolism* set and in **Figs S24** to **S30** for the other sets. Each figure presents a heatmap of *Z*-scores along with an alluvial plot displaying both the shared explained variance between phenotypes and the proportion of explained variance by clusters for each phenotype. The complete list of SNP for the *Metabolism* set per cluster is presented in **Table S16**. The multivariate effects vary substantially from one cluster to another. For instance, in *Metabolism* clusters, SNPs from the cluster 3 display increased HDL-C and decrease triglycerides, while SNPs from cluster 5 are more specific to triglycerides. We ensured the uniformity of the multitrait association profiles inside clusters by filtering out SNPs with uncertain cluster assignation (*i.e.* those with entropy above 0.75, see Fig. 3C and **Supplementary Material**).

**Figure 3.**
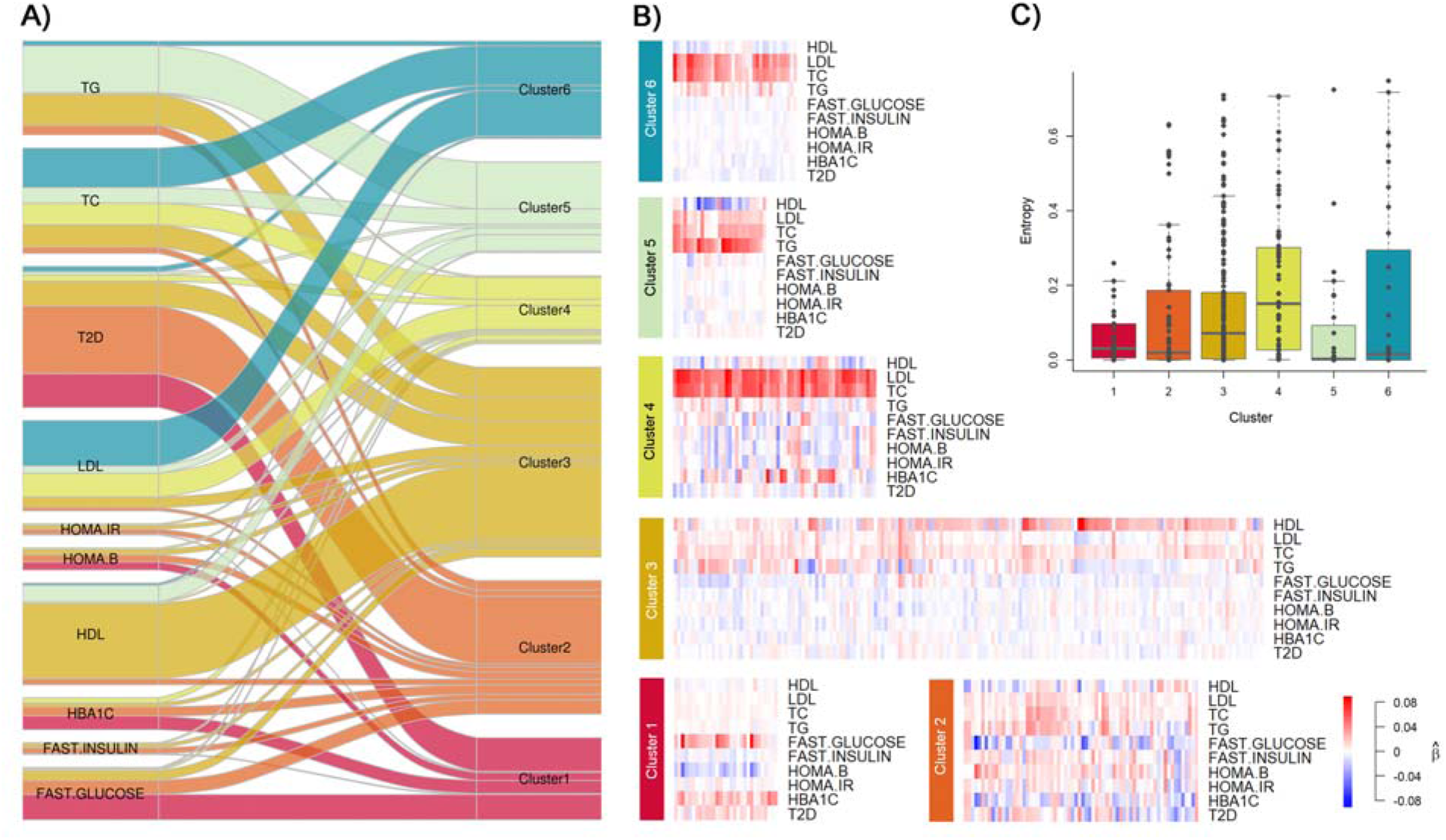
Multitrait genetic association clusters for the *Metabolism* set. The panels summarize the clustering of the 392 independent SNPs selected from the *Metabolism* set analysis. The set includes 10 phenotypes: triglyceride (TG), total cholesterol (TC), type 2 diabetes (T2D), low-density lipoprotein cholesterol (LDL-C), high-density lipoprotein cholesterol (HDL-C), glycated hemoglobin (HbA1c), Homeostasis model assessment of []-cell function (HOMA-B), homeostasis model assessment of insulin resistance (HOMA-IR), fasting insulin, and fasting glucose. The alluvial plot in panel **A**) represents the decomposition of univariate genetic association and its rewiring to the six inferred clusters. The flow widths represent the proportion of phenotype’s variance explained by the subset of SNPs assigned to each specific cluster, relative to the total genetic variance explained by all 392 SNPs. For example, SNPs from cluster 6 capture approximatively 41.7% and 54.6% of that genetic variance for TC and LDL, respectively. For clarity, flows explaining less than 0.1% of the variance are not represented. Panel **B**) shows the heatmap of normalized beta coefficients per phenotype within each cluster. Each column is a SNP, with blue and red colors indicating negative and positive beta, respectively. Coded alleles have been defined according to the per cluster first principal component. The boxplots in panel **C**) shows the distribution per cluster of SNP’s entropy, an indicator of the fitness of the SNP-cluster assignment. SNPs perfectly assigned are expected to have entropy close to zero.

The alluvial figures and heatmaps provide an overview of the magnitude of genetic effect from one cluster to another. To further characterize concordance or discordance of genetic contributions across phenotypes, we computed the pairwise SNP-based genetic correlations for each cluster (see **Supplementary Material**). Fig. 4 presents those estimates for a subset of phenotypes within the *Metabolism* and *Immunity* phenotype sets. In the *Immunity* set, the correlations between Rheumatoid Arthritis (RA), Ulcerative Colitis (UC) and Crohn disease (CD) provide a striking illustration how the genome-wide genetic correlation can be composed of smaller structures. The genome-wide genetic correlations between UC and CD is strong (0.41), but near 0 and not significant for RA (see **Table S3**). In Fig. 4B, we can yet notice a fairly large negative correlation in cluster 2 and 3 between RA versus CD or UC, whereas, the cluster 5 captures a group of variants displaying strong positive correlation across the three traits. Similar negligible genome-wide correlation along opposite genetic correlation across clusters are observed in the *Metabolism* set. For example, variants from cluster 1 display strong concordant effect between LDL and T2D, but variants from cluster 6 harbor an equally strong negative correlation. Fig. 4 also highlights that significant genome-wide genetic correlation across highly related phenotypes such as UC-CD and LDL-TG are not distributed evenly across variants.

**Figure 4.**
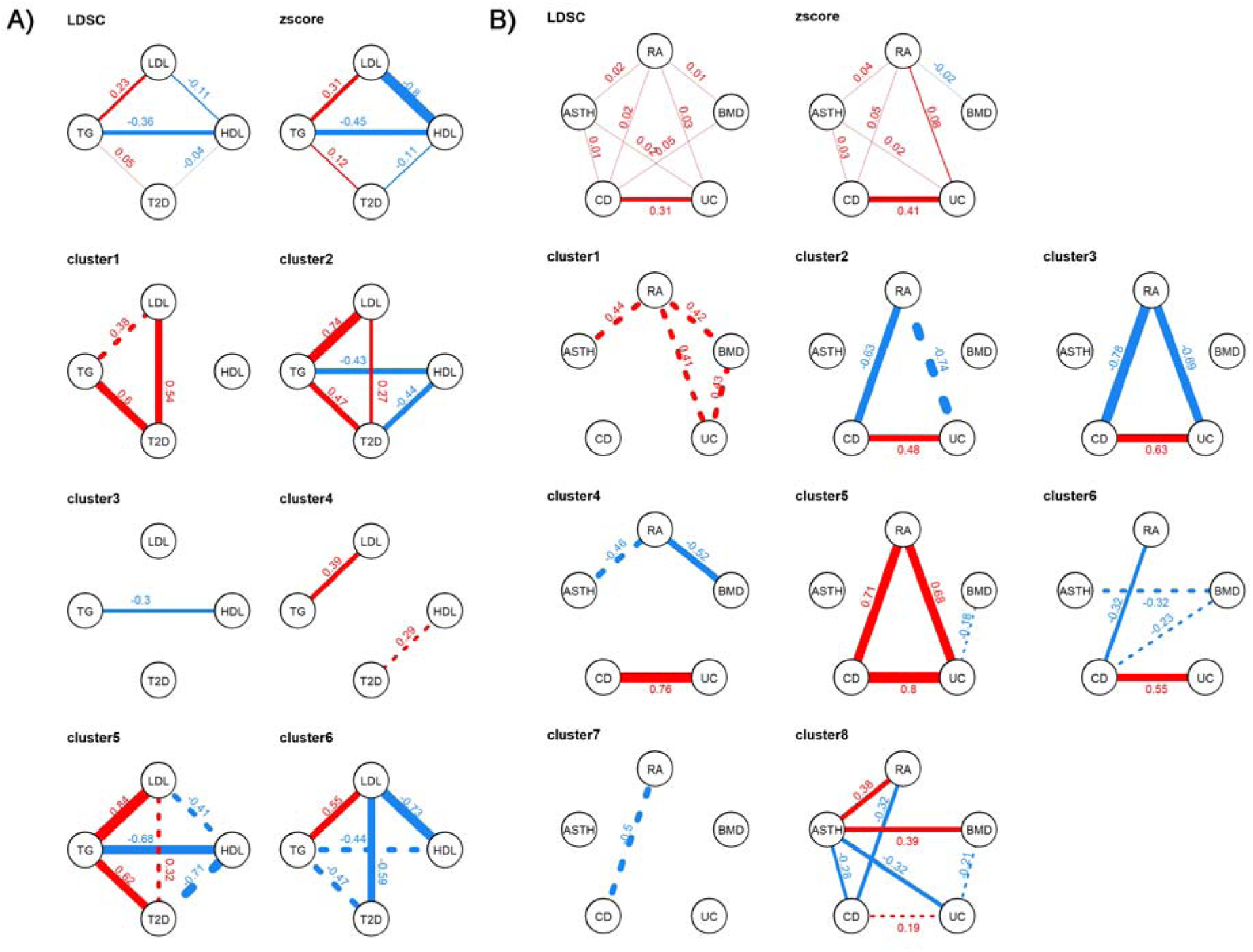
Heterogeneity of genetic correlation across clusters for the *Metabolism* and *Immunity* sets. We derived the genome-wide genetic correlation between phenotypes using *LDscore* regression and using Pearson correlation from all SNP Z-scores (top panels), and for SNPs within the identified clusters. Results for the *Metabolism* set are presented in panel (**A**) using only the four key traits, LDL, HDL, Triglyceride (TG) and type 2 diabetes (T2D). Results for the *Immunity* set are presented in the panel (**B**). For clarity only significant correlation are represented. The boldness of the line is proportional to the strength of the genetic correlation. Positive correlations are represented in blue and negative correlations in red. The values of the genetic correlation are indicated by the blue number next to the trait. Solid lines represent significant correlation (after Bonferroni correction) whereas dashed lines represent correlation significant only before Bonferroni correction. Note that because the clusters are inferred from the multivariate associations, the absolute value of the significance of the correlations is of limited interest. Nevertheless, it provides a useful descriptive statistic to identify the key structures within each cluster.

### Biological meaning of genetic clusters

These distinct multitrait association profiles might arise because their variants belong to distinct genetic functional groups. Understanding whether those genetic functional groups are only statistical construction or correspond to meaningful biologically mechanism is critical. In the latter, it means that data-driven approach, such as the one proposed in the present study, can be used to dissect the genetic contribution of many complex human phenotypes. To assess this hypothesis, we conducted series of *in silico* functional analyses with the objective of mapping clusters to candidate biological functions. For each phenotype set, we evaluated two types of enrichment: tissue-specific chromatin mark enrichment per cluster (**Table S13**), and pathway enrichment framework (**Tables S14** and **S15**) which integrates multiple databases such as Gene Ontology (GO) and KEGG. Here, we focused on the *Immunity* and *Metabolism* sets as a case study.

For the *Immunity* set, clusters 1 and 4 are predominantly capturing genetic effect on bone-mineral density; clusters 2, 3 and 5 effect on inflammatory bowel disorder (IBD); and clusters 6, 7 and 8 capture variants with pleiotropic effects on rheumatoid arthritis and IBD (**Fig. S26**). Both enrichment analyses pointed toward an overrepresentation of the immune system with all clusters –even the ones affecting primarily bone-mineral density– being enriched for at least one immunologic pathway or one immunological tissue. We highlight the top enriched tissues and top pathways in Table 1. Concerning pathway enrichment, immune related pathways regulating the shape of the immune response such as cytokines and the JAK-STAT signaling pathway were recurrent. Interestingly, variants from those clusters map to a distinct set of cytokines and cluster of differentiation genes (e.g. IL4, IL13, IL33 for cluster 1 and IL3, IL5, IL10, IL19, IL20, IL21, IL27 for cluster 5) which suggests that they may impact different components of the immune system. Concerning tissue-specific active chromatin mark enrichment, clusters 2 and 3 contain multiple SNPs enriched primarily in transcriptionally active regions of “Primary Natural Killer cells from peripheral blood” whereas cluster 7 and 8 are enriched for “Primary T helper cells.” We also observed enrichment in the tissue where the immune damages occur for the cluster 5 (colonic mucosa) which highlight the complex interaction between the immune system and the inflamed tissue.

**Table 1.**
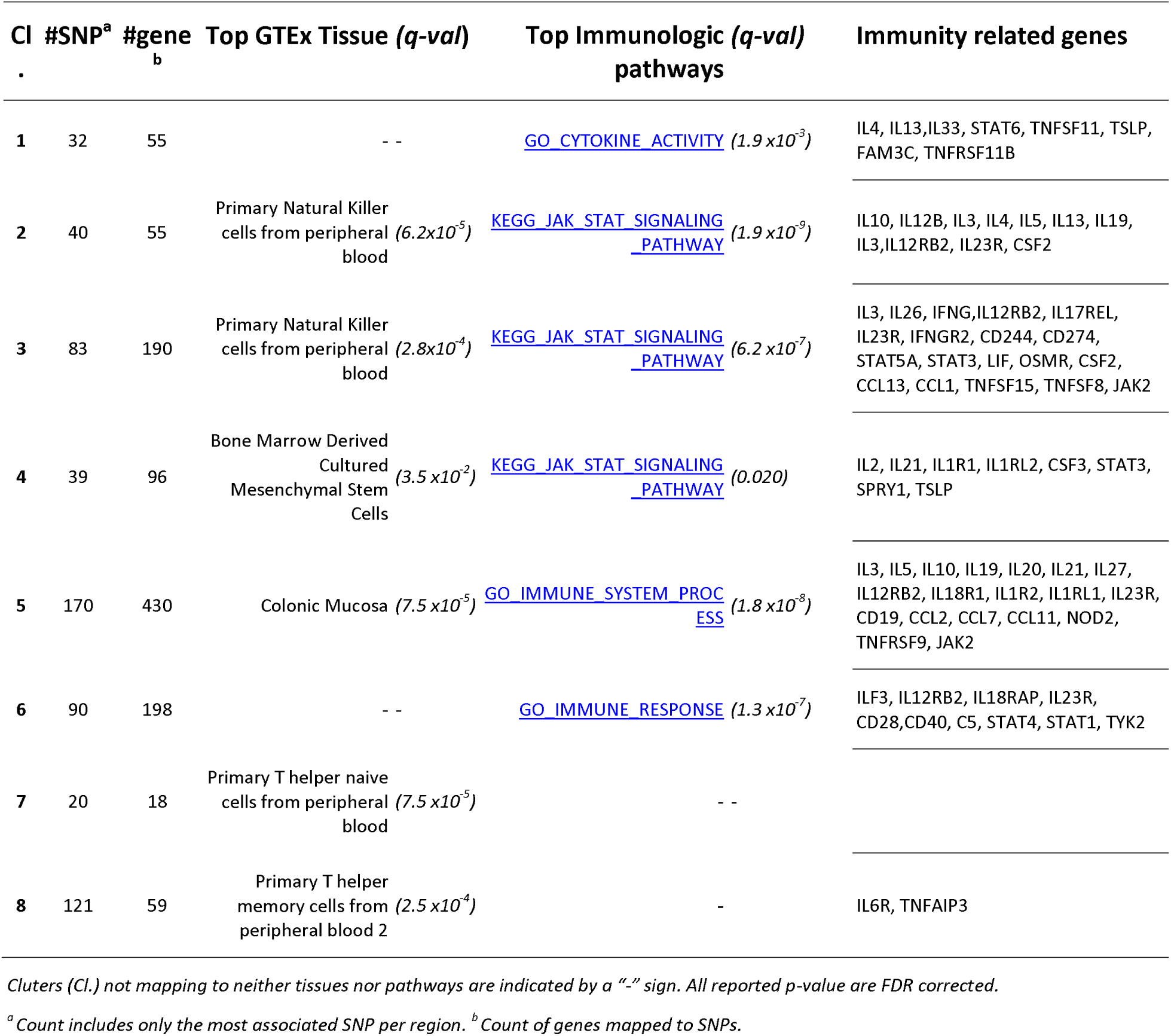
Top tissue associations and Immune related Genes by Clusters for the Immunity set.

The *Metabolism* set includes several molecular phenotypes, which we expect to be closer to biological mechanisms than some of the macro-phenotypes from other sets. Overall, cluster 1 is mostly associated to an increase of fasting glucose and an impaired β-cell function; cluster 2 is highly pleiotropic and notably increases the risk of T2D, clusters 3 to 6 are mostly associated with lipids, and with LDL-TC, HDL-TC-TG (Fig. 3). Accounting for the direction of effects, we also note that the genetic associations in cluster 5 match the known phenotypic correlation with the inverse relationship between circulating levels of HDL-C with those of LDL-C and more especially TG observed in epidemiological studies^34^. At the tissue level, we observed modest enrichment for adipocytes in clusters 1 and 2 (FDR *p*-value 0.028 and 0.01 respectively, **Table S13**) and cluster 3 SNPs are up-regulated in the Liver (FDR *p*-value 0.005).

As shown in **Table S14**, each cluster was significantly enriched for a large number of GO terms. We report some specific and illustrative examples: cluster 1 is enriched for carbohydrate homeostasis set (*q*-value= 2.5 × 10^−3^), cluster 3 is enriched for reverse cholesterol transport set (*q*-value= 2.8 × 10^−13^), cluster 4 is enriched for plasma lipoprotein clearance set (*q*-value= 1.7 × 10^−5^), cluster 5 is enriched for protein lipid complex assembly set (q-value= 1.08×10^−9^) and cluster 6 is enriched for low density lipoprotein particle remodeling set (*q*-value= 1.07×10^−2^). Cluster 4 also exhibits active chromatin tissue enrichment in immune T cells (q-value=2.3×10^−3^), highlighting the link between cholesterol and *Immunity*. Indeed, cholesterol as well as modified forms of cholesterol such as oxidized cholesterol and cholesterol crystals, promote inflammatory and immune responses through multiple pathways including activation of the Toll-like receptor (TLR) signaling, *NLRP3* inflammasome and myelopoiesis^35,36^. While the promotion of inflammation and immunity is carried by LDL particles, HDL particles were proposed to counteract this effect in part through reverse cholesterol transport^37^. However, cluster 3 which is enriched for reverse cholesterol transport did not exhibit such a tissue enrichment in immune T cells indicating that the link between HDL and *Immunity* may harbor more complexity, as recently pointed by out by Madsen et al^38^.

### Metabolism pathways and diseases

To provide a perspective on the specificity of genetic variants across clusters and their potential contribution to human diseases, we investigated the lipids from the *Metabolism* set. We first projected each cluster gene onto KEGG pathways. Here, we used only maps corresponding to enriched GO gene sets identified or to tissue identified in the enrichment analysis at the previous stages (**Tables S14** and **S15**): fat digestion and absorption, cholesterol metabolism, PPAR signaling pathways. We constructed a synthesis of these observations on the metabolic map presented on Fig. 5A-B. Genes associated to clusters (**Table S16**) had functions in agreement with their effects on blood lipid levels: cluster 3 (HDL-C++) is enriched in genes involved in HDL-C biogenesis and metabolism (*LCAT, ABCA1, SR-B1, CETP, PLTP, LIPG, APOAx* and *APOCx*), clusters 4 and 6 with genes related to LDL-C clearance (*SORT1, PCSK9, LDLR, LDLRAP1, APOB* and *APOE*), and cluster 5 to genes related to triglycerides and chylomicron transport (*LPL, APOAx* and *APOCx*).

**Figure 5.**
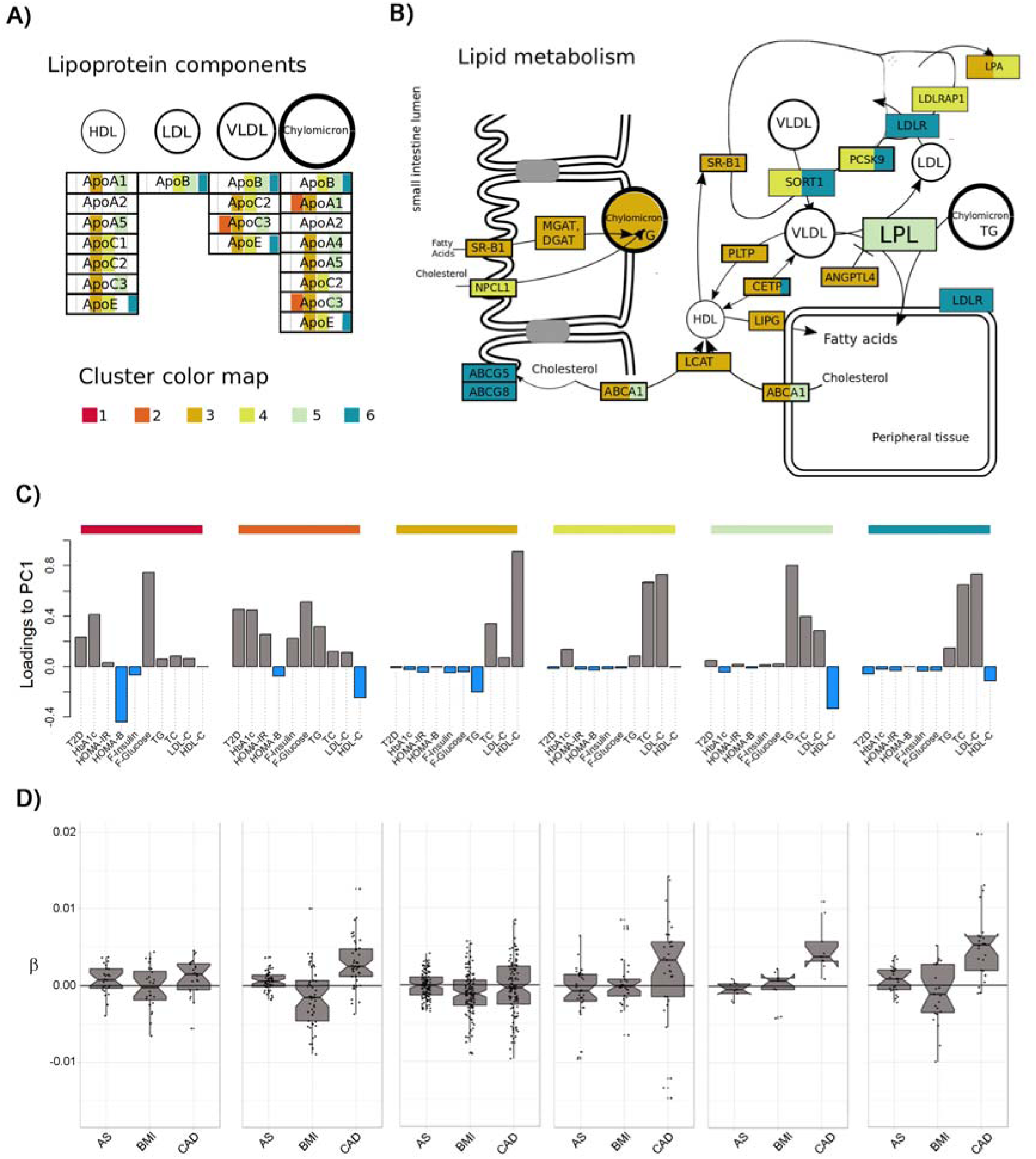
Mapping clusters to pathways. We projected cluster’s genes from the *Metabolism* phenotype set onto KEGG pathways and reconstructed a synthetic metabolic map. Panel **A**) presents the results for the lipoprotein component and panel **B**) for the lipid component. Gene names are highlighted by the colors of their associated clusters. When a gene is associated to several SNPs belonging to different clusters it is represent with several colors. To improve interpretation, we also present in panel **C**) a proxy for the relative contribution of each phenotype per cluster, defined as the loadings of the first principal component derived from the matrix of Z-score for the subset of SNPs in that cluster. Finally, panel **D**) shows the distribution of standardized beta for association between SNPs from each cluster and three diseases: any stroke (AS), coronary artery disease (CAD), and obesity (using body mass index as a proxy).

We then assessed the effect of variants from each cluster with three diseases known to be associated with serum lipids: coronary artery diseases (CAD), stroke, and obesity (defined as a BMI > 30) (**Table S19**). Within each cluster, we aligned the SNPs alleles with the main trend of the corresponding cluster, so that all coded alleles fit the multitrait pattern defined in Fig. 5C (see **Supplementary Material**). For example, all SNPs from cluster 5 were re-coded to be associated with an increase in TG, TC and LDL-C, and a decrease in HDL-C. We plotted in Fig. 5D and in **Fig. S31** the genetic effect of each SNP on the three diseases (using effect on BMI as a proxy for obesity) after the aforementioned alignment, and performed a sign test to assess the significance of the observed trend (**Table S19**). SNPs from several clusters display a significant increase in risk of CAD: cluster 2 (*P*=6.6×10^−5^), cluster 4 (*P*=2.9×10^−2^), cluster 5 (*P*=3.9×10^−3^) and cluster 6 (*P*=2.8×10^−4^). SNPs from cluster 2 also display a nominally significant increase in risk of stroke (*p*-value =1.6×10^−2^). Finally, a large fraction of SNPs from cluster 3 has negative effect on BMI (*P* =6.4×10^−4^). Interestingly, several SNPs from this cluster show association with CAD, but with heterogeneous effects –some associated with an increased risk and other associated with a decreased risk– so the absence of a global trend. The associations of cluster 4 and 6 with CAD add to the evidences of a causal effect of LDL-C on CAD^39^, which has been established by prospective epidemiological studies^40^, mendelian randomization^41^ and randomized clinical trials evaluating the effect of LDL-C reducing therapies^42^. Moreover, cluster ^5^ association to CAD risks corroborates a potential causal role of TG^5^ and remnant cholesterol^43,44^ on CAD. The role of TG in CAD has also been confirmed by epidemiological studies^45^, genome-wide association studies^5^, mendelian randomization studies^46^ and randomized controlled trials aiming the lowering of TG^47^. Cluster 3 which is associated with increases in HDL-C does not have a protective effect on CAD is again in agreement with mendelian randomization analyses reporting no link between HDL and CAD^41,48^. Finally, the association of cluster 2 with CAD and Strokes supports further the potential causal effect of type 2 diabetes on CAD and stroke^49^.

As a final exploratory analysis, we reported the cluster and multitrait genetic effect of genes targeted to mitigate hyperlipemia to prevent CAD (Table 2). It shows that drug target corresponding to the cluster 3 (*ABCA1, CETP, NR1H3*) did not lead to successful clinical trials whereas targets (*PCSK9, NPC1L1, APOC3, HMGCR*) in cluster 4, 5 and 6 are mostly successful or promising. The example of the *CETP* gene which is classified in cluster 3, a cluster not associated with CAD, is of particular interest. *CETP* has been the target of failed clinical trials which attempted to prevent CAD by inhibiting *CETP* and consequently increasing circulating HDL-C^50–52^. Cholesteryl ester transfer protein (*CETP*) promotes the heteroexchange of cholesteryl esters and TG between HDL-C and APOB-containing lipoproteins connecting HDL-C and TG *Metabolism*50. Pharmacological inhibition of *CETP* was motivated by GWAS^53^ and prospective cohorts^54^ that indicated that *CETP* variants were associated with higher circulating HDL-C levels, lower LDL-C, TG and CVD risk. However, although all *CETP* inhibitors achieved an effective increase in HDL-C, only *anacetrapib* led to a significant lower incidence of major coronary events^55^ in patient who were receiving statin therapy, an effect which might be accounted for the reduction of *ApoB* (non-HDL-C) rather than the elevation of HDL as suggested by mendelian randomization analyses^56^. Additionally to these well-known drugs, we provide a systematic listing of potential drug targets by cluster (**Table S20**) based on the druggable genome database^57^.

**Table 2.** Drug target genes and associated SNPs in the metabolism set

| Target<sup>a</sup> | Drug (phase) | rsID<sup>b</sup> | Clu. | SNP-phenotype association<sup>c</sup> |  |  |  |  |  |  | Comment |
| --- | --- | --- | --- | --- | --- | --- | --- | --- | --- | --- | --- |
|  |  |  |  | HDL | LDL | TC | TG | CAD | AS | BMI |  |
| ABCA1 | Probucol (4) | rs11789603 | 3 | 7.70 | 1.6 | 4.66 | 2.07 | 1.42 | 0.25 | -1.50 | This LDL-c lowering drug was approved but subsequently discontinued because of its lowering effect on HDL-c |
| CETP | Cetrapid (4) | rs12448528 | 3 | 27.79 | -4.61 | 4.96 | -4.60 | 0.25 | -0.28 | 1.21 | Three clinical trials were halted because they showed adverse effect and/or no therapeutic efficacy, except in the case of anacetrapid use for preventing new acute coronary events in high-risk individuals. |
| NR1H3 | HDCA (1) | rs12575609 | 3 | 9.11 | -0.19 | 1.89 | -3.26 | 0.76 | 0.06 | -3.9 | The RCT results were not produced due to AtheroNova Inc. bankruptcy. |
| PCSK9 | alirocumab ,<br>Evolocumab (4) | rs7523242 | 4 | -1.16 | 10.49 | 9.28 | 1.92 | 3.25 | 1.03 | -0.29 | Approved second line treatment for high cholesterol individuals whose cholesterol is not controlled by Statin alone. |
| NPC1L1 | Ezetimibe (4) | rs217386 | 4 | -0.80 | 6.60 | 5.96 | 2.44 | 2.19 | 0.86 | -1.47 | Currently used to lower the absorption of cholesterol and is often used in association with statin. |
| APOC3,<br>APOA1 | Volanesorsen (3) | rs1815787 | 5 | -2.09 | 5.45 | 9.41 | 16.60 | 0.39 | 0.26 | 0.614 | A triglyceride-reducing drug currently in phase 3 RCT. |
| HMGCR | Statins (4) | rs59014134 | 6 | 0.79 | 15.79 | 16.06 | 1.34 | 2.01 | -0.36 | -4.59 | The most common cholesterol lowering drugs. |
| APOB | Mipomersen (4) | rs1041968 | 6 | -6.96 | 22.94 | 20.92 | 9.38 | 2.45 | -1.20 | -1.75 | Can be used for risk management in familial hypercholesterolemia but can cause fatty liver disease. |

<sup>a</sup>Note that for probucol, the molecule inhibit ABCA1, but is not specific to ABCA1.
<sup>b</sup>Primary associated SNP and corresponding cluster. But note that for several loci, there is a few other SNPs from other cluster.
<sup>c</sup>Define as the association Z-score for the most associated variant in the gene..

Altogether, those results suggest that drug development might be more effective by accounting for the gene context, *i.e.* by selecting candidate gene not from their individual feature, but based on the disease association trend of genes displaying similar multitrait association profile. Under this working hypothesis, the proposed inference of genetic functional groups can provide a means to identify those genes and therefore to select potential candidates.

## Discussion

In this study, we conducted a multitrait analyses of GWAS summary statistics from 36 human phenotypes combining association tests and clustering to detect the shared and specific genetic substructure underlying those phenotypes, and explore the links between those substructures and biological pathways and diseases. The question of substructures underlying genome-wide genetic correlation has been partially explored in other recent studies^8,10^. Our work is in agreement with these studies, confirming the presence of regional genetic correlation differences and offering a data-driven approach for identifying primary substructures across millions of possibilities. Using two complementary functional enrichment analysis, we mapped these multitrait association profiles to pathways, and report a detailed view of these profiles for the *Immunity* and the *Metabolism* phenotype set.

The variability in pleiotropy profiles across identified GWAS SNP has been previously discussed. For example, earlier reports^58^ on inflammatory diseases have highlighted such patterns, or proposed grouping of SNPs based on the direction of association^59^. However, those studies used only a handful of SNPs identified at the time of publication. Our analysis based on a formal clustering and functional enrichment analyses, and using GWAS results perform in much larger sample size, offers a new and much more detailed qualitative perspective on these profiles. More recent publications have also discussed approaches focusing on the characterization of SNPs displaying pleiotropic effets^60^, the inference of shared and distinct genetic pathways between related phenotypes^61^, and on the identification of genetic components linked to disease subtypes^62,63^. Our approach shares objectives with some of these methods but has also unique features and advantages. Approaches that rely on individuals’ genotypes are limited by the ethical and practical cumbersome aspects tied to this type of data^63^. Studies based on component decomposition techniques alike principal component analysis^61,62^, while being efficient as data compression techniques, yields endotypes based on components that are of interest from a biological standpoint, but do not provide the SNP-level genetic decomposition that we are addressing.

Past studies showed that sufficiently curated genetic information can enhance the chance of success of clinical trials^64,65^. We further argue that fine analysis of pleiotropic effects, as performed in the present study, is a very promising path forward to help identifying drug targets with a minimal risk of serious side effects. In particular, the picture of the links between coronary artery diseases risk and lipid pathways inferred from our analysis are coherent with the state-of-the-art, while providing critical new evidences. While the association of LDL-C and TG with CAD is largely documented^39,66^, the relation linking HDL-C with CAD is more complex as both low and high HDL-C levels have been associated with a risk of cardiovascular disease and mortality^67,68^. Recent studies pointed out that functionality of HDL rather that the static measure of its circulating cholesterol level accounts for the relationship between HLD-C and CVD and mortality^68,69^, with a potential role of HDL in the remnant cholesterol transport. Overall, evidence for the presence or absence of a causal effect between lipid cholesterol measures and CAD as reported by mendelian randomization analyses should be considered with caution as lipid traits result from a complex interconnexion of multiple biological pathways. Our analysis suggests that the genetic contribution to the established negative correlation between HDL-C and CAD might be driven only by a subset of genes within a few specific genetic pathways. Under this hypothesis, drugs targeting mechanisms outside these pathways would be ineffective in decreasing CAD risk.

A number of further analyses can be conducted base on the results we obtained. First, we focused on a limited number of phenotype sets. Extending analyses to other sets of phenotypes might help refining potential genetic functional groups and better characterize theirs link to biological mechanisms. To our knowledge, there are no trivial solutions to solve the intrinsic combinatorial issue (i.e. one can build over 6×10^10^ sets of phenotypes from 36 GWAS). Also note that we worked with a data freeze dated from December 2018. Hence, at the date of the publication of this paper, newer summary statistics are available for few traits. We accounted for these new publications when counting newly identified variants by filtering associations reported in the latest version of the GWAS catalogue. Another critical component of our analysis is the methodological choices for clustering. Here we considered a Gaussian mixture model, mainly to enable missing values and used BIC and silhouette for deciding the optimal number of clusters. Other methods and alternative criteria might result in slightly different clusters. Moreover, we assume that genetic variants belong to distinct clusters, but it is likely that some variants belong to multiple biological pathways. Note that GMM provides posterior probability of cluster assignment and has the potential to explore overlapping clusters, but better approaches might potentially exist to address that specific question. Also, our implementation does not automatically address the problem of allele coding (i.e. the choice of the coded allele) inducing, in some cases, symmetric clusters which we had to merge *a posteriori*. Again, alternative approaches might offer the possibility of solving this issue.

To summarize, we ensured the theoretical reliability of a panel of multitrait tests and demonstrated their capacity to detect new associations on diverse set of traits. Considering independent significant associations, we stratified SNPs in multitrait profiles corresponding to biological pathways. We believe this stratification to be relevant for multiple applications ranging from functional annotation to drug targeting.

## Online methods

### Multivariate association test

Consider a vector **z** of *K Z*-scores statistics for a single nucleotide polymorphism (SNP) obtained from standard univariate genome-wide association screenings of *K* phenotypes. Under the null hypothesis, **z** = (*z*_1_,…, *z*_*K*,_) follows a normal distribution *N*(0, **Σ**_r_), where **Σ**_r_ is the residual phenotypic covariance matrix (**Supplementary Note**), while under the alternative, **z** is expected to display additional covariance due to shared genetics (defined by a genetic correlation matrix **Σ**_g_). We first considered an *Omnibus* test of the vector of Z-scores, which can be performed using the multivariate Wald statistics:

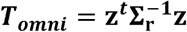

where *T*_*omni*_ follows a chi-square with *K* degree of freedom (df) under the null hypothesis of no phenotype-genotype association. We also considered a classic weight-based test which defined as:

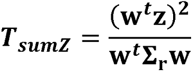

where **w** is a vector of *K* weights applied to the *Z*-score. Under the null, *T*_*sumz*_ follows a chi-squared distribution with 1 degree of freedom. Note that this approach shares similarities with both standard fixed effect meta-analysis^14^ and with dimensionality reduction methods (e.g. principal component analysis^25^). One can also note that the *Omnibus* statistics can be expressed as a combination of the *sumZ* statistics over all eigenvectors of **Σ**_*r*_ (**Supplementary Note**). We note **v**_*i*_ the i^th^ eigen vector of **Σ**_**r**_:

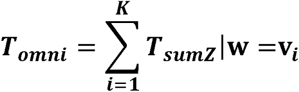

We considered four weighting schemes for the *sumZ* tests: (i) in the SumZ_1_, **w** is equal to the unit vector so all traits have the same weight; (ii) in the SumZ_r_, **w** is equal to the first eigen vector of **Σ**_r_ so its direction represents phenotypic correlation between traits, (iii) in the SumZ_g_, **w** is equal to the first eigen vector to **Σ**_g_ so its direction represents genetic correlation between traits, (iii) in the SumZ_ica_ **w** is computed by applying an Independent component analysis (ICA) to the complete matrix of Z-score. To compute the weight vector **w** of the SumZ_ica_, for a given phenotype set, the genome wide Z-score matrix was extracted and an independent component analysis was performed with the scikit-learn python package. The component yielding the most novel association was selected as loadings. We verified that this selection procedure did not lead to an inflation under the null hypothesis by simulation (see **Fig. S2**).

Performing the omnibus test requires inverting the *Z*-score covariance matrix ir. When this matrix **Σ**_r_ does not have a full rank, we use a pseudo inverse of the matrix based on the singular value decomposition (**Supplementary Note**). Briefly, as **Σ**_r_ is a variance-covariance matrix, it can be written **PDP**^**t**^ where **D** = *diag*((*λ*_*k*_)_*k*=1…*k*_), (*λ*_*k*_)_*k*=1…*k*_) are the eigenvalues of **Σ**_r_ and **P** is the orthogonal matrix whose columns correspond to the eigenvectors of **Σ**_r_. If it is not invertible, only *K’* eigenvalues are different from 0 (where *K’* denotes the rank of **Σ**_r_) and an inverse 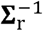 of the matrix can be computed as 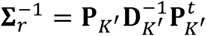, where 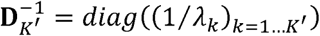 and **P**_*K’*_ denotes the *K* x *K’*, matrix whose columns are the *K’*, eigenvectors corresponding to the eigenvalues different from 0. Note that the Omnibus statistics computed with 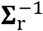 follows a *χ*^2^ distribution with *K’* degree of freedom.

### Characterization and validation of the multitrait tests

In simulation under an ideal situation, that is in the absence of missing data and knowing the true Z-score covariance matrix under the null (**Σ**_r_), the two models show correct type I error rate (**Figs S1** to **S2**). Using both simulated data and over 330K individuals and 5 quantitative traits from *UK Biobank* cohort, we next show that in the specific case of complete sample overlap between GWAS, the omnibus test is asymptotically similar to a MANOVA applied to individual level data (**Figs S4** to **S6** and **Supplementary Note**). The major potential source of bias we identified is the misspecification of **Σ**_r_ which can lead to severe type I error inflation (**Figs S7** and **S8**). Comparing various approaches, we found that **Σ**_r_ can be accurately estimated using the *LDscore* regession^9^ (**Fig. S9**), which was therefore used to estimate **Σ**_r_ along the genome-wide genetic correlation (**Σ**_g_) for the 36 phenotypes analyzed (**Tables S2** and **S3**). Nevertheless, as **Σ**_r_ depends on the sample overlap between traits, we found that even though **Σ**_r_ is correctly estimated, one can face invalid inferences for variants with statistics derived from a smaller subset of individuals than the average, a common situation in consortium studies (**Fig. S10**). To address this issue, we implemented additional tools to estimate the per SNPs sample size when missing and subsequently filter the variants with heterogeneous sample size (**Figs S11** and **S12**). Finally, another challenging issue was the merging of multiple GWAS that have missing data. Indeed, out of 10 million variants reported for some GWAS, fewer than 1,000 had complete summary statistics for all 36 phenotypes analyzed. While methods exist to impute missing GWAS statistics, they appear inaccurate for multitrait analyses and we implemented an approach we recently developed to ensure valid imputation for our context^70^ (**Fig. S13**). All pre-processing steps were also recently incorporated into a publicly available toolset^71^. After applying our pre-processing pipeline to all 36 GWAS analyzed, there remained 6,978,319 SNPs with a missing data rate of 45% (59% before imputation).

### Robust estimation of Z-score covariance

The validity of the proposed multivariate tests mostly relies on the accurate estimation of **Σ**_r_. In practice, the covariance between *Z*-scores from null SNPs from two GWAS will deviate from 0 when there is both sample overlap and correlation among the traits analyzed. When combining results from two independent studies, or when the trait analyzed has negligible correlation, **Σ**_r_ will be a diagonal matrix, so that the *Omnibus* test can be performed by summing chi-squared statistics for each SNP to form a *K* degree of freedom chi-square, and the *sumZ* test becomes a standard weighted meta-analysis of fixed effect. Yet, in the large-scale GWAS era, this situation is unlikely as most of the large GWAS are conducted in the consortium setting, where samples likely overlap across multiple GWAS. It follows that **Σ**_r_ can contain non-zero off-diagonal terms. Under the complete null model, the expected *Z*-score covariance for null SNPs between two traits equals 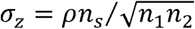 where *n*_1_ is the sample size of the first study, *n*_2_ is the sample size of the second study and *p* is the phenotypic covariance among the *n*_*s*_ overlapping samples (*see* **Supplementary Note** and e.g. ^37,38^). In some specific cases, one can obtain these parameters directly from the data (e.g. when analyzing multivariate omics data). Conversely, obtaining all four parameters (*ρ*, *n*_*s*_,*n*_1_, *n*_2_) from consortium GWAS based on dozen or even hundreds of cohorts can be a practically daunting and risky task. Moreover, accurate phenotypic covariance estimation would be particularly challenging when study-specific and trait–specific covariates adjustment has been performed. Recent studies proposed to estimate **Σ**_r_ using available SNPs from the GWAS in question using all available single SNPs *Z*-score^72^ or using a random subset of pruned variants^73^, though some discussed removing GWAS hits^15^, focusing on a subset of SNPs in regions less likely to contain causal variants^74^, or using tetrachoric estimator^16^. The validity of these estimators mostly relies on the assumption that the vast majority of the SNP effects in the genome are distributed under the null hypothesis. While this is likely to be true in some cases, associated variants can potentially lead to either upward or downward pairwise covariance between *Z*-scores. Instead, we leverage recent work by Bulik-Sullivan et al^3,9^ that allows for estimation of this covariance (and the diagonal variance terms) under a polygenic model and assuming multivariate normality of effect sizes across traits (*see* **Supplementary Note**). The estimation of **Σ**_r_ was performed on Z-scores before the imputation step described in the next section. For a few traits the estimated variance is markedly inferior to 1. As indicated in the LDSC regression method, this phenomenon happens when the original GWAS was corrected with a genomic control factor.

### Data pre-processing: an overview

The analysis of the 36 GWAS required substantial pre-processing, including the inference of several parameters. First, for many publicly available GWAS, sample size per SNP was not readily available and retrospectively collecting this information can be very challenging as it implies requesting this information from each individual cohort. For such a situation, we propose inferring a proxy for missing sample size as 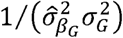, where 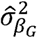 is the variance of 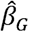, the estimated SNP effect, and 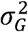 the variance of the SNP, derived from the coded allele frequency which is either provided with the GWAS or extracted from a reference panel (see **Supplementary Note**). For linear regression this approximate 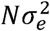, where *N* is the true sample size and 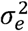 is a residual variance of the outcome in the regression model. For logistic regression our estimator is a proxy for the term *Np*(1 − *p*), where *p* is the in-sample proportion of cases, and it therefore assumes that the proportion of cases is relatively stable across SNPs with different sample size.

Another challenging issue was the merging of multiple GWAS with different set of assayed SNPs. Indeed, out of 10 million variants reported for some GWAS, fewer than 1,000 had complete summary statistics for all 36 phenotypes analyzed. We performed an imputation of missing *Z*-scores in each study using the RAISS^70^ method we recently developed. The approach uses correlation between SNPs to predict *Z*-score at missing SNPs using available ones and achieves a level of imputation accuracy suitable for multitrait analysis (**Supplementary Note**). Here we used the European panels from the 1,000 Genomes project^75^ as a reference for the estimation of the correlation between SNPs. Overall, imputation did not lead to any observable inflation of the *omnibus* statistic (**Fig. S13**). Nevertheless, as a supplementary quality control (QC), we excluded significant SNPs that were not surrounded by SNPs in linkage disequilibrium with significant or near significant *p*-values (*P* < 10^−6^).

These two parameter inferences were integrated along other pre-processing operations into a pipeline that is fully described here^76^. Given a reference panel with no ambiguous strand, it consists in the following steps (i) Extract, the coded and alternative alleles, signed statistics (regression coefficient or odds ratio), standard error, p-value, and sample size; (ii) Remove all SNPs that are not in the reference panel; (iii) Derive *Z*-score for each SNP from signed statistics and *p*-value; (iv) Infer sample size when not available; (v) Remove all SNPs whose sample size is less than 70% of the maximum sample size; and (vi) Infer missing *Z*-scores statistics based on the 1K genome reference panel. After applying our pre-processing pipeline to all 36 GWAS analyzed, there remained 6,978,319 SNPs with a missing rate of 45% (59% before imputation).

### Characterization of new loci

To determine new and existing trait-associated loci we used genome regions formed by linkage disequilibrium (LD) blocks as defined in Berisa et al^77^ using a reference panel of individuals of European ancestry. It included a total of 1,704 independent regions ranging from 10 kb to 26 Mb in length, with an average size of 1.6 Mb. For each independent LD region, we extracted the minimum *p*-value over all SNPs contained in the region, and a single univariate analysis *p*-value defined as the minimum across all single phenotype GWAS and all SNPs in the region. We consider that a region is newly detected by a multitrait test if the joint analysis *p*-value is genome-wide significant while its univariate *p*-value is not (joint analysis *p*-value < 1×10^−8^ and univariate *p*-value > 1x 10^−8^). We determined SNPs carrying the signal inside significant region with the plink “clump” function using the following parameters: --clump-p1, 10^−8^; --clump-r2, 0.2. We kept the lead SNP by clump for further analysis (gene mapping and clustering).

To report associations exclusively detected in the current report (**Table S4** to **S10**), we filtered out association present in the GWAS catalogue^1^ at the date of the 14^th^ of September 2020 (univariate *p*-value > 5x 10^−8^) for traits corresponding to our phenotype set. The following trait labels were used to retrieve associations: (Metabolism set) ‘Fasting blood glucose’, ’Triglycerides’, ‘LDL cholesterol’, ‘LDL cholesterol levels’, ‘HDL cholesterol’, ‘HDL cholesterol levels’, ‘Total cholesterol levels’, ‘HOMA-B’, ‘HOMA-IR’, ‘Hemoglobin A1c levels’, ‘Type 2 diabetes’; (Psychiatric set) ‘Schizophrenia’, ‘Bipolar disorder’, ‘Major depressive disorder’, ‘Alzheimer’s disease’, ‘Educational attainment’; (Anthropometry set) ‘Height’, ‘Waist circumference’, ‘Waist-hip ratio’, ‘Body mass index’, ‘Hip circumference’; (*Immunity* set) ‘Bone mineral density’, ‘Rheumatoid arthritis’, ‘Ulcerative colitis’, ‘Inflammatory bowel disease’, ‘Crohn’s disease’, ‘Asthma’; (Cardiovascular set) ‘Coronary artery disease’, ‘Ischemic stroke’, ‘Large artery stroke’, ‘Stroke’, ‘Atrial fibrillation’, ‘Heart rate’, ‘Heart rate variability traits’; (Composite set) ‘Body mass index’, ‘Waist-hip ratio’, ‘Triglycerides’, ‘LDL cholesterol’, ‘LDL cholesterol levels’, ‘HDL cholesterol’, ‘HDL cholesterol levels’, ‘Total cholesterol levels’.

### FUN-LDA tissue enrichment

We computed enrichment for SNPs belonging to regions of open chromatin (more likely to contain expressed genes^78,79^) in specific tissues in three cases: i) when comparing results across phenotype sets, ii) when comparing univariate results, and iii) when comparing results across clusters. For all analyses we used functional annotations on 127 Roadmap tissues and cell lines defined by integrating activating histone marks (H3K4me1, H3K4me3, H3K9ac, and H3K27ac) with a latent Dirichlet allocation model as implemented in FUN-LDA^80^. The enrichment score for a tissue is based on the number significant SNPs compared with the total number of SNPs in open chromatin region (see **Supplementary Note**). Enrichment results are reported in **Tables S11** to **S13**.

### Multitrait genetic association clustering and selection of the optimal number of clusters

We performed a clustering of top associated SNPs for each phenotype set using a Gaussian Mixture model (GMM). One major difficulty in applying the GMM was to deal with incomplete data. Indeed, even after imputation of some missing statistics, our datasets still contained some missing values. To solve the clustering, we implemented the statistical framework described by Ghahramani et al^81^ which we recently implemented in a R package MGMM^82^. The model gives for each SNP the posterior probabilities to belong to each cluster, and was therefore assigned to its most likely cluster, as long as its entropy was larger than 0.75. For a given variant *SNP*_*i*_, the entropy was derived as follow:

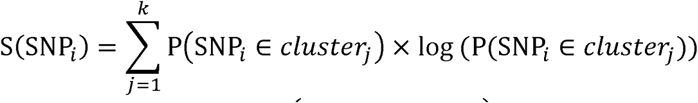

where *k* is the total number of clusters and P(SNP_*i*_ ∈ *cluster*_*j*_) is the posterior probability of SNP_*i*_ to belong to cluster *j*. The higher the entropy the more the SNP attribution to one cluster is ambiguous. SNPs with an entropy higher than 0.75 were filtered out of the clustering results.

Clustering was performed on all independent significant SNPs. For each SNP, we defined three *p*-values on the phenotypic group traits: the minimum univariate *p*-value (P_univ_), the *SumZ*_*ica*_ *p*-value and the *omnibus p*-value. All SNPs with at least one of the three *p*-value under 10^−8^ were selected for further analysis. For the *Metabolism* univariate clustering, we only considered the univariate *p*-value to perform the selection. We then applied the plink^83^ clump function to retrieve practically independent associations using the 0.2 as clump-r2 parameter and 10^−8^ as clump-p1 parameter. For each clump we selected a representative SNPs as the one with the smallest *p*-value across the three tests and having more than 60% of its values observed. Note that for a negligible number of occurrences, the representative SNPs has a *p*-value above 10^−8^ (**Table S15** and **Table S16**). We applied MGMM within each phenotype set and varied the pre-specified number of clusters between 2 and 10. To select the optimal number of clusters *k*, we performed the clustering 100 times on a random subset of 80% of the SNPs for each k. For each resulting clustering we computed the Bayesian Information Criteria and the Silhouette^84^ (see **Figs S22** and **S23**). Except for the *Metabolism* set, the silhouette appears conservative and the BIC criterion anticonservative, i.e. the latter criteria tends to select a larger number of clusters. We decided to use the following ad hoc compounded criterion:

1. If the optimal number of clusters determined by the BIC criteria is higher than the one determined by the silhouette criteria, starting from the silhouette optimal, increase the number of clusters until one of these two conditions is met: 1) adding one cluster significantly decrease the silhouette criterion, 2) the BIC optimal number is reached.
2. In other cases, set the optimal number of clusters to the one determined by silhouette.

### Cluster genetic correlation

We defined pairwise genetic covariance per cluster for as 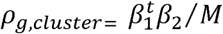 where *β*_1_ and *α*_2_ are the vector of genetic effects for the pair of phenotypes considered and *M* is the number of SNPs in the cluster. To estimate properly this quantity from the observed 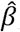, we accounted for the bias introduced by sample overlap and phenotypic correlation using the following estimator (see **supplementary notes**):

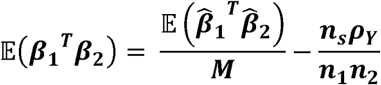

where *ρ*_*Y*_ is the phenotypic covariance, and *n*_*s*_, *n*_1_ and *n*_2_ are respectively the sample size shared between the two traits, for the trait 1, and for the trait 2. To assess whether the estimated genetic covariances are significantly different from zero, we performed for each pair of phenotypes within each cluster, a *t*-test on the vector of random variables (*X*_1_,*X*_2_,…,*X*_*M*_), were 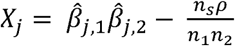 is the contribution of SNP *j* to the covariance. Note that we used only independent SNPs selected using LD-clumping with squared-correlation parameter equals 0.2.

### Functional enrichment of metabolism clusters

We used FUMA^85^ *SNP2GENE* function to associate SNPs with genes based on two criteria, the physical position (in 30kb radius of a protein coding gene) and eQTLs (all significant cis-eQTL from GTEx up to a distance of 1Mb). Note that we restrained the eQTLs to the one that were found in relevant tissue for the Immunity and Metabolism set: immune cells for Immunity and adipose, intestine, liver and brain tissues for Metabolism (see **Supplementary Data** 1 for complete parameters). After chaining genes to clusters based on SNPs, we performed a functional enrichment for pathways defined in KEGG^86^ and GO^87^ databases and derived report *p*-values using FUMA *GENE2FUNC* function. Here, cluster’s gene were compared against a background of protein coding genes. Finally, we used the R package pathview^88^ to project genes onto KEGG pathways maps.

### Disease-clusters association

For the *metabolism* phenotype set, to provide an indicator of the relative contribution of genetic variants to phenotypes in each cluster from the *Metabolism* set, we performed a principal component analysis (PCA) of the SNP-by-phenotype association matrix within each cluster. For this analysis, we used scaled beta coefficients, i.e. *Z*-scores divided by the square root of the phenotype GWAS sample size. To avoid bias in due to the arbitrary choice of the coded allele, we randomly shuffled 20 times the coded allele, and repeated the PCA after each shuffling. We report in Fig. 5, the average of the loadings of the first PC over all shuffling. Note that the first PC only provides the multidimensional direction explaining the largest variance and therefore do not fully capture the distribution of genetic effect within each cluster. Nevertheless, those first PCs explained a substantial amount of the total variance, equal to 75%, 38%, 53%, 64%, 80% and 93% of the variance in betas for cluster 1 to 6, respectively.

Then, we assessed the association between SNPs within the inferred cluster and three traits (none of which being included in the *Metabolism* set): cardiovascular diseases, any stokes and BMI. SNP alleles were aligned according to the first principal by clusters determined in the last section. We applied a sign test to assess the concordance of the sign of the projection on PC1 and the sign of Z-score for on the three tested additional traits. For this analysis we used more stringent criteria to ensure the SNPs independence. We selected the subset of *Metabolism* SNPs for which linkage disequilibrium does not exceed 0.2 (clump-r2 set to 0.05), which diminishes the number of SNPs considered from 391 to 285. Concerning the association of SNPs to drug target, we associated drug target to a representative SNPs by selecting the SNP with the lowest entropy and having a positive silhouette.

## Acknowledgements

This work has been conducted as part of the INCEPTION program (ANR-16-CONV-0005). It was also supported by NIH grant R03DE025665 to H.A.

## Author contributions

**Conceptualization**: HA, BW; **Formal analysis**: HJ, HA, BW, VL, ZH, CL, AZ, AV; **Software**: HJ, VG, PL, HM, ZRM; **Supervision**: HA; **Original draft preparation**: HJ, HA; **Review and editing**: VL, ZRM, WL, MPD, PK, IIL, BW

## Data and materials availability

All GWAS summary statistics data used in this study are publicly available. Links to each dataset are provided in Table S1. All other derived data are available in the main text or the supplementary materials.

## URL Resources

JASS_Preprocessing: https://gitlab.pasteur.fr/statistical-genetics/jass_preprocessing

JASS: https://gitlab.pasteur.fr/statistical-genetics/jass

RAISS: https://gitlab.pasteur.fr/statistical-genetics/raiss

MGMM: https://github.com/zrmacc/MNMix

## Supplementary Materials

Materials and Methods

Supplementary Text

Figures S1-S32

Tables S1-S19

External Dataset S1

