## Supplementary materials for "Multitrait genetic-phenotype associations to connect disease variants and biological mechanisms"

### Supplementary Note

##### *Multivariate statistical test*

The *Omnibus* test is a standard generalization of Student's t-statistic to the multivariate case. It aims at testing the null hypothesis of no association between a single SNP and none of the outcomes. Briefly, assuming a vector of z-scores $\mathbf{z}$ of size $K$ follows a multivariate normal with mean 0 and covariance $\boldsymbol{\Sigma}_\mathbf{r}$, the probability that an occurrence $\mathbf{z}$ belong to the multivariate distribution can be estimated by comparing $T_{omni}$, the square of its *Mahalanobis* distance (a generalization of the mean to standard deviation distance), to a chi-squared distribution with $K$ degree of freedom (although note that for the special case of small sample size relative to the number of outcomes, alternative distribution should be used, see the section about MANOVA). In practice, $T_{omni}$ is defined as:

$$\boldsymbol{T_{omni}} = \mathbf{z}^t \boldsymbol{\Sigma}_\mathbf{r}^{-1} \mathbf{z} \sim \chi^2_{Kdf}$$

The *sumZ* statistics consists in testing whether a specific linear combination of the individual univariate statistics follows a null distribution between the tested SNP and the outcomes. We defined a simple weighted sum of z-score, $S = \mathbf{w}^t\mathbf{z}$. Again, assuming $\mathbf{z} \sim \mathbf{MVN}(\mathbf{0}, \boldsymbol{\Sigma}_\mathbf{r})$, where $\mathbf{MVN}$ is the multivariate normal distribution, the statistics $S$ being a sum of normally distributed variables, also follows a normal distribution with mean and variance equal to:

$$\mathbb{E}[\mathbf{w}^t\mathbf{z}] = \mathbb{E}\left[\sum_{i=1}^{K} w_i z_i\right] = \sum_{i=1}^{K} w_i \mathbb{E}[z_i] = \mathbf{0}$$

$$var[\mathbf{w}^t\mathbf{z}] = var\left[\sum_{i=1}^{K} w_i z_i\right] = \sum_{i=1}^{K}\sum_{j=1}^{K} w_i w_j var[z_i z_j] = \mathbf{w}^t \boldsymbol{\Sigma}_\mathbf{r} \mathbf{w}$$

Test of association can thus be defined as a standard two-sided *t*-test, i.e.:

$$t_{sumZ} = \frac{\mathbf{w}^t\mathbf{z} - \mathbb{E}[\mathbf{w}^t\mathbf{z}]}{\sqrt{var[\mathbf{w}^t\mathbf{z}]}} = \frac{\mathbf{w}^t\mathbf{z}}{\sqrt{\mathbf{w}^t\boldsymbol{\Sigma}_\mathbf{r}\mathbf{w}}}$$

Or more commonly, to a 1 degree of freedom chi-squared test statistics:

$$T_{sumZ} = \frac{(\mathbf{w}^t\mathbf{z})^2}{\mathbf{w}^t\boldsymbol{\Sigma}_\mathbf{r}\mathbf{w}} \sim \chi^2_{1df}$$

**Figure S1** shows the correct calibration of the two statistics under null model.

##### *Relationship between the proposed statistical tests*

The two proposed statistics can be linked in the special case where the weights $\mathbf{w}$ equal $\mathbf{v}$, the eigenvector of $\boldsymbol{\Sigma}_\mathbf{r}$:

$$T_{omni} = \sum_{i=1}^{K} T_{sumZ} | \mathbf{w} = \mathbf{v}_i$$

<u>*Proof:*</u> Consider the $K \times K$ variance-covariance matrix $\boldsymbol{\Sigma}_\mathbf{r}$ of the z-scores. As $\boldsymbol{\Sigma}_\mathbf{r}$ is symmetric and using the spectral theorem, there exists an orthogonal matrix $\mathbf{P}$ and a diagonal matrix $\mathbf{D}$ such that:

$$\boldsymbol{\Sigma}_\mathbf{r} = \mathbf{P}\mathbf{D}\mathbf{P}^\mathbf{t}$$

where the columns of $\mathbf{P}$ correspond to the eigenvectors $\mathbf{v}_{i=1\ldots K}$ of $\boldsymbol{\Sigma}_\mathbf{r}$ and the diagonal of $\mathbf{D}$ corresponds to the eigenvalues $\lambda_{i=1\ldots K}$ of $\boldsymbol{\Sigma}_\mathbf{r}$. The covariance matrix can then be further written as:

$$\boldsymbol{\Sigma}_\mathbf{r} = \sum_{i=1}^{K} \lambda_i \mathbf{v}_i \mathbf{v}_i^t$$

It follows that the *Omnibus* test can be rewritten as:

$$
\begin{aligned}
\boldsymbol{T_{omni}} &= \boldsymbol{z^t \Sigma_r^{-1} z} \\
&= \boldsymbol{z^t (PDP^t)^{-1} z} \\
&= \boldsymbol{z^t (P^t)^{-1} D^{-1} P^{-1} z} \\
&= \boldsymbol{z^t P D^{-1} P^t z} \\
&= \mathbf{z}^t \left( \sum_{i=1}^{K} \frac{\mathbf{1}}{\boldsymbol{\lambda_i}} \mathbf{v}_i \mathbf{v}_i^t \right) \mathbf{z} \\
&= \sum_{i=1}^{K} \frac{\mathbf{1}}{\boldsymbol{\lambda_i}} (\mathbf{z}_i^t \mathbf{v}_i)(\mathbf{z}_i^t \mathbf{v}_i)^t \\
&= \sum_{i=1}^{K} \frac{\mathbf{1}}{\boldsymbol{\lambda_i}} (\mathbf{z}_i^t \mathbf{v}_i)^2
\end{aligned}
$$

On the other hand, applying the *sumZ* statistic with weights $\mathbf{w} = \mathbf{v}_i$, where $\mathbf{v}_i$ is the eigenvector $i$ of $\mathbf{\Sigma_r}$ corresponding to $\lambda_i$,we have:

$$\boldsymbol{T_{sumZ}} = \frac{(\mathbf{v}_i^t \mathbf{z})^2}{\mathbf{v}_i^t \mathbf{\Sigma_r v_i}}$$

As $\mathbf{v}_i$ is a principal component of $\mathbf{\Sigma_r}$, by definition we have $\mathbf{v}_i^t \mathbf{\Sigma_r} \mathbf{v}_i = \lambda_i \mathbf{v}_\mathrm{i}^t \mathbf{v}_i$, but also $\mathbf{v}_i^t \mathbf{v}_i = 1$. It follows that:

$$\boldsymbol{T_{sumZ}} = \frac{\mathbf{1}}{\boldsymbol{\lambda_i}} (\mathbf{v}_i^t \mathbf{z})^2$$

##### *Solving singular covariance matrix*

Consider the case where the estimated variance-covariance matrix $\widehat{\mathbf{\Sigma}}_\mathbf{r}$ is not invertible. We compared the performances of three methods described below to derive a pseudo-inverse of $\widehat{\mathbf{\Sigma}}_\mathbf{r}$. Using the spectral theorem, $\widehat{\mathbf{\Sigma}}_\mathbf{r}$ can be written:

$$\widehat{\mathbf{\Sigma}}_\mathbf{r} = \mathbf{PDP}^{\boldsymbol{t}}$$

where $\mathbf{D} = diag((\lambda_k)_{k=1 \ldots K})$, $(\lambda_k)_{k=1 \ldots K}$ are the eigenvalues of $\widehat{\mathbf{\Sigma}}_\mathbf{r}$ and $\mathbf{P}$ is the orthogonal matrix which columns correspond to the eigenvectors of $\widehat{\mathbf{\Sigma}}_\mathbf{r}$.

***<u>Strategy 1:</u>*** The first strategy consists in considering only eigenvalues greater than a specified threshold $\epsilon$. Suppose only $K'$ eigenvalues out of $K$ are greater than $\epsilon$. Let $D_{K'} = diag((\lambda_k)_{k=1 \ldots K'})$ and $\mathbf{P}_{K'}$ denote the $K \times K'$ matrix which columns are the $K'$ eigenvectors corresponding to the eigenvalues greater than $\epsilon$. We then define the pseudo-inverse of $\widehat{\mathbf{\Sigma}}_\mathbf{r}$ by:

$$\widehat{\mathbf{\Sigma}}_\mathbf{r}^* = \mathbf{P}_{\boldsymbol{K'}} \mathbf{D}_{\boldsymbol{K'}}^{-1} \mathbf{P}_{\boldsymbol{K'}}^{\boldsymbol{t}}$$

***<u>Strategy 2:</u>*** The second strategy is relatively similar to the first one. Instead of considering only the eigenvalues greater than a threshold $\epsilon$, eigenvalues below this threshold are set to $\epsilon$. Let $\mathbf{D}_\epsilon = diag((\lambda'_k)_{k=1 \ldots K'})$ where $\lambda'_k$ is equal to $\lambda_k$ if $\lambda_k$ is greater than $\epsilon$ and $\epsilon$ otherwise. We then define the pseudo-inverse of $\widehat{\mathbf{\Sigma}}_\mathbf{r}$ by:

$$\widehat{\mathbf{\Sigma}}_\mathbf{r}^* = \mathbf{P}\mathbf{D}_{\boldsymbol{\epsilon}}^{-1}\mathbf{P}^{\boldsymbol{t}}$$

***<u>Strategy 3:</u>*** The third strategy consists in adding a small value $\epsilon$ to the diagonal terms of $\widehat{\mathbf{\Sigma}}_\mathbf{r}$. The pseudo-inverse of $\widehat{\mathbf{\Sigma}}_\mathbf{r}$ is defined by:

$$\widehat{\mathbf{\Sigma}}_\mathbf{r}^* = \left(\widehat{\mathbf{\Sigma}}_\mathbf{r} + \boldsymbol{\epsilon} \times \mathbf{I}_{\boldsymbol{K}}\right)^{-\mathbf{1}}$$

where $\mathbf{I}_K$ denote the $K \times K$ identity matrix.

We conducted simulation series to compare the relative performances of the three strategies for three different thresholds ($\epsilon = 1 \times 10^{-3}, 1 \times 10^{-6}, 1 \times 10^{-9}$). As showed in **Figure S1,** the distributions of the *p*-values were correctly calibrated for all three tests for the two highest thresholds, but remain correct only for strategy 1 for the lowest $\epsilon$. The latter strategy was therefore used in all analyses, as implemented in the JASS python package[1].

##### *Theoretical comparison with MANOVA*

In the special case of complete sample overlap and no missing phenotypic value, the proposed *Omnibus* statistics is asymptotically similar to the one-way MANOVA. Consider $K$ correlated traits $Y_k$, $k = 1, \dots, K$ and let $\mathbf{Y}$ denote the $n \times K$ matrix of traits for all $n$ individuals. For a given variant, let $\mathbf{x}$ be $n \times 1$ vector of predictors. The underlying model in the MANOVA model can be written as:

$$\mathbf{Y} = \mathbf{x}\boldsymbol{\beta}^{\mathbf{t}} + \boldsymbol{\varepsilon}$$

where $\boldsymbol{\beta}$ is the $1 \times K$ vector of genetic effect on the $K$ phenotypes and $\varepsilon$ is the matrix of errors. As for the proposed *Omnibus* test, the null hypothesis tested in the MANOVA is $\boldsymbol{\beta} = 0$, and it does not rely on an assumption about the direction of the effects. The Wilk's Lambda test statistic is defined as follows:

$$\boldsymbol{W} = \frac{\boldsymbol{det}(\mathrm{E})}{\boldsymbol{det}(\mathrm{H} + \mathrm{E})} = \frac{\boldsymbol{det}\left(\boldsymbol{Y^t Y} - \widehat{\boldsymbol{\beta}}(\mathbf{x^t x})\widehat{\boldsymbol{\beta}}^{\boldsymbol{t}}\right)}{\boldsymbol{det}(\mathbf{Y^t Y})}$$

where $\mathbf{E} = \mathbf{Y}^t\mathbf{Y} - \widehat{\boldsymbol{\beta}}(\mathbf{x}^t\mathbf{x})\widehat{\boldsymbol{\beta}}^t$, $\mathbf{H} = \widehat{\boldsymbol{\beta}}(\mathbf{x}^t\mathbf{x})\widehat{\boldsymbol{\beta}}^t$ and $\widehat{\boldsymbol{\beta}} = \mathbf{Y}^t\mathbf{x}(\mathbf{x}^T\mathbf{x})^{-1}$. Under the null, $W$ follows a Wilks' lambda distribution $\Lambda(K, n-1, 1)$. In practice, the statistical test is commonly performed using a *Fisher* distribution through the approximation $\frac{1-\Lambda(K,n-1,1)}{\Lambda(K,n-1,1)} \sim \frac{K}{n-K} F_{K,n-K}$ where $F_{p,q}$ denotes the Fisher distribution with $p$ and $q$ degrees of freedom. Furthermore, note that when $n$ is large compared to $K$, the quantity $\left(\frac{K}{2} - n + 1\right)\log(W)$ can be approximated by a chi-square distribution with $K$ degrees of freedom under the null. More details on *Wilks'* lambda distribution and its approximation can be found in Mardia et al.[2] and Bartlett[3]. Importantly, the Fisher approximation of the *Wilks'* statistics can be directly derived using only summary statistics since one can use the phenotypic variance-covariance matrix derived using the LD Score to estimate the matrix $\mathbf{Y}^t\mathbf{Y}$, while $\widehat{\boldsymbol{\beta}}(\mathbf{x}^t\mathbf{x})\widehat{\boldsymbol{\beta}}^t$ can be estimated by $\mathbf{z}^t\mathbf{z}$. Nevertheless, although the null hypothesis tested in the MANOVA and the *Omnibus* test is the same, a direct theoretical comparison of the two approaches is non-trivial. Instead, we performed series of simulation studies (**Figures S4** and **S5**) to assess the relative performances of these approaches.

Overall, all analyses confirm the three tests show similar results as long as $K \gg n$ as it is the case in large consortia and most of recent GWAS. In the special case where $n$ is relatively close to $K$ (see e.g. **Figures S4c** and **S5c,** where $K$=100 and $N$=500), the *Omnibus* test shows deflated statistics and therefore, decreased power as compared to the MANOVA. For such settings, we recommend using the Fisher approximation, which shows improved adequacy with the MANOVA. Finally, for the special case when $n$ is relatively close to $K$, but outcomes have only partial sample overlap, there are no trivial solution and further development is therefore required.

##### *Validation of the approach using UK Biobank data*

We used data from the UK Biobank cohort[4,5] to validate our approach and our theoretical comparison between the *Omnibus* approach and the MANOVA. In brief, we selected five anthropometric traits and 619,017 high-quality genotyped SNPs with minor allele frequencies (MAF) > 1% available in a subset of 336,347 unrelated individuals of British ancestry. For ease of comparison, we projected out all relevant covariates and used the residual as primary outcomes in all individual-level data analyses.

As showed in **Figure S7** and **S8**, a correct estimation of the covariance matrix between GWAS summary statistics under the null is critical and can be impacted by true genetic effects. We first aimed at confirming the validity of the LDSC estimator. We performed multiple analyses while varying the sample size use to generate summary statistics by randomly sub-sampling the complete data. In a first analysis, we down-sampled the whole dataset by removing 50%, 90% and 99% of all individuals; the resulted sample size was 168,173, 33,635 and 3,363 individuals, respectively. In a second analysis, we down-sampled the individuals but for a single trait at the same rates (i.e. removing 50%, 90% and 99% of individuals), thus inducing different sample overlaps between the down-sampled trait and other traits. In all scenarios considered, we observed a very strong concordance between the LDSC estimates and the expected value derived using the estimated phenotypic correlation and the true sample size overlap (**Figure S9**), demonstrating the adequacy of the LDSC regression models, at least for the anthropometric traits used in our analysis which are known to show high polygenicity[6].

For the comparison with the MANOVA, we ran its implementation in PLINK[7]. For the *Omnibus* test, we proceeded in three steps: (1) compute single-trait GWAS for each of the five anthropometric traits and store the summary statistics; (2) estimate the phenotypic correlation matrix by LDSC[8]; and (3) computed the *Omnibus* test to get multi-trait statistics. Looking at differences between -log10(*p*-values) from MANOVA and the *Omnibus* for a subset of 14,718 genotyped SNPs on Chromosome 20, we observed almost perfect correlation between the two approaches (**Figure S6**). Note that once the individual phenotype summary statistics are derived, the computation time of MANOVA was considerably larger than for the *Omnibus* test (approximately 50 min *vs.* 5 min).

##### *Expected z-score covariance for null SNPs*

Consider two standardized quantitative phenotypes $\mathbf{y}_1$ and $\mathbf{y}_2$ that have been tested for association with a set of $m$ single nucleotide polymorphisms (SNP) in studies with sample size $n_1$ and $n_2$, respectively, and $n_s$ samples shared between the two studies. We aim at estimating the expected covariance between the resulting z-score statistics for a given null predictor –i.e. for a putative variant drawn independently from others SNPs and not associated with the outcomes under study. We denote $\mathbf{x}_1$ and $\mathbf{x}_2$ the column vectors of standardized predictors for such a null variants in dataset 1 and 2. We can write $\mathbb{E}[z_1 z_2]$, the expected covariance between the z-scores at for $\mathbf{x}_1$ and $\mathbf{x}_2$, as:

$$\begin{aligned}\mathbb{E}[\boldsymbol{z_1 z_2}] &= \mathbb{E}\left[\mathbb{E}\left[\frac{\left((\mathbf{x}_1^t\mathbf{x}_1)^{-1}\mathbf{x}_1^t\mathbf{y}_1\right)}{\sqrt{(\mathbf{x}_1^t\mathbf{x}_1)^{-1}}}\left(\frac{\left((\mathbf{x}_2^t\mathbf{x}_2)^{-1}\mathbf{x}_2^t\mathbf{y}_2\right)}{\sqrt{(\mathbf{x}_2^t\mathbf{x}_2)^{-1}}}\right)^{\boldsymbol{t}} |\mathbf{x}_1,\mathbf{x}_2,\mathbf{y}_1,\mathbf{y}_2\right]\right]\\ &= \mathbb{E}\left[\mathbb{E}\left[\frac{\mathbf{x}_1^t\mathbf{y}_1(\mathbf{x}_2^t\mathbf{y}_2)^{\boldsymbol{t}}}{\sqrt{\boldsymbol{n_1}}\sqrt{\boldsymbol{n_2}}}|\mathbf{x}_1,\mathbf{x}_2,\mathbf{y}_1,\mathbf{y}_2\right]\right]\\ &= \frac{\mathbf{1}}{\sqrt{\boldsymbol{n_1 n_2}}}\mathbb{E}[\mathbf{x}_1^t\mathbf{y}_1\mathbf{y}_2^t\mathbf{x}_2]\\ &= \frac{\boldsymbol{\rho n_s}}{\sqrt{\boldsymbol{n_1 n_2}}}\end{aligned}$$

where $\rho$ is the phenotypic covariance derived over the $n_s$ overlapping samples.

Note that this is in agreement with recent work on the LD-score regression by Bulik-Sullivan et al[8]. Indeed, consider further that $\mathbf{y}_1 = \mathbf{G}_1\boldsymbol{\beta}_1 + \boldsymbol{\varepsilon}_1$, and $\mathbf{y}_2 = \mathbf{G}_2\boldsymbol{\beta}_2 + \boldsymbol{\varepsilon}_2$, where $\mathbf{G}_1$ and $\mathbf{G}_2$ are $n_1 \times m$ and $n_2 \times m$ matrices of standardized genotypes for the causal variants of outcome 1 and outcome 2, respectively, and $\boldsymbol{\varepsilon}_1$ and $\boldsymbol{\varepsilon}_2$ are there residual variance. Now assume the effects of $\mathbf{G}_1$ and $\mathbf{G}_2$ are distributed as $\boldsymbol{\beta}_1 \sim \mathcal{N}(0, h_1^2/m)$ and $\boldsymbol{\beta}_2 \sim \mathcal{N}(0, h_2^2/m)$, where $\boldsymbol{\beta}_1$ and $\boldsymbol{\beta}_2$ are two column vectors of size $m$ with $cov(\boldsymbol{\beta}_1, \boldsymbol{\beta}_2) = \sigma_g = r_g\sqrt{h_1^2 h_2^2}/m$, and that $\mathbf{y}_1$ and $\mathbf{y}_2$ also shared non-genetic variance, equals to $\sigma_e$. We can further re-write $\mathbb{E}[z_1 z_2]$ as:

$$\mathbb{E}[\boldsymbol{z_1 z_2}] = \frac{\mathbf{1}}{\sqrt{\boldsymbol{n_1 n_2}}}\mathbb{E}[\mathbf{x}_1^t\mathbb{E}[(\mathbf{G}_1\boldsymbol{\beta}_1 + \boldsymbol{\varepsilon}_1)(\mathbf{G}_2\boldsymbol{\beta}_2 + \boldsymbol{\varepsilon}_2)^{\boldsymbol{t}}|\mathbf{x}_1,\mathbf{x}_2,\mathbf{G}_1,\mathbf{G}_2]\mathbf{x}_2]$$

$$= \frac{\mathbf{1}}{\sqrt{\boldsymbol{n_1 n_2}}} \mathbb{E}[\mathbf{x}_1^t \mathbb{E}[(\mathbf{G_1}\boldsymbol{\beta_1} + \boldsymbol{\varepsilon_1})(\boldsymbol{\beta}_2^t \mathbf{G}_2^t + \boldsymbol{\varepsilon}_2^t)|\mathbf{x_1}, \mathbf{x_2}, \mathbf{G_1}, \mathbf{G_2}]\mathbf{x_2}]$$
$$= \frac{\mathbf{1}}{\sqrt{\boldsymbol{n_1 n_2}}} \mathbb{E}[\mathbf{x}_1^t (\mathbf{G_1}\mathbb{E}[(\boldsymbol{\beta_1}\boldsymbol{\beta}_2^t)]\mathbf{G}_2^t + \mathbb{E}[\boldsymbol{\varepsilon_1}\boldsymbol{\beta}_2^t \boldsymbol{G}_2^t] + \mathbb{E}[\mathbf{G_1}\boldsymbol{\beta_1}\boldsymbol{\varepsilon}_2^t] + \mathbb{E}[\boldsymbol{\varepsilon_1}\boldsymbol{\varepsilon}_2^t])\mathbf{x_2}]$$
$$= \frac{\mathbf{1}}{\sqrt{\boldsymbol{n_1 n_2}}} \mathbb{E}[\mathbf{x}_1^t \mathbf{G_1}\mathbb{E}[(\boldsymbol{\beta_1}\boldsymbol{\beta}_2^t)]\mathbf{G}_2^t \mathbf{x_2}] + \frac{\mathbf{1}}{\sqrt{\boldsymbol{n_1 n_2}}} \mathbb{E}[\mathbf{x}_1^t \mathbb{E}[\boldsymbol{\varepsilon_1}\boldsymbol{\varepsilon}_2^t]\mathbf{x_2}]$$
$$= \frac{\mathbf{1}}{\sqrt{\boldsymbol{n_1 n_2}}} \mathbb{E}\left[\mathbf{x}_1^t \mathbf{G_1}\frac{\boldsymbol{\sigma_g}}{\boldsymbol{m}}\mathbf{I}_m \mathbf{G}_2^t \mathbf{x_2}\right] + \frac{\mathbf{1}}{\sqrt{\boldsymbol{n_1 n_2}}} \boldsymbol{\sigma_e n_s}$$
$$= \frac{\mathbf{1}}{\sqrt{\boldsymbol{n_1 n_2}}} \frac{\boldsymbol{\sigma_g}}{\boldsymbol{m}} \mathbb{E}[\mathbf{x}_1^t \mathbf{G_1}\mathbf{G}_2^t \mathbf{x_2}] + \frac{\boldsymbol{\sigma_e n_s}}{\sqrt{\boldsymbol{n_1 n_2}}}$$

Furthermore, we have $\mathbb{E}\left[\mathbf{x}_{1i}^t \mathbf{G}_1 \mathbf{G}_2^t \mathbf{x}_2\right] = \mathbb{E}[(\mathbf{r}n_1 + \boldsymbol{\theta}_1)^t(\mathbf{r}n_2 + \boldsymbol{\theta}_2)]$, where $\mathbf{r}$ is a vector of the true correlation between the $\mathbf{x}_.$ and all $m$ SNPs from $\mathbf{G}_.$, and $\boldsymbol{\theta}_1$ and $\boldsymbol{\theta}_2$ are the residuals noise due to finite sample size. Now, as compared to the derivation from Bulik-Sullivan et al[8], we consider here the special case where $\mathbf{x}_1$ and $\mathbf{x}_2$ are not correlated with any causal (i.e. independent of $\mathbf{G}_1$ and $\mathbf{G}_2$, respectively), so that $\mathbf{r} = \mathbf{0}$ and $\mathbb{E}\left[\mathbf{x}_{1i}^t \mathbf{G}_1 \mathbf{G}_2^t \mathbf{x}_2\right] = \mathbb{E}[\boldsymbol{\theta}_1^t \boldsymbol{\theta}_2]$. In the special case of no sample overlap, $\boldsymbol{\theta}_1$ and $\boldsymbol{\theta}_2$ are independent, so that the latter expectation is null. However, when there is sample overlap, subsets of $\mathbf{x}_1$ and $\mathbf{x}_2$, as well as $\mathbf{G}_1$ and $\mathbf{G}_2$, will be identical. We refer to these group as $\mathbf{x}_s$ and $\mathbf{G}_s$, respectively, and to $\mathbf{x}_1^*$, $\mathbf{x}_2^*$, $\mathbf{G}_1^*$,and $\mathbf{G}_2^*$, for their respective complements. We have now:

$$\mathbb{E}[\boldsymbol{\theta}_1^t \boldsymbol{\theta}_2] = \mathbb{E}\left[\left(\sum\nolimits_{i\in n_s} \mathbf{x}_{si}^t \mathbf{G}_{si} + \sum\nolimits_{i\notin n_s} \mathbf{x}_{1i}^{*t} \mathbf{G}_{1i}^*\right)^t \left(\sum\nolimits_{j\in n_s} \mathbf{x}_{sj}^t \mathbf{G}_{sj} + \sum\nolimits_{j\notin n_s} \mathbf{x}_{2j}^{*t} \mathbf{G}_{2j}^*\right)\right]$$
$$= \mathbb{E}\left[\left(\sum\nolimits_{i\in n_s} \mathbf{x}_{si}^t \mathbf{G}_s\right)^t \left(\sum\nolimits_{i\in n_s} \mathbf{x}_{si}^t \mathbf{G}_s\right)\right]$$
$$= \mathbb{E}\left[\sum\nolimits_{i\in n_s} (\mathbf{x}_{si}^t \mathbf{G}_{si})^2\right]$$
$$= \mathbb{E}\left[\sum\nolimits_{i\in n_s} \boldsymbol{m}\right]$$
$$= \boldsymbol{n_s m}$$

It follows that we obtain again:

$$\mathbb{E}[\boldsymbol{z_1 z_2}] = \frac{\mathbf{1}}{\sqrt{\boldsymbol{n_1 n_2}}} \frac{\boldsymbol{\sigma_g}}{\boldsymbol{m}} (\boldsymbol{n_s m}) + \frac{\boldsymbol{\sigma_e n_s}}{\sqrt{\boldsymbol{n_1 n_2}}} = \frac{(\boldsymbol{\sigma_g} + \boldsymbol{\sigma_e})\boldsymbol{n_s}}{\sqrt{\boldsymbol{n_1 n_2}}} = \frac{\boldsymbol{\rho N_s}}{\sqrt{\boldsymbol{n_1 n_2}}}$$

##### ***Simulation***

We simulated a series of 100 replicates each including 100,000 SNPs genotyped for 50,000 individuals. Coding allele frequency was uniformly sampled from [0.05, 0.95] and all SNPs were generated independently. For each replicates we generated two phenotypes with heritability drawn uniformly from [0.1, 0.75] and considered genetic correlation $r_g$ in [0.05, 0.5, 0.8]. We set the shared environmental variance $r_e$ so that the overall phenotypic correlation $r = r_g + r_e$ is equal to 0.6. With those parameters settings the phenotypic correlation can vary from high heritability, with $r$ fully explained by the genetic correlation to low heritability, with $r$ fully explained by the shared environment. We assumed only a small number $N_c$ of the total number of SNPs to be causal with effect sizes drawn from the multivariate normal distribution with mean 0 and covariance $\sigma_g = r_g \sqrt{h_1^2 h_2^2 / N_c}$:

$$\begin{pmatrix} h_1^2 & \sigma_g \\ \sigma_g & h_2^2 \end{pmatrix}$$

#### Sample size inference for linear regression

Let's consider a vector $\mathbf{X} = (\mathbf{1}, \mathbf{G}, \mathbf{U})$ of length $n$, where $\mathbf{1}$ is a vector of 1, $\mathbf{G}$ is a genetic variant. For simplicity we assume $\mathbf{U}$ is a covariate that captures the aggregated effect of all other predictors in the model. When testing the association between $\mathbf{X}$ and an outcome $\mathbf{Y}$, the variance of the least square estimates $\widehat{\boldsymbol{\beta}} = (\widehat{\boldsymbol{\beta}}_{\mathbf{0}}, \widehat{\boldsymbol{\beta}}_G, \widehat{\boldsymbol{\beta}}_U)$ equals:

$$\boldsymbol{var}(\widehat{\boldsymbol{\beta}}) = \boldsymbol{\sigma}_e^2(\mathbf{X}^t\mathbf{X})^{-1} = \boldsymbol{\sigma}_e^2\begin{pmatrix} \sum_N \mathbf{1}^2 & \sum_N \boldsymbol{G} & \sum_N \boldsymbol{U} \\ \sum_N \boldsymbol{G} & \sum_N \boldsymbol{G}^2 & \sum_N \boldsymbol{GU} \\ \sum_N \boldsymbol{U} & \sum_N \boldsymbol{GU} & \sum_N \boldsymbol{U}^2 \end{pmatrix}^{-1}$$

For large sample size and assuming independence between predictors, the covariance matrix simplified to:

$$\boldsymbol{var}(\widehat{\boldsymbol{\beta}}) \approx \frac{\boldsymbol{\sigma}_e^2}{\boldsymbol{n}}\begin{pmatrix} \mathbf{1} & \boldsymbol{\mu}_G & \boldsymbol{\mu}_U \\ \boldsymbol{\mu}_G & (\boldsymbol{\mu}_G^2 + \boldsymbol{\sigma}_G^2) & (\boldsymbol{\mu}_G\boldsymbol{\mu}_U) \\ \boldsymbol{\mu}_U & (\boldsymbol{\mu}_G\boldsymbol{\mu}_U) & (\boldsymbol{\mu}_U^2 + \boldsymbol{\sigma}_U^2) \end{pmatrix}^{-1}$$

It follows that:

$$\boldsymbol{var}(\widehat{\boldsymbol{\beta}}_G) \approx \frac{\boldsymbol{\sigma}_e^2}{\boldsymbol{n}} \frac{\boldsymbol{det}\begin{pmatrix} \mathbf{1} & \boldsymbol{\mu}_U \\ \boldsymbol{\mu}_U & (\boldsymbol{\mu}_G^2 + \boldsymbol{\sigma}_U^2) \end{pmatrix}}{\boldsymbol{det}\begin{pmatrix} \mathbf{1} & \boldsymbol{\mu}_G & \boldsymbol{\mu}_U \\ \boldsymbol{\mu}_G & (\boldsymbol{\mu}_G^2 + \boldsymbol{\sigma}_G^2) & (\boldsymbol{\mu}_G\boldsymbol{\mu}_U) \\ \boldsymbol{\mu}_U & (\boldsymbol{\mu}_G\boldsymbol{\mu}_U) & (\boldsymbol{\mu}_U^2 + \boldsymbol{\sigma}_U^2) \end{pmatrix}}$$

$$\approx \frac{\boldsymbol{\sigma}_e^2}{\boldsymbol{n}} \frac{(\boldsymbol{\mu}_G^2 + \boldsymbol{\sigma}_U^2) - \boldsymbol{\mu}_U^2}{(\boldsymbol{\mu}_G^2 + \boldsymbol{\sigma}_G^2)(\boldsymbol{\mu}_U^2 + \boldsymbol{\sigma}_U^2) - (\boldsymbol{\mu}_G\boldsymbol{\mu}_U)^2 - \boldsymbol{\mu}_G^2(\boldsymbol{\mu}_U^2 + \boldsymbol{\sigma}_U^2) + \mathbf{2}\boldsymbol{\mu}_G\boldsymbol{\mu}_U(\boldsymbol{\mu}_G\boldsymbol{\mu}_U) - \boldsymbol{\mu}_U^2(\boldsymbol{\mu}_G^2 + \boldsymbol{\sigma}_G^2)}$$

$$\approx \frac{\boldsymbol{\sigma}_e^2}{\boldsymbol{n}} \frac{\boldsymbol{\sigma}_U^2}{\boldsymbol{\sigma}_G^2\boldsymbol{\sigma}_U^2}$$

$$\approx \frac{\boldsymbol{\sigma}_e^2}{\boldsymbol{n}\boldsymbol{\sigma}_G^2}$$

The sample size $n$ can therefore be approximated by $\frac{\sigma_e^2}{var(\widehat{\boldsymbol{\beta}}_G)\sigma_G^2}$. As showed in **Figure S11**, this approximation shows reasonable accuracy as long as the provided variant frequency of the variant is within ±0.01 of the true frequency. For larger differences, the estimated $n$ can vary substantially for relative rare variants (i.e. MAF<5%).

#### Sample size inference for logistic regression

A relation exists between maximum likelihood estimation using Fisher scoring and weighted least squares estimation[9]. Indeed, maximum likelihood equation for the $(t+1)^{\text{th}}$ iteration has the form: $\boldsymbol{\beta}^{(t+1)} = \left(\mathbf{X}'\mathbf{W}^{(t)}\mathbf{X}\right)^{-1}\mathbf{X}'\mathbf{W}^{(t)}\mathbf{z}^{(t)}$, where $\mathbf{z}^{(t)}$ is the linearized form of the logit link function for the sample data. Using this formulation, the standard error estimate equals $\hat{\sigma}_\beta^2 \approx 1/(np(1-p)\sigma_G^2)$, where $p$ is the in-sample proportion of cases. Hence, similarly to the linear regression case, we propose using the term $1/\left(\hat{\sigma}_{\beta_G}^2\sigma_G^2\right)$ as a proxy for $W = np(1-p)$, which is proportional to sample size. Consider $w_F = n_F p_F(1-p_F)$, the parameter $W$ obtained for the full sample, where $n = n_F$ and $p = p_F$. In theory, $w^*$, a given occurrence of $W$, can equals $w_F$, even though $n \neq n_F$ and $p \neq p_F$. However, assuming the number of cases and controls can only decrease, one can show that $w^* \leq w_F$, and therefore $W$ remains effective as a tool to filter SNP which statistics has been derived with missing individuals. Indeed, $w_F$ can be written as a function of $n_{ca}$ and $n_{co}$, the number of cases and controls in the full sample:

$$\boldsymbol{w_F = np(1-p) = (n_{ca}+n_{co})\frac{n_{ca}}{n_{ca}+n_{co}}\frac{n_{co}}{n_{ca}+n_{co}} = \frac{n_{ca}n_{co}}{n_{ca}+n_{co}}}$$

In comparison, $w^*$ is derived in situations where the number of cases or control differ using $n^*_{ca} = n_{ca} \times c_1$ and $n^*_{co} = n_{co} \times c_2$, where $c_1$ and $c_2$ are strictly positives:

$$\boldsymbol{w^* = \frac{n_{ca}n_{co}c_1c_2}{n_{ca}c_1 + n_{co}c_2} = \frac{n_{ca}n_{co}}{\frac{n_{ca}}{c_2}+\frac{n_{co}}{c_1}} = W\frac{n_{ca}+n_{co}}{\frac{n_{ca}}{c_2}+\frac{n_{co}}{c_1}}}$$

It follows that if $c_1$ and $c_2$ are both strictly smaller than 1, $w^* < w_F$.

As showed in **Figure S12**, this indicator provides a reasonable proxy proportional to the true sample size.

##### *Imputation of missing z-score statistics*

The number of SNPs reported in GWAS summary statistics commonly differs between studies. All proposed multivariate approaches require complete data, so that jointly analyzing multiple GWAS might result in the removal of large number of variants. In order to increase the number of SNPs in cross-trait analyses, we performed z-scores imputations at missing variants within each single GWAS using high accuracy python package (RAISS) we recently developed[10]. RAISS improved on pre-existing solutions to reach a level of imputation accuracy suitable for multitrait analysis[11]. RAISS uses external information on linkage disequilibrium (LD) from a reference panel and z-scores for typed variants included in the study. Briefly, let $Z_t \sim \mathcal{N}(0, A)$ denote the vector of z-scores for the typed variants, and $\Sigma_{t,t}$ denote the correlation matrix of typed genotypes. The vector of imputed z-scores $Z^*_i$ at non-typed variants can be computed as linear combination of typed z-scores $Z_t$ with weights $W = \Sigma_{i,t}\Sigma^{-1}_{t,t}$ where $\Sigma_{i,t}$ is the SNP correlation matrix including imputed and typed variants:

$$\boldsymbol{Z^*_i = \frac{W}{\sqrt{WAW^t}} Z_t}$$

The theoretical accuracy of the imputation is assessed using the variance of the conditional random variable $Z^*_{i|t}$ and defined as $r^2pred = 1 - \Sigma_{i|t}$, where $\Sigma_{i|t} = WAW^t$.

As demonstrated below, the imputed SNPs have the same expected covariance as the genotyped SNP. Consider two GWAS studies $y_1$ and $y_2$ and suppose (without loss of generality) that the sets of genotyped SNPs are identical in the two studies. Consider a SNP $j$ missing in the two studies. Hence, linear regression z-scores are computed as:

$$\boldsymbol{z^*_{1j} = \frac{\Sigma_{j,t}\Sigma^{-1}_{t,t}}{\sqrt{\Sigma_{j,t}\Sigma^{-1}_{t,t}A\Sigma^{-1}_{t,t}\Sigma^t_{j,t}}} Z_t}$$

$$\boldsymbol{z^*_{2j} = \frac{\Sigma_{j,u}\Sigma^{-1}_{u,u}}{\sqrt{\Sigma_{j,u}\Sigma^{-1}_{u,u}A\Sigma^{-1}_{u,u}\Sigma^t_{j,u}}} Z_u}$$

It follows that the covariance between the imputed z-score equals:

$$\begin{aligned}\mathbb{E}[\boldsymbol{z^*_{1j}z^*_{2j}}] &= \mathbb{E}\left[\frac{\boldsymbol{\Sigma_{j,t}\Sigma^{-1}_{t,t}}}{\boldsymbol{r_{1,pred}}}\boldsymbol{Z_{1t}}\left(\frac{\boldsymbol{\Sigma_{j,u}\Sigma^{-1}_{u,u}}}{\boldsymbol{r_{2,pred}}}\boldsymbol{Z_{2u}}\right)^{\boldsymbol{t}}\right] \\ &= \frac{\boldsymbol{1}}{\boldsymbol{r_{1,pred}r_{2,pred}}}\mathbb{E}\left[\boldsymbol{\Sigma_{j,t}\Sigma^{-1}_{t,t}Z_{1t}(\Sigma_{j,u}\Sigma^{-1}_{u,u}Z_{2u})^t}\right] \\ &= \frac{\boldsymbol{1}}{\boldsymbol{r_{1,pred}r_{2,pred}}}\mathbb{E}\left[\boldsymbol{\Sigma_{j,t}\Sigma^{-1}_{t,t}Z_{1t}Z^t_{2u}\Sigma^{-1}_{u,u}\Sigma^t_{j,u}}\right]\end{aligned}$$

$$= \frac{\mathbf{1}}{\boldsymbol{r_{1,pred} r_{2,pred}}} \boldsymbol{\Sigma_{j,t} \Sigma_{t,t}^{-1}} \mathbb{E}[\boldsymbol{Z_{1t} Z_{2u}^{t}}] \boldsymbol{\Sigma_{u,u}^{-1} \Sigma_{j,u}^{t}}$$

However, we have $\mathbb{E}[Z_{1t} Z_{2u}^{t}] = \mathbb{E}[z_{1j} z_{2j}] \Sigma_{t,u}$, where $\Sigma_{t,u}$ is the covariance matrix between SNPs $t$ and $u$. It follows that:

$$\begin{aligned}
\mathbb{E}[\boldsymbol{z_{1j}^{*} z_{2j}^{*}}] &= \frac{\mathbf{1}}{\boldsymbol{r_{1,pred} r_{2,pred}}} \boldsymbol{\Sigma_{j,t} \Sigma_{t,t}^{-1}} \left[\mathbb{E}[\boldsymbol{z_{1j} z_{2j}}] \boldsymbol{\Sigma_{t,u}}\right] \boldsymbol{\Sigma_{u,u}^{-1} \Sigma_{j,u}^{t}} \\
&= \frac{\mathbb{E}[\boldsymbol{z_{1j} z_{2j}}]}{\boldsymbol{r_{1,pred} r_{2,pred}}} \boldsymbol{\Sigma_{j,t} \Sigma_{t,t}^{-1} \Sigma_{t,u} \Sigma_{u,u}^{-1} \Sigma_{j,u}^{t}} \\
&= \frac{\mathbb{E}[\boldsymbol{z_{1j} z_{2j}}]}{\boldsymbol{r_{1,pred} r_{2,pred}}} \boldsymbol{\Sigma_{j,t} \Sigma_{t,u}^{-1} \Sigma_{j,u}^{t}} \\
&= \mathbb{E}[\boldsymbol{z_{1j} z_{2j}}]
\end{aligned}$$

As a naïve sanity check we plotted in **Figure S13** the QQplot for multiple traits before and after imputation and did not observed any inflation.

##### *Unbiased genetic covariance estimator by clusters*

We defined the genetic covariance by clusters for trait 1 and trait 2 as:

$$\rho_{g,cluster} = \frac{\beta_1^t \beta_2}{M}$$

Where $\beta_1$ and $\beta_2$ are the the vector of genetic effects of the M SNPs contained in the cluster. However to estimate properly this quantity from the observed $\hat{\beta}$, we need to take into account the bias introduce by sample overlap:

$$\frac{\mathbb{E}(\widehat{\beta}_1{}^T \widehat{\beta}_2)}{M} = \frac{1}{M} \sum_{j=1}^{M} \mathbb{E}\left( \frac{x_{j,1}{}^T y_1}{x_{j,1}{}^T x_{j,1}} \times \frac{x_{j,2}{}^T y_1}{x_{j,2}{}^T x_{j,2}} \right) = \frac{1}{M} \sum_{j=1}^{M} \mathbb{E}\left( \frac{x_{j,1}{}^T (\beta_1 x_{j,1} + \varepsilon_1)}{n_1} \times \frac{x_{j,2}{}^T (\beta_2 x_{j,2} + \varepsilon_2)}{n_2} \right)$$

$$= \frac{1}{M} \sum_{j=1}^{M} \mathbb{E}\left( \frac{x_{j,1}{}^T x_{j,1} \beta_{j,1}}{x_{j,1}{}^T x_{j,1}} \times \frac{x_{j,2}{}^T x_{j,2} \beta_{j,2}}{x_{j,2}{}^T x_{j,2}} + \frac{x_{j,1}{}^T \varepsilon_1}{n_1} \times \frac{\varepsilon_2{}^T x_{j,2}}{n_2} \right)$$

$$= \frac{1}{M} \sum_{j=1}^{M} \beta_{j,1} \beta_{j,2} + \mathbb{E}\left( \varepsilon_1{}^T \begin{pmatrix} I_{n_s} & O_{n_s \times (n_2 - n_s)} \\ O_{n_s \times (n_1 - n_s)} & O_{(n_1 - n_s) \times (n_2 - n_s)} \end{pmatrix} \varepsilon_2 \right)$$

$$= \frac{1}{M} \sum_{j=1}^{M} \beta_{j,1} \beta_{j,2} + \frac{n_s \rho}{n_1 n_2}$$

Hence an unbiased estimator of the mean genetic covariance is :

$$\mathbb{E}(\beta_1{}^T \beta_2) = \frac{\mathbb{E}(\widehat{\beta}_1{}^T \widehat{\beta}_2)}{M} - \frac{n_s \rho}{n_1 n_2}$$

***Tissue enrichment analysis***

We tested enrichment for SNPs belonging to regions of open chromatin (more likely to contain expressed genes[12,13]) in specific tissues in three cases: i) when comparing results across GWAS sets, ii) when comparing univariate results, and iii) when comparing results across clusters. For all analyses we used functional annotations on 127 Roadmap tissues and cell lines defined by integrating activating histone marks (H3K4me1, H3K4me3, H3K9ac, and H3K27ac) with a latent Dirichlet allocation model as implemented in FUN-LDA[14]. Let $R_1$,…, $R_{127}$ be the 127 Roadmap tissues. We are interested in identifying the Roadmap tissue $R_j$ with the highest enrichment in significant SNPs (*p*-value < $5 \times 10^{-8}$) from a given set of results $G$. We first define:

$$\boldsymbol{p_{G|R_j}} = \frac{\textbf{\#significant SNPs in functional component } \mathbf{R_j}}{\textbf{\#SNPs in functional component } \mathbf{R_j}}$$

where the functional component is defined by FUN-LDA score > 0.9 (probability to be functional >0.9) for $R_j$. To test whether there is an enrichment in the functional component of Roadmap tissue $R_j$, we compare $p_{G|R_j}$ against:

$$\boldsymbol{p_{G|R_{-j}}} = \frac{\textbf{\#significant SNPs in functional components excluding } \mathbf{R_j}}{\textbf{\#SNPs in functional components excluding } \mathbf{R_j}}$$

where functional components excluding $R_j$ is defined by FUN-LDA score < 0.1 (probability to be functional <0.1) for $R_j$ but >0.9 for any other tissues/cell types. The null hypothesis of our test is $H_0$: $p_{G|R_j} = p_{G|R_{-j}}$ *versus* $H_1$: $p_{G|R_j} > p_{G|R_{-j}}$. For each analysis, we applied a two-sample proportion test for each Roadmap tissue $R_j$ to derive the p-value, and reported the enrichment ratio $p_{G|R_j}/p_{G|R_{-j}}$. To account for the total number of tests, we further applied an FDR adjustment. When comparing results from GWAS sets we adjusted for 23 tests × 127 tissues/cell types. When comparing results from univariate analyses, we adjusted for 36 phenotypes × 127 tissues/cell types. Finally, when comparing clusters, we adjusted for 41 clusters × 127 tissues/cell types.

#### Figure S1. Distribution of the proposed statistics under the null

We simulated series of 10,000 replicates, each including z-score statistics for 5 (panels a-c) and 100 (panels d-f) phenotypes and 100K SNPs. For each replicate we applied the *Omnibus* test (panels a and d), the *sumZ* test using weights of 1 (panels c and f) and test *sumZ* test using weights equals to loading of the first principal component of the phenotypic correlation matrix. Left panels show the observed chi-square distribution in blue against the expected one in red. Right panels show the corresponding *p*-value histograms.

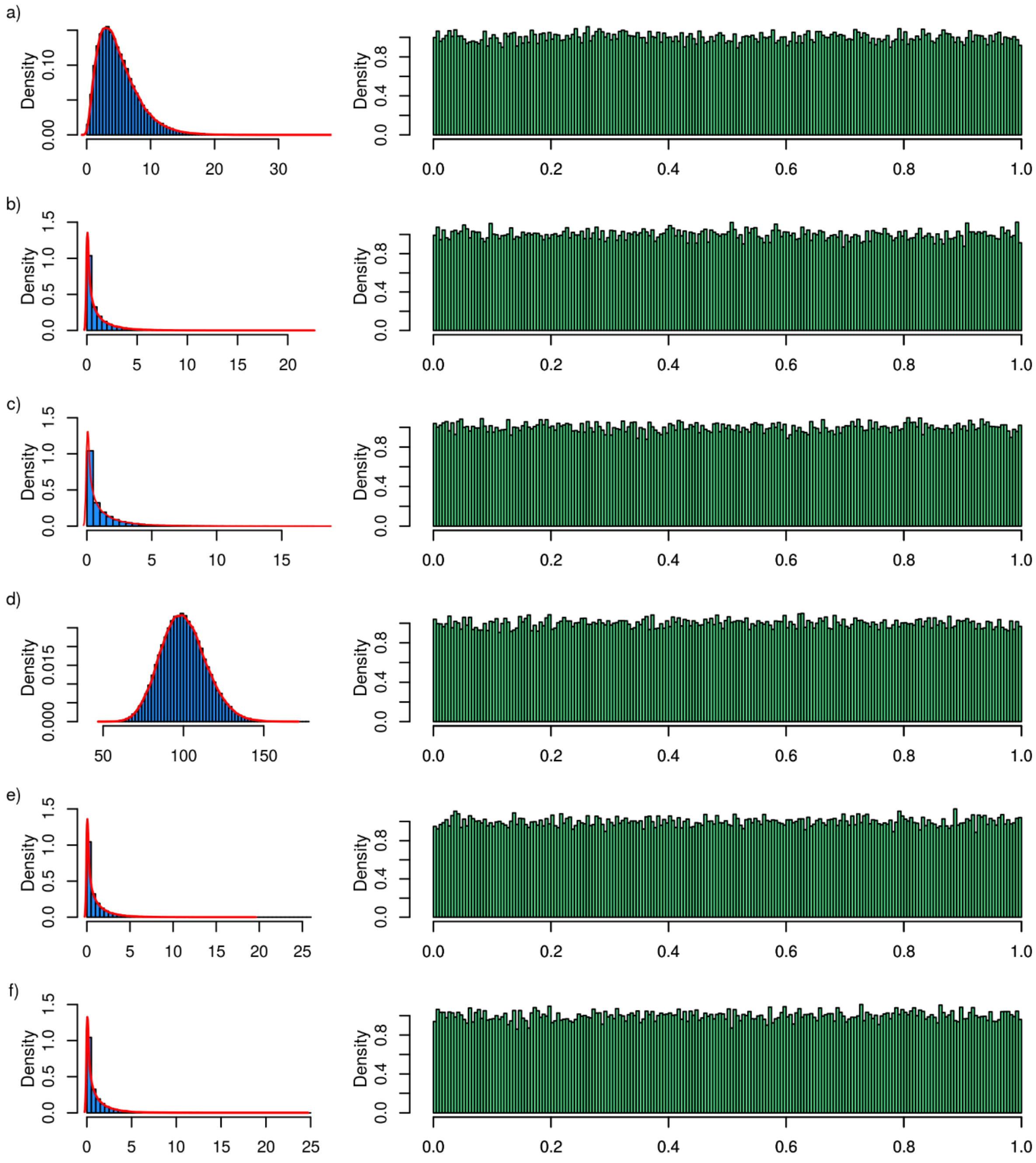

#### Figure S2. Solving singular covariance matrices

We simulated series of replicates, each including 20,000 individuals, 10,000 SNPs additively coded with frequencies ranging from 0.01 to 0.99, and 100 phenotypes. The phenotypes were drawn from a multivariate normal distribution with mean 0 and a variance-covariance $\mathbf{\Sigma_r}$ of rank 50. For each SNP, we computed the z-score of association with each phenotype using linear regressions and then derived *p*-values from the *Omnibus* test using three strategies to derive the inverse of $\mathbf{\Sigma_r}$: i) considering only eigenvalues greater than a specified threshold $\epsilon$ (left column), ii) replacing eigenvalues below a threshold $\epsilon$ by $\epsilon$ (middle column), and iii) adding a small value $\epsilon$ to the diagonal terms of $\mathbf{\Sigma_r}$ (right column). For each strategy we considered three different $\epsilon$ values: $10^{-3}$(first line), $10^{-6}$(second line), and $10^{-9}$(last line). The plots show the histogram of *ss*-value distribution.

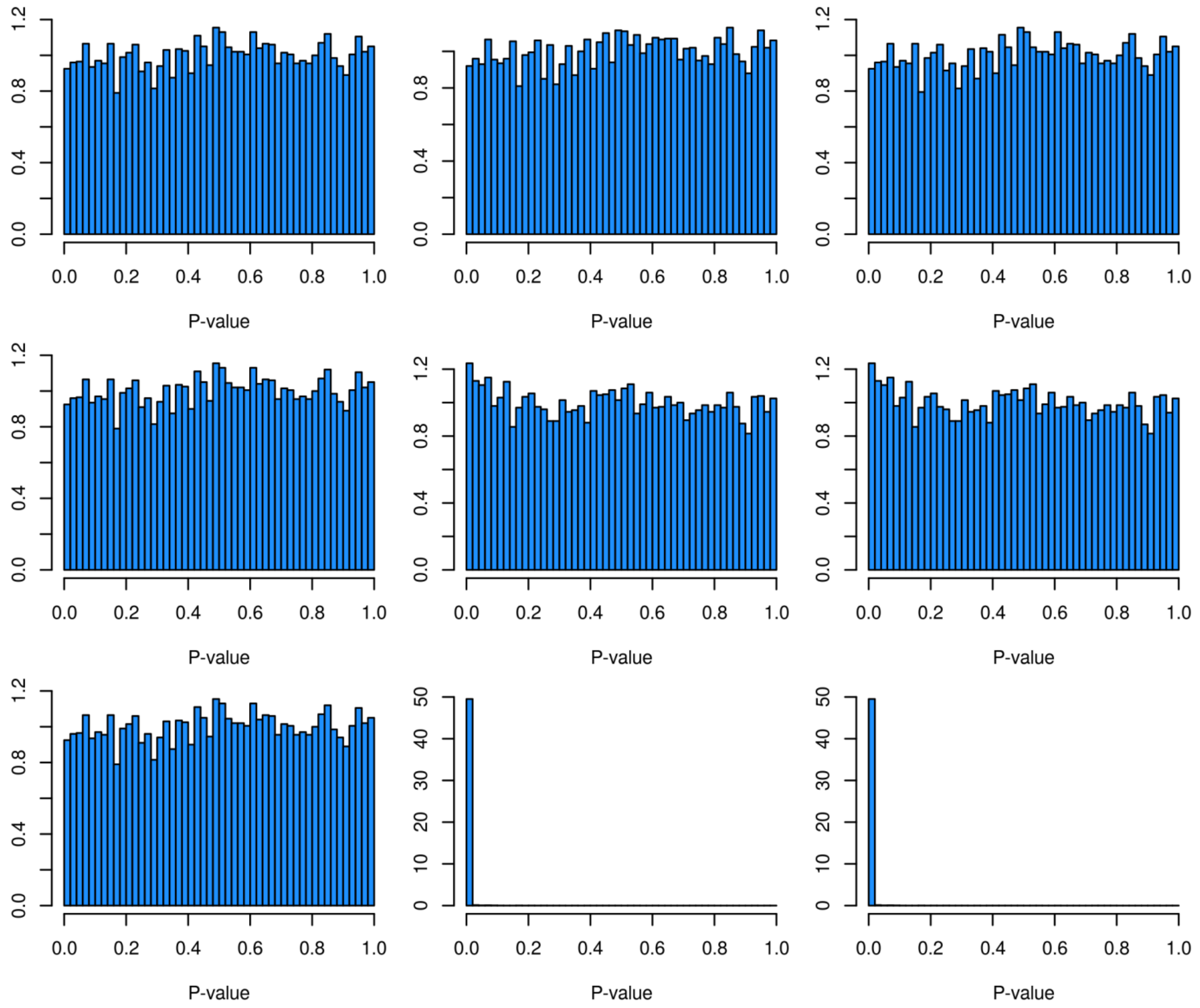

#### Figure S3. ICA component selection inflation control

We simulated 1000 z-score vectors under the null hypothesis for each of the 7 GWAS set. The z-score vectors under the null hypothesis follow a multivariate Gaussian of null mean and of covariance given by the intercept of the LDSC regression. The ICA was applied on the 1000 simulated z-scores. For each component, the sumZ test using the component as weight vector. We then selected the component yielding the most association as final component. We computed the p-value for the 1000 point for the optimal component. The y-axis represents the observed p-value quantiles with respect to the theoretical p-value quantiles.

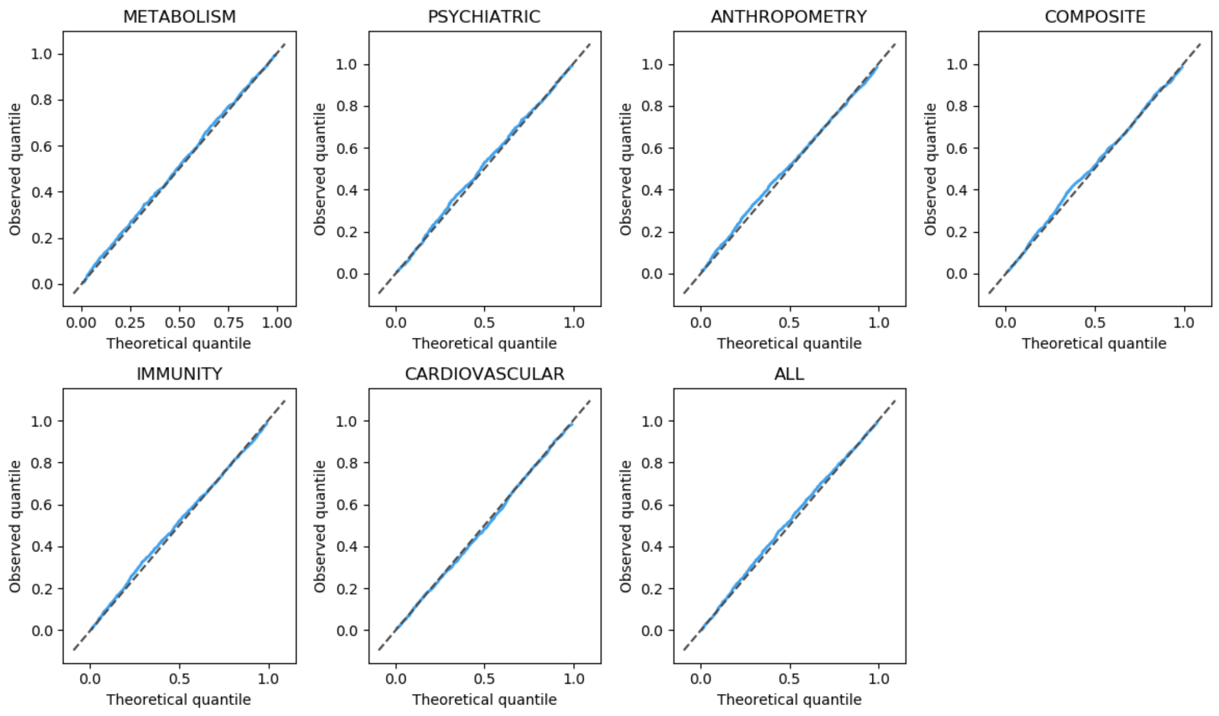

#### Figure S4. Summary statistics versus individual-level data tests under the null

We simulated replicates of 10,000 predictors for 500 (left column), 5,000 (middle column) and 10,000 individuals (right column). Predictors were drawn from a binomial with *n*=2 and probability varying in [0.01 – 0.99] to mimic genotype data, and further normalized to have mean 0 and variance 1. We then simulated a) 5, b) 20 and c) 100 correlated outcomes independent from the genotypes. To assess the impact of non-normal outcomes on the relationship between the summary-statistics based test and the individual-level data test, the outcomes were first drawn from a multivariate normal distribution with pairwise correlation ranging in [-0.7; 0.7] and then transform to non-normal distributions based on their quantiles, so that on average, 33% of them follow a uniform distribution, 33% a Laplace distribution, and 33% an exponential distribution. For each SNP, we first applied a Multivariate ANOVA (MANOVA, blue line). We then conducted association screenings for each single phenotype separately, and applied the proposed *Omnibus* test (purple line) and the Wilk's approximation (orange line) on the resulting summary statistics. The plots show the observed -log10(*p*-value) of each test against the expected value under the null.

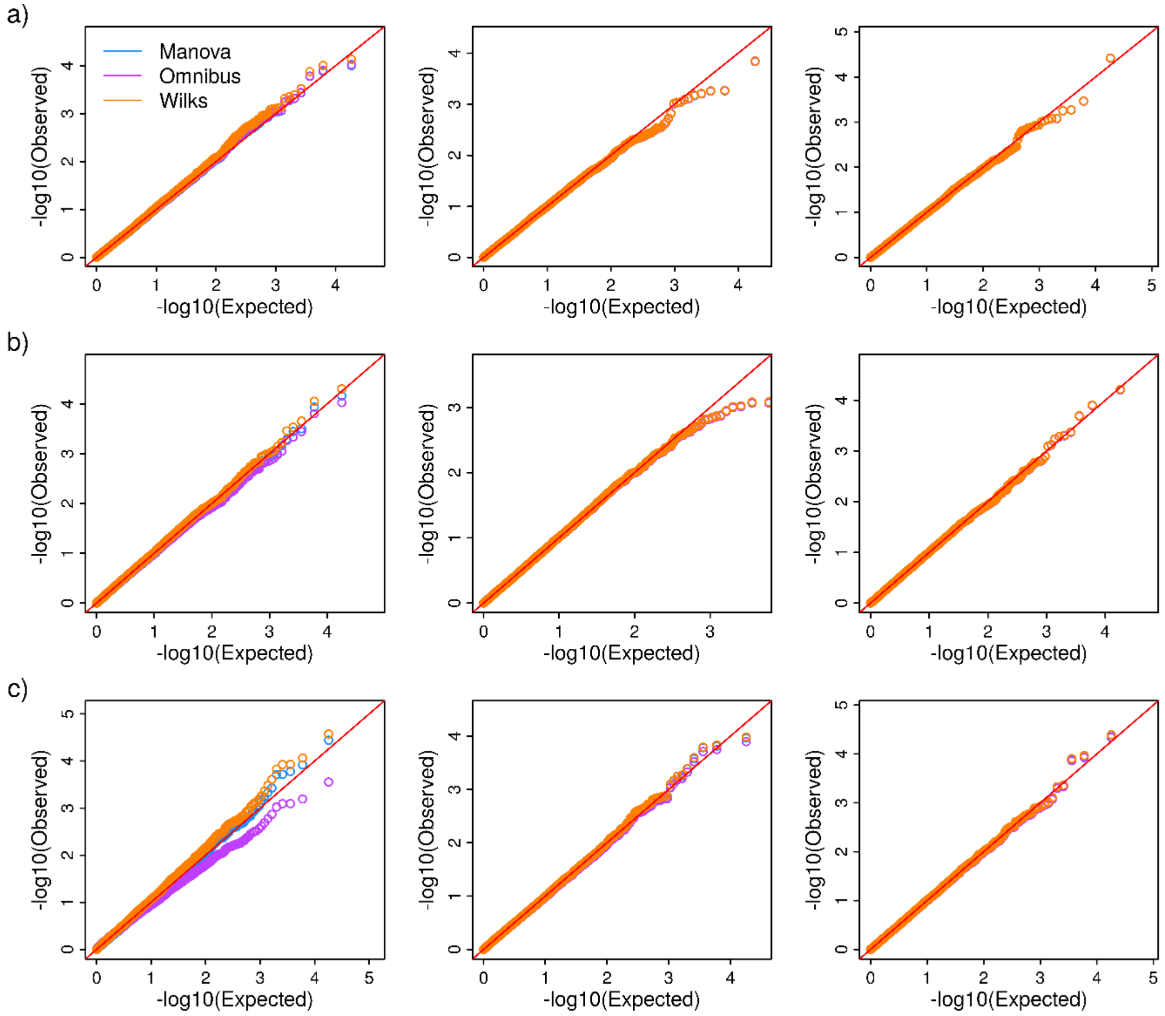

#### Figure S5. Summary statistics versus individual-level data tests under the alternative

We simulated replicates of 10,000 predictors for 500 (left column), 5,000 (middle column) and 10,000 individuals (right column). Predictors were drawn from a binomial with *n*=2 and probability varying in [0.01 – 0.99] to mimic genotype data, and further normalized to have mean 0 and variance 1. We then simulated a) 5, b) 20 and c) 100 correlated outcomes. We assumed that 10% of the predictors were causal, each causal predictor being randomly chosen and associated with up to 50% outcomes, randomly chosen with equal probability. Effect sizes for each causal predictor were drawn from a normal distribution with mean 0 and variance 0.0025 (i.e. the total phenotypic variance explained by the predictors). To assess the impact of non-normal outcomes on the relationship between the summary-statistics based test and the individual-level data test, the outcomes were first drawn from a multivariate normal distribution with pairwise correlation ranging in [-0.7; 0.7] and then transform to non-normal distributions based on their quantiles, so that on average, 33% of them follow a uniform distribution, 33% a Laplace distribution, and 33% an exponential distribution. For each SNP, we first applied a Multivariate ANOVA (MANOVA, blue line). We then conducted association screenings for each single phenotype separately, and applied the proposed *Omnibus* test (purple line) and the Wilk's approximation (orange line) on the resulting summary statistics. The plots show the observed -log10(*p*-value) of each test against the expected value under the null.

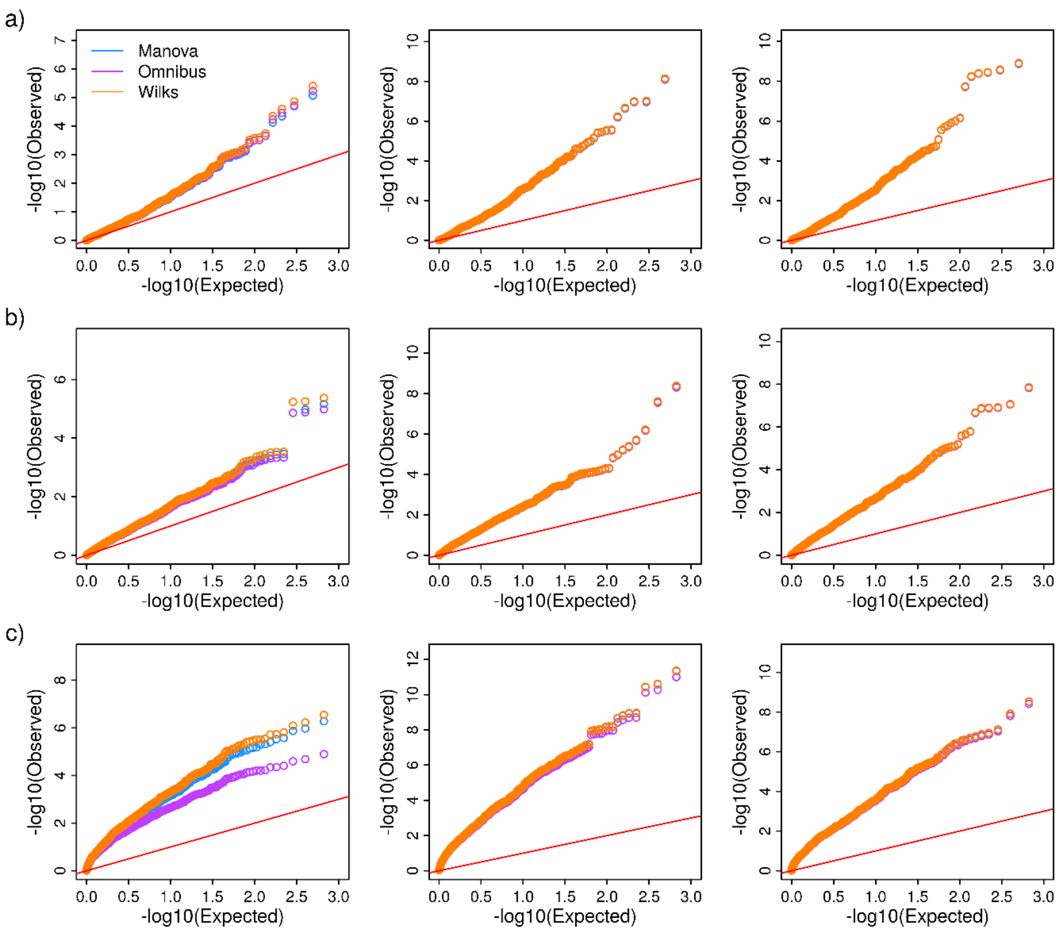

#### Figure S6. Validation of the summary-statistics approach in UK Biobank individuals

Multi-trait GWAS was performed in real data from the UK Biobank using five traits, 336,347 unrelated individuals, and 14,718 genotyped SNPs on Chromosome 20. We used two methods: MANOVA as implemented in PLINK, and the *Omnibus* test from JASS. The plot shows the -log10(*p*-values) of the Omnibus approach as a function of the -log10(*p*-values) derived with the MANOVA.

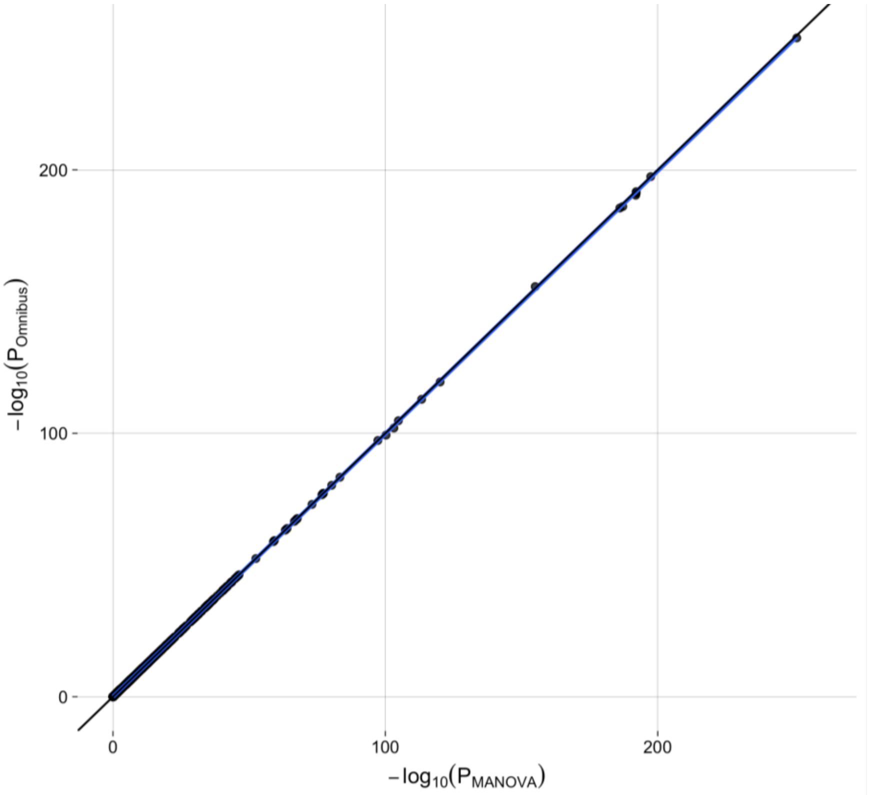

15

#### Figure S7. Impact of covariance bias on multivariate tests.

We simulated three correlated vectors of z-scores for 10,000 SNPs. We derived the multivariate test for each SNP using either the true covariance matrix ($\Sigma_{true}$), an upward-bias covariance matrix ($\Sigma_{upBias}$, left panels) or a downward-bias covariance matrix ($\Sigma_{downBias}$, right panels). For both scenarios we derived the *Omnibus* multivariate test using either the true or biased covariance matrix. Invalidity of the test based on the biased covariance matrix is illustrated in the resulting QQplots and genomic inflation factor $\lambda_{GC}$ (bottom panels).

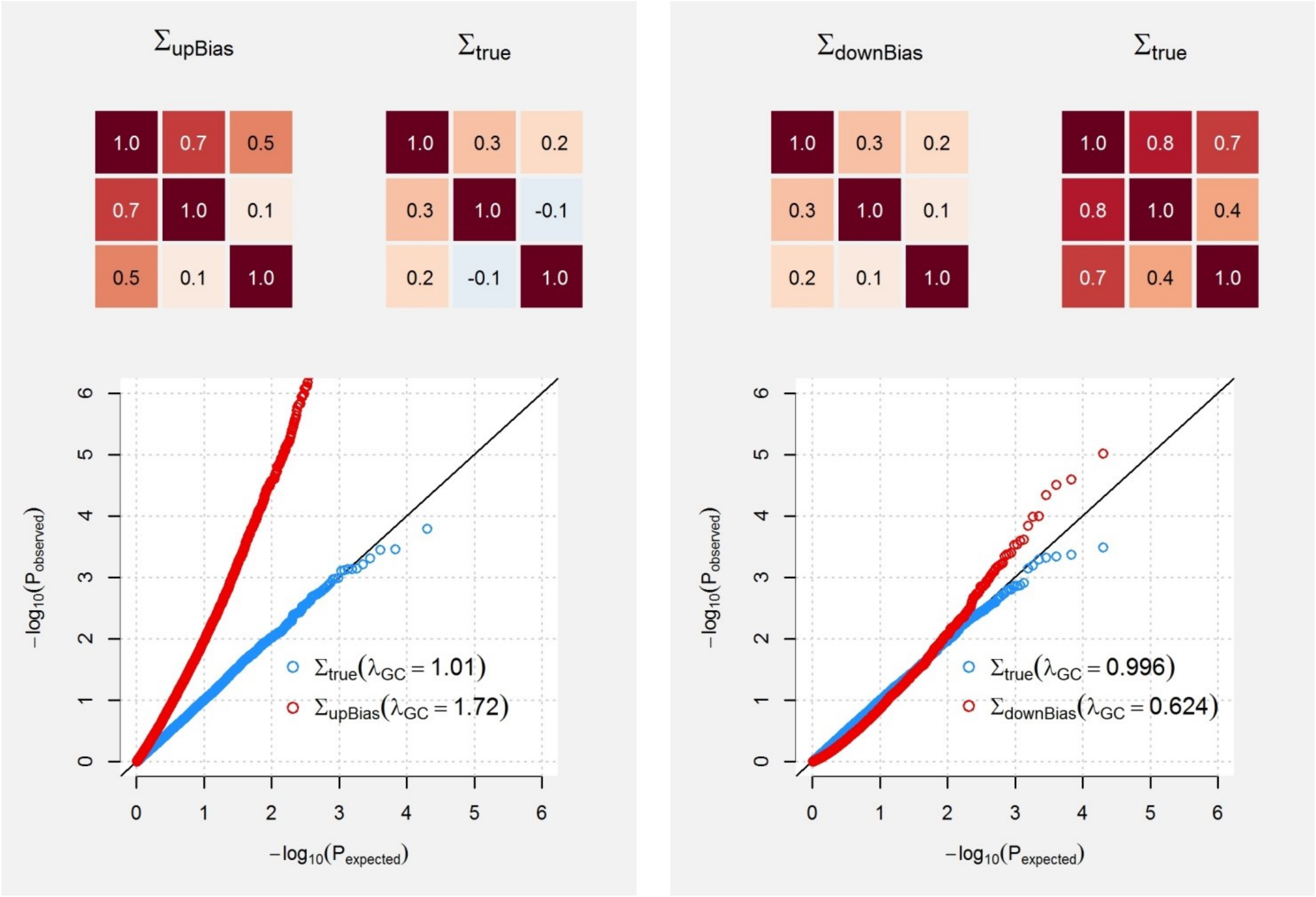

#### Figure S8. Impact of causal variants on the covariance estimation.

We simulated series of correlated z-scores for 100,000 SNPs from two outcomes $Y_1$ and $Y_2$. For each simulation we generated a matrix of true genetic effect $\mathbf{b} = (\boldsymbol{\beta_1} \quad \boldsymbol{\beta_2})$ of $m$ standardized and independent genotypes for two phenotypes $Y_1$ and $Y_2$ from a multivariate normal with means of 0 variance $h_1^2/m$, and $h_2^2/m$, respectively, and covariance $\sigma_g$, where $h_1^2 = 0.3$ and $h_2^2 = 0.6$ are the heritability of $Y_1$ and $Y_2$. We then generated $\hat{\mathbf{b}}$ defined as $\hat{\mathbf{b}} = \mathbf{b} + \boldsymbol{\varepsilon}$ where $\boldsymbol{\varepsilon}$ was also drawn from a multivariate normal with means 0, variance $1/N_1$ and $1/N_2$, respectively, and covariance $r_e\, N_s/\sqrt{N_1 N_2}$, where $N_1$, $N_2 = N_1/2$ and $N_s = N_2/2$ are the sample sizes for $Y_1$ and $Y_2$ , and the number of shared samples, respectively, and $r_e = 0.4$ is the correlation between $Y_1$ and $Y_2$. Finally, we derived the expected z-score for each genotype $\mathbf{z} = \left(\widehat{\boldsymbol{\beta}}_{\mathbf{1}}\sqrt{N_1} \quad \widehat{\boldsymbol{\beta}}_{\mathbf{2}}\sqrt{N_2}\right) = (\boldsymbol{z_1} \quad \boldsymbol{z_2})$, and $\sigma_z = cov(\boldsymbol{z_1}, \boldsymbol{z_2})$, the covariance between $\boldsymbol{z_1}$ and $\boldsymbol{z_2}$. The left panel (a) show the covariance between z-score for null variants in red, and the observed covariance between all z-scores except those harboring a *p*-value below a given threshold (0 (i.e. no SNP removed), 5x10$^{-8}$, 5x10$^{-5}$, 5x10$^{-4}$, 5x10$^{-3}$, 1x10$^{-2}$, and 5x10$^{-8}$). The right panel (b) shows the same results while assuming only 10% of the variants are causal, while all remaining have $\mathbf{b} = \mathbf{0}$.

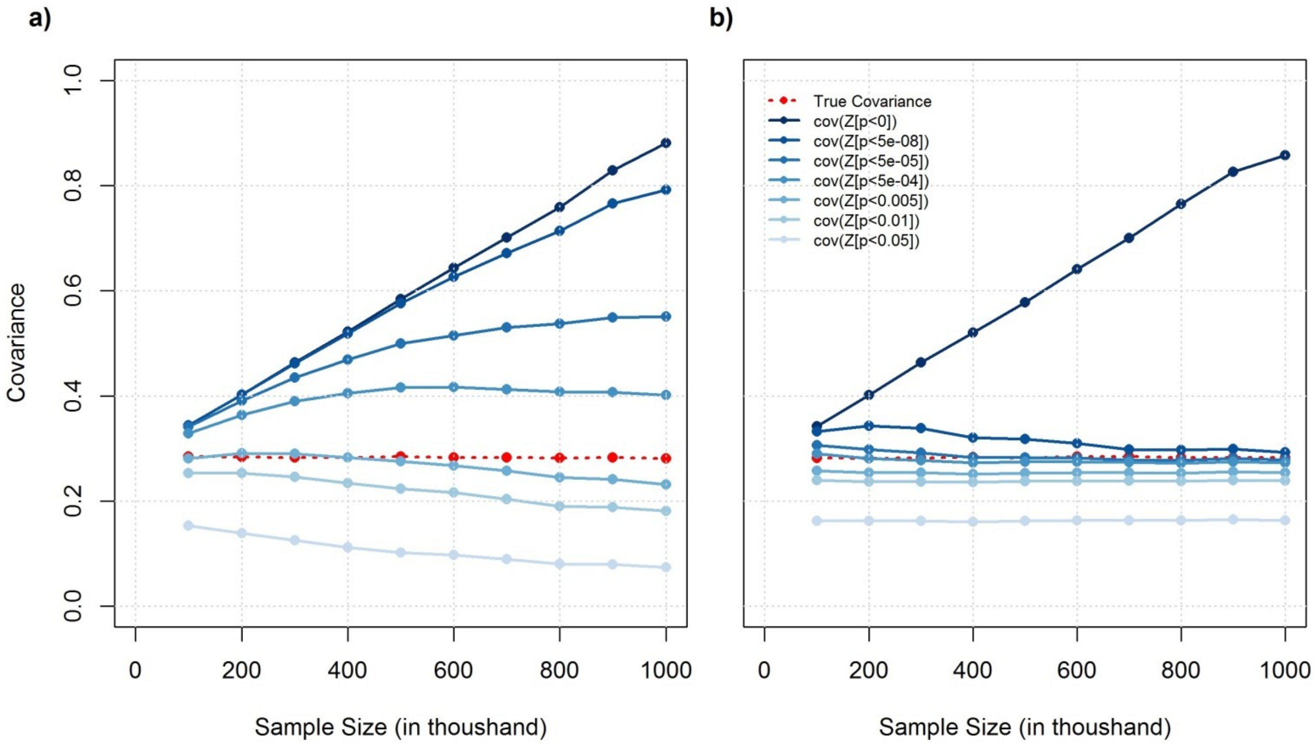

#### Figure S9. Validation of the LDSC covariance estimate accuracy in UK Biobank

We compared LDSC estimates of covariance between summary statistics against its expected value, which equals $\rho N_s/\sqrt{N_1 N_2}$, where $\rho$ is the phenotypic correlation among overlapping sample and $N_s$, $N_1$ and $N_2$ are the sample overlap, the sample size for phenotype 1 and the sample size for phenotype 2, respectively. We used individual-level data for five anthropometric traits and 619,017 SNPS measured in 336,347 individuals from the UK Biobank cohort. We first considered scenarios with complete sample overlap (i.e. $N_s = N_1 = N_2$, panels a-d), so that LDSC estimate is expected to equal the phenotypic correlation. We compared estimates while using 100% (a), 50% (b), 10% (c) and 1% (ds) of the total sample size respectively. The lower matrix triangle in pale red are estimated phenotypic correlation from individual-level data, while the upper triangle in pale blue shades are estimated correlation derived using the LDSC. We then considered scenario where sample overlap is only partial by sub-sampling individuals only for BMI, using 100% (e),50% (f), 10% (g) and 1% (h) of the total sample for that phenotype. The first row, in red, is the expected GWAS covariance knowing $\rho$, $N_s$, $N_1$ and $N_2$. The second row, in blue, is the estimate derived using LDSC.

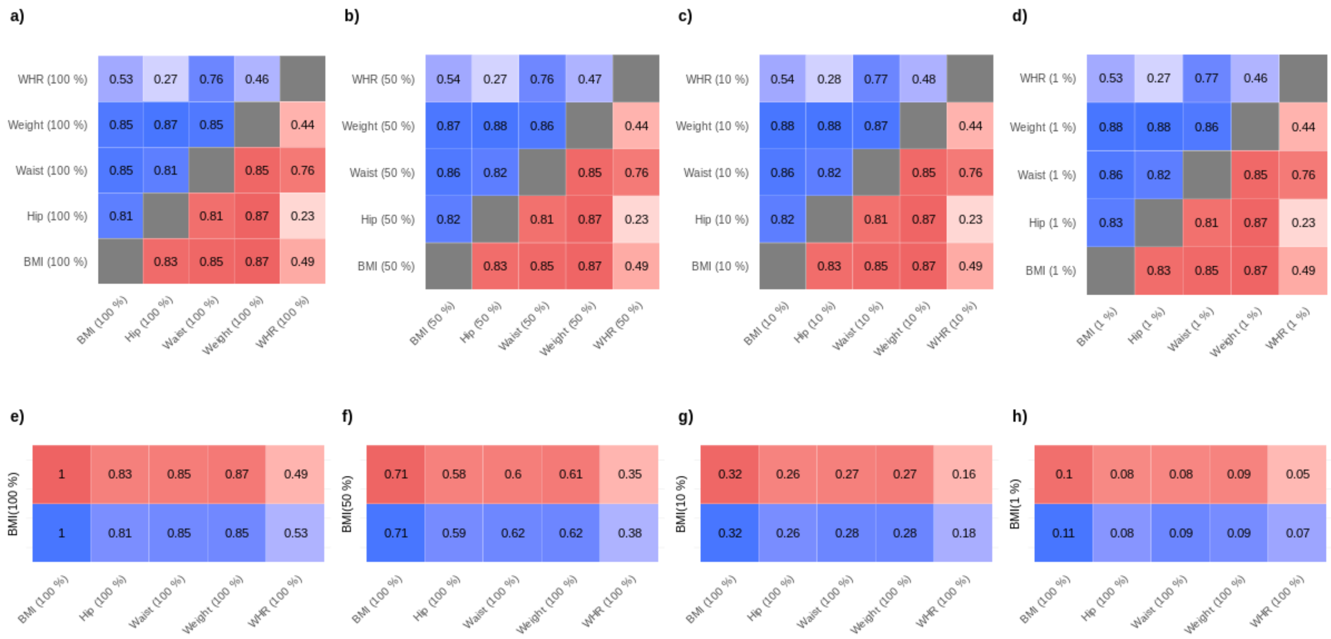

#### Figure S10. Bias due to sample size heterogeneity in the GLG consortium GWAS data.

We performed the *Omnibus* test to the four traits from the GLG consortium: high density lipoprotein (HDL), low density lipoprotein (LDL), total cholesterol (TC), and total triglyceride (TG) using all available SNP with complete summary statistics. We plotted the median chi-squared of the resulting test across bin of SNPs defined based the per-SNP standard deviation in sample size across the four traits ($\sigma_{\text{sample size}}$) for SNPs with a minor allele frequency (MAF) below 5% (a), and above 5% (b). The red dashed line indicates the expected 4 degree of freedom chi-square under the null. The shade of blue is proportional to the number of SNPs in each bin.

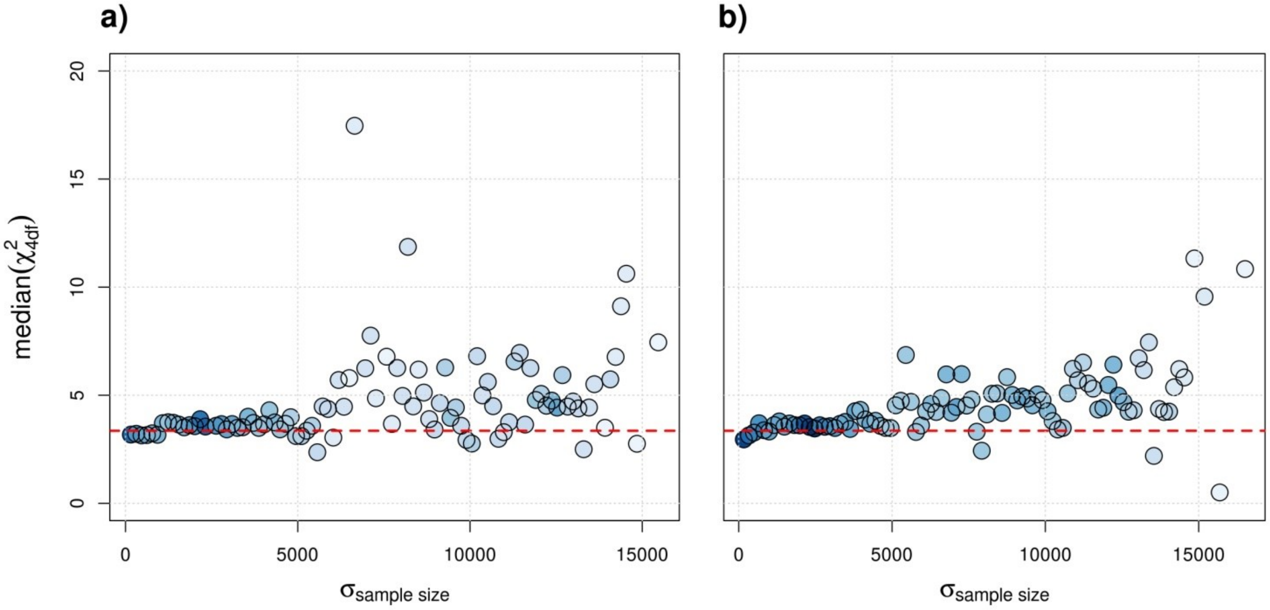

#### Figure S11. Impact of allele frequency error on sample size inference.

We simulated 1,000 replicates including 20,000 individuals were a phenotype $Y$ is simulated independently of a genotype $G$ with frequency randomly sampled in [0.001, 0.999]. For each replicate we tested for association between $G$ and $Y$ and we inferred $w$ the weights, which equals the sample size times a constant, using the standard error of the effect estimate and either the in-sample allele frequency or the in-sample allele plus a noise term sample from a uniform with min and max in [0.01, 0.05, 0.10]. When allele frequency plus noise was larger or smaller than 0 or 1, the value was set to 0.001 and 0.999, respectively. Upper and lower panels show the inferred sample size as a function of the true allele frequency when using the identity and logit link functions for modelling and testing for association, respectively. Note that for the later, for each replicate, we simulated 50,000 individuals and considered a disease prevalence of 25%. We then randomly sampled 10,000 cases and 10,000 controls to form replicates of 20,000 individuals. Also, for the sake of comparison, we scaled $w$ by a constant in the logit model so that the target is the true sample size.

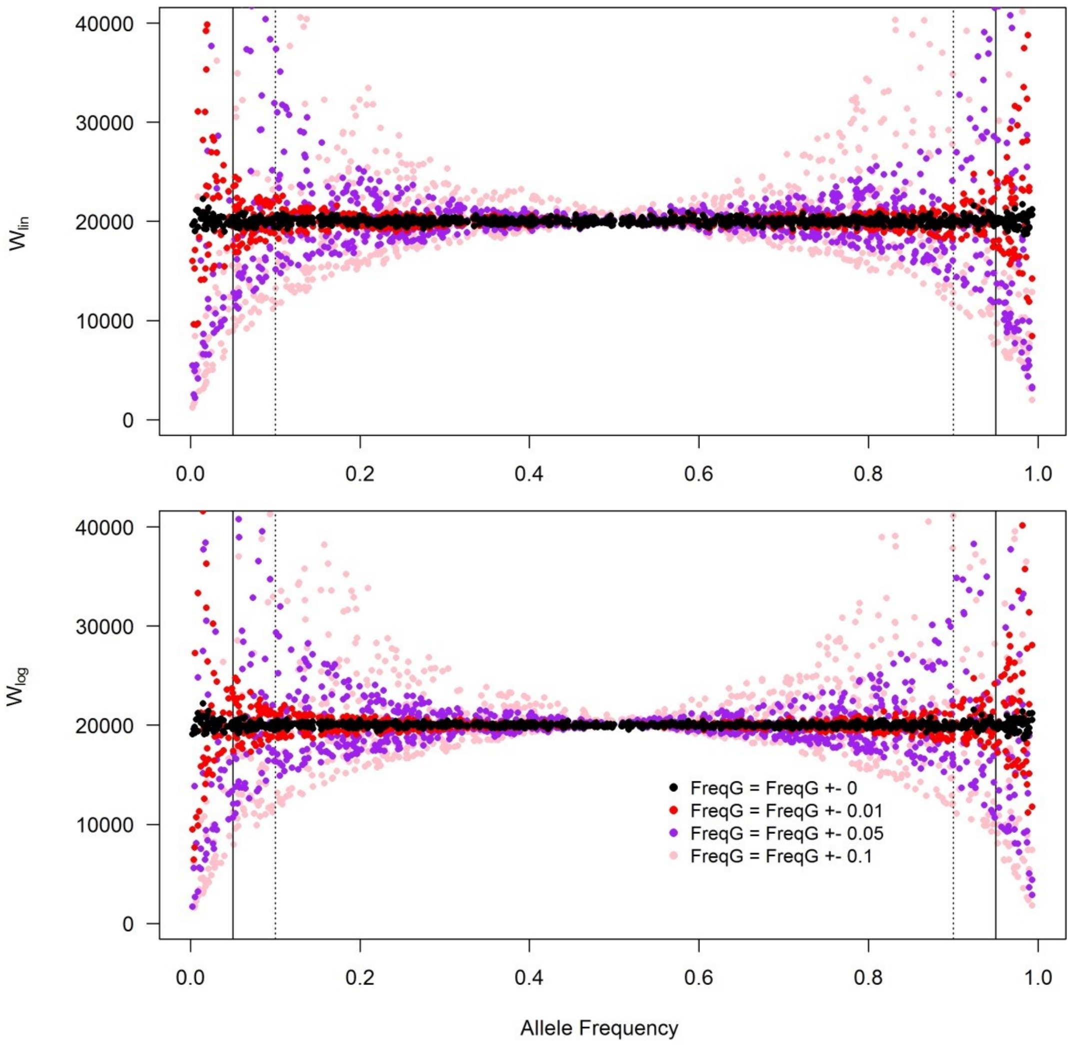

#### Figure S12. Impact of case/control ratio misspecification on sample size inference.

We simulated 1,000 replicates including 50,000 individuals. For each individual we generated a disease status assuming a prevalence of 25% and an independent genotype $G$ with frequency randomly sampled in [0.001, 0.999]. For each replicate we randomly sampled 5,000 cases and 20,000 controls and tested for association between $G$ and $Y$ using these samples and after sub-sampling cases (i.e. using a sub-sample of the 5,000 cases and the 20,000 controls, top panel) or controls (i.e. using the 5,000 cases and a sub-sample of the 20,000 controls, bottom panel), in order to mimic situation where either cases or controls would be missing for some SNPs. For each experiment we inferred $W_{log} = Np(1-p)$, where $N$ is the sample size and $p$ is the case-control ratio, using the standard error of the effect estimate and the in-sample allele frequency. In both situations (sub-sampling cases or sub-sampling controls), $W_{log}$ is decreasing with increasing percentage of missingness.

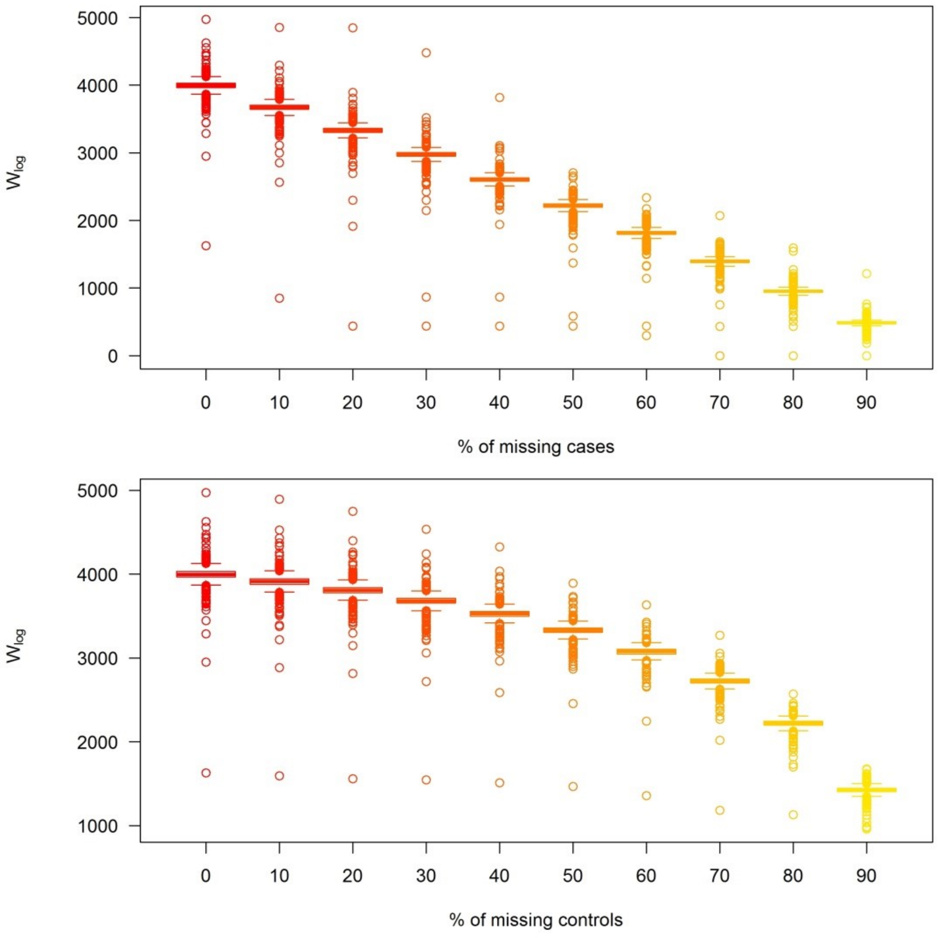

#### Figure S13. Effect of imputation of missing z-scores on the Omnibus test statistic.

Each line corresponds to a GWAS set. The left column is the empirical quantile of the *Omnibus* statistic versus the theoretical quantile before imputation. The middle column is the same after imputation. The right column is the empirical quantile of the *Omnibus* statistic before imputation versus the same quantity after imputation.

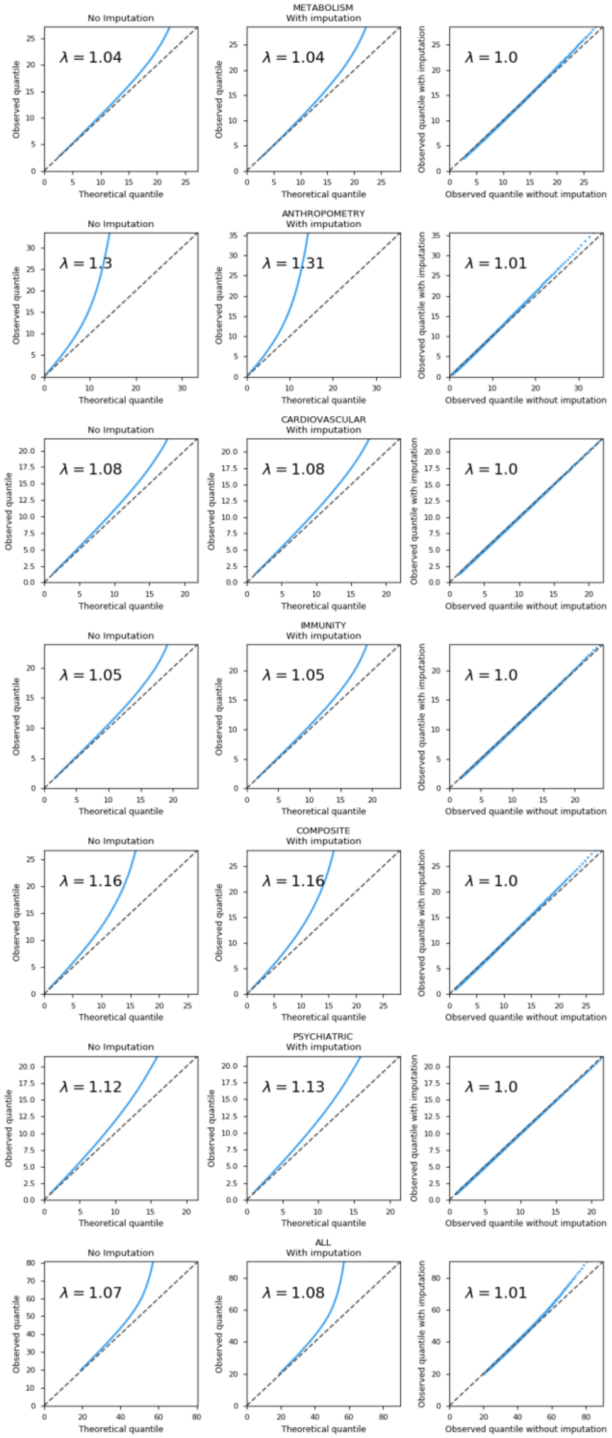

#### Figure S14. Signal comparison for all phenotypes.

The upper panel shows independent variants detected across phenotype groups and across approaches represented as an *UpSetR* visualization. Matrix lines correspond to a test, each column to a set of significant variants. For each set, the test for which variants are significant are represented with a black dot on the test line. The barplot on the left of the matrix represents the number of significant independent signals detected by each approach. The barplot on the top of the matrix represents the cardinality of the sets. The sets are ordered by cardinality from the largest to the leftmost to the smallest to the rightmost. The bottom panels show quadrant plots, i.e. the -log10($p$-value) for the most significant SNP per region for the *Omnibus* test as a function of the -log10($p$-value) for the most significant SNP per region across all univariate GWAS. Complete results are presented in the left panel, and a zoom around the genome-wide significance threshold is presented on the right panel.

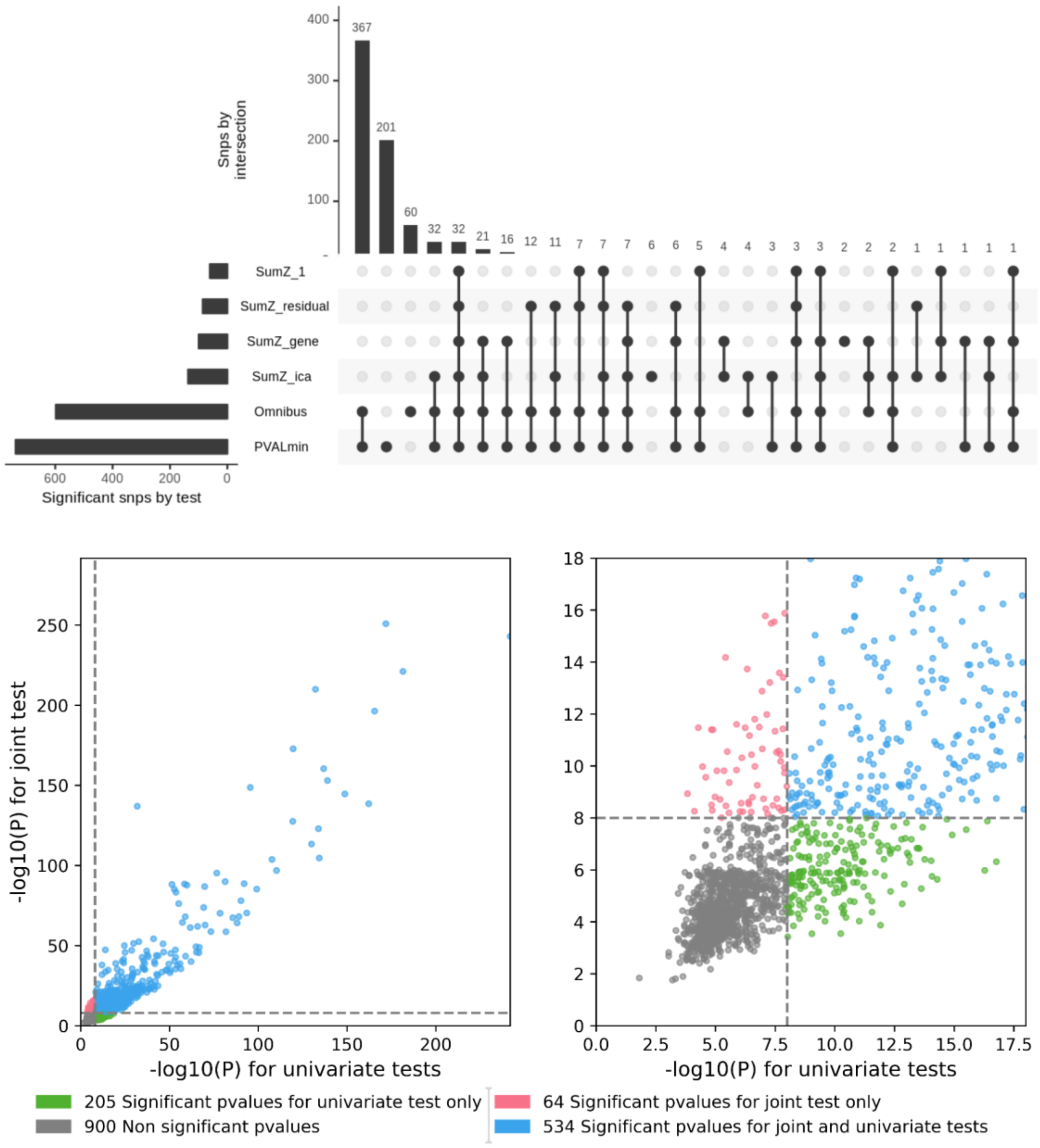

#### Figure S15. Signal comparison for anthropometric traits.

The upper panel shows independent variants detected across phenotype groups and across approaches represented as an *UpSetR* visualization. Matrix lines correspond to a test, each column to a set of significant variants. For each set, the test for which variants are significant are represented with a black dot on the test line. The barplot on the left of the matrix represents the number of significant independent signals detected by each approach. The barplot on the top of the matrix represents the cardinality of the sets. The sets are ordered by cardinality from the largest to the leftmost to the smallest to the rightmost. The bottom panels show quadrant plots, i.e. the -log10(*p*-value) for the most significant SNP per region for the *Omnibus* test as a function of the -log10(*p*-value) for the most significant SNP per region across all univariate GWAS. Complete results are presented in the left panel, and a zoom around the genome-wide significance threshold is presented on the right panel.

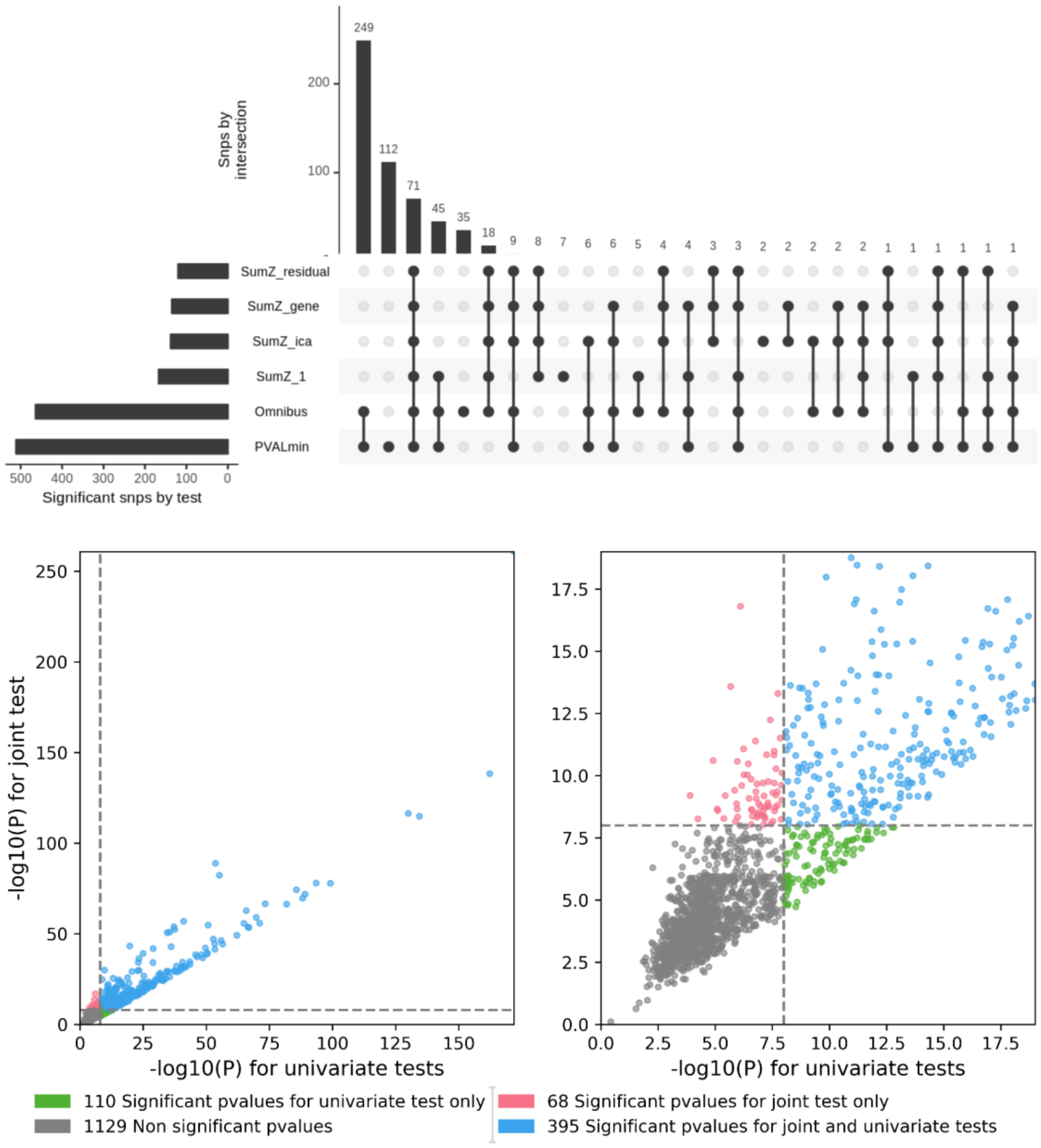

#### Figure S16. Signal comparison for cardiovascular phenotypes.

The upper panel shows independent variants detected across phenotype groups and across approaches represented as an *UpSetR* visualization. Matrix lines correspond to a test, each column to a set of significant variants. For each set, the test for which variants are significant are represented with a black dot on the test line. The barplot on the left of the matrix represents the number of significant independent signals detected by each approach. The barplot on the top of the matrix represents the cardinality of the sets. The sets are ordered by cardinality from the largest to the leftmost to the smallest to the rightmost. The bottom panels show quadrant plots, i.e. the -log10(*p*-value) for the most significant SNP per region for the *Omnibus* test as a function of the -log10(*p*-value) for the most significant SNP per region across all univariate GWAS. Complete results are presented in the left panel, and a zoom around the genome-wide significance threshold is presented on the right panel.

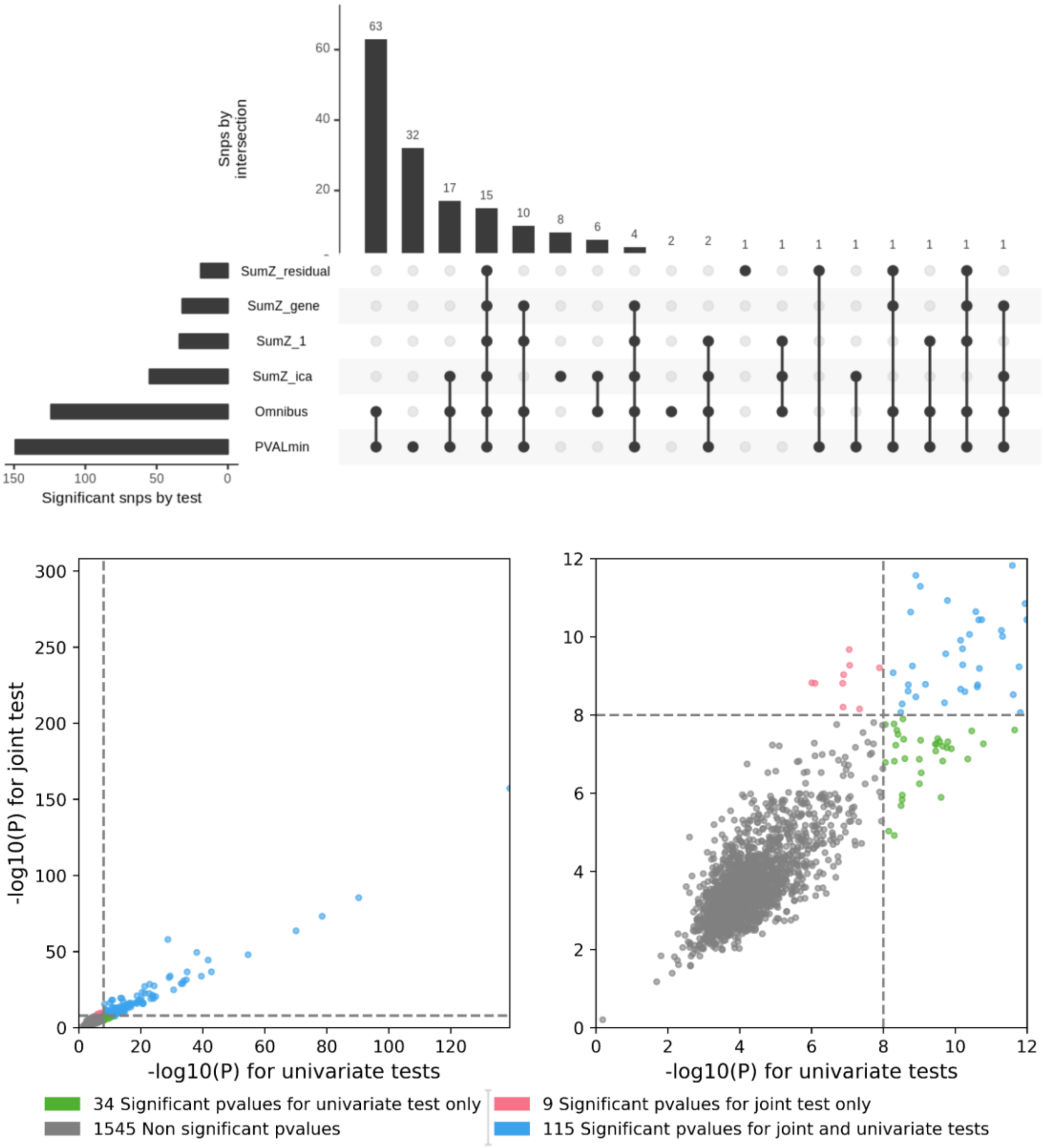

#### Figure S17. Signal comparison for composite phenotypes set.

The upper panel shows independent variants detected across phenotype groups and across approaches represented as an *UpSetR* visualization. Matrix lines correspond to a test, each column to a set of significant variants. For each set, the test for which variants are significant are represented with a black dot on the test line. The barplot on the left of the matrix represents the number of significant independent signals detected by each approach. The barplot on the top of the matrix represents the cardinality of the sets. The sets are ordered by cardinality from the largest to the leftmost to the smallest to the rightmost. The bottom panels show quadrant plots, i.e. the -log10(*p*-value) for the most significant SNP per region for the *Omnibus* test as a function of the -log10(*p*-value) for the most significant SNP per region across all univariate GWAS. Complete results are presented in the left panel, and a zoom around the genome-wide significance threshold is presented on the right panel.

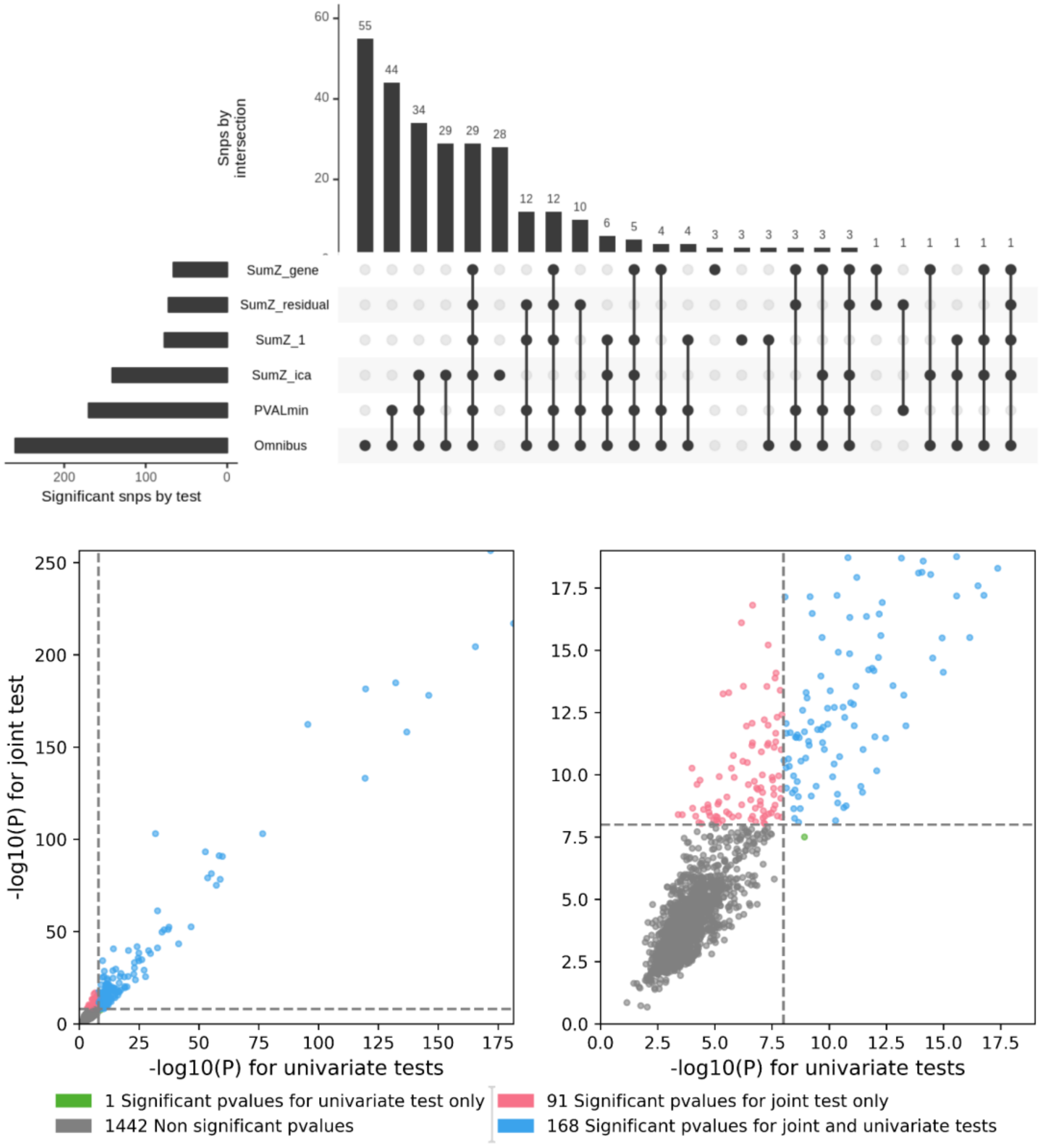

#### Figure S18. Signal comparison for immunity phenotypes.

The upper panel shows independent variants detected across phenotype groups and across approaches represented as an *UpSetR* visualization. Matrix lines correspond to a test, each column to a set of significant variants. For each set, the test for which variants are significant are represented with a black dot on the test line. The barplot on the left of the matrix represents the number of significant independent signals detected by each approach. The barplot on the top of the matrix represents the cardinality of the sets. The sets are ordered by cardinality from the largest to the leftmost to the smallest to the rightmost. The bottom panels show quadrant plots, i.e. the -log10(*p*-value) for the most significant SNP per region for the *Omnibus* test as a function of the -log10(*p*-value) for the most significant SNP per region across all univariate GWAS. Complete results are presented in the left panel, and a zoom around the genome-wide significance threshold is presented on the right panel.

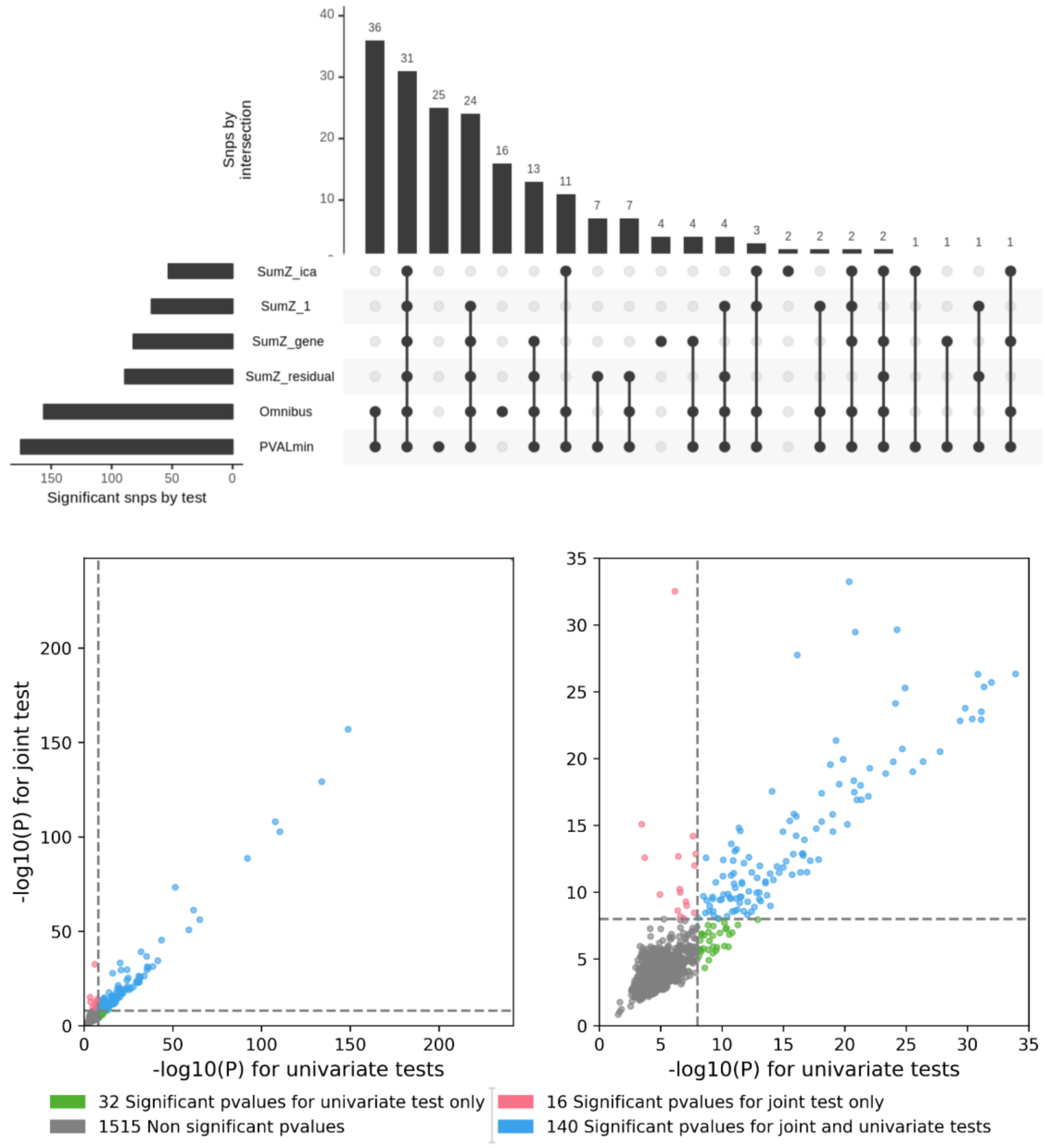

#### Figure S19. Signal comparison for metabolism phenotypes.

The upper panel shows independent variants detected across phenotype groups and across approaches represented as an *UpSetR* visualization. Matrix lines correspond to a test, each column to a set of significant variants. For each set, the test for which variants are significant are represented with a black dot on the test line. The barplot on the left of the matrix represents the number of significant independent signals detected by each approach. The barplot on the top of the matrix represents the cardinality of the sets. The sets are ordered by cardinality from the largest to the leftmost to the smallest to the rightmost. The bottom panels show quadrant plots, i.e. the -log10(*p*-value) for the most significant SNP per region for the *Omnibus* test as a function of the -log10(*p*-value) for the most significant SNP per region across all univariate GWAS. Complete results are presented in the left panel, and a zoom around the genome-wide significance threshold is presented on the right panel.

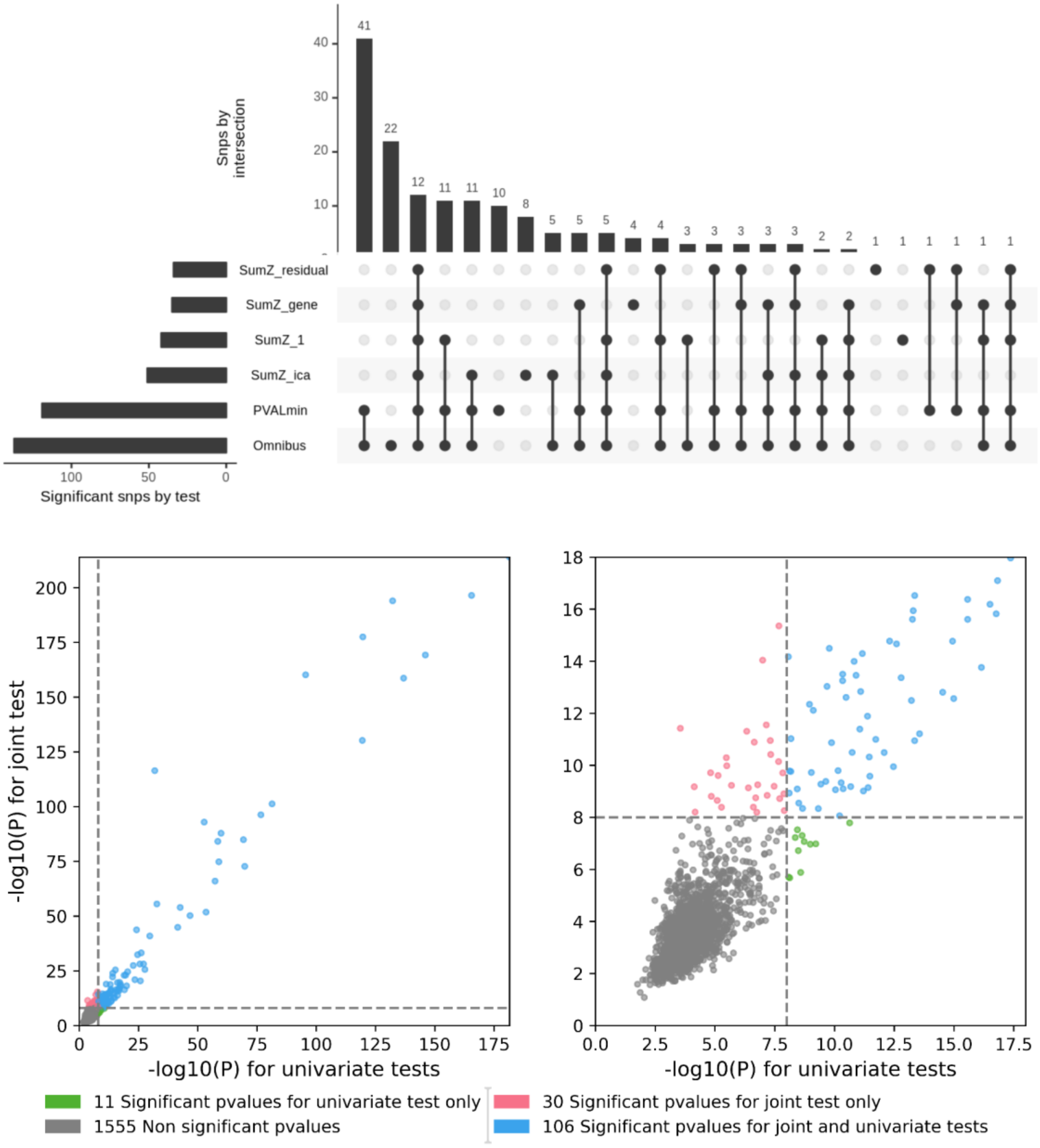

#### Figure S20. Signal comparison for Neurology phenotypes.

The upper panel shows independent variants detected across phenotype groups and across approaches represented as an *UpSetR* visualization. Matrix lines correspond to a test, each column to a set of significant variants. For each set, the test for which variants are significant are represented with a black dot on the test line. The barplot on the left of the matrix represents the number of significant independent signals detected by each approach. The barplot on the top of the matrix represents the cardinality of the sets. The sets are ordered by cardinality from the largest to the leftmost to the smallest to the rightmost. The bottom panels show quadrant plots, i.e. the -log10(*p*-value) for the most significant SNP per region for the *Omnibus* test as a function of the -log10(*p*-value) for the most significant SNP per region across all univariate GWAS. Complete results are presented in the left panel, and a zoom around the genome-wide significance threshold is presented on the right panel.

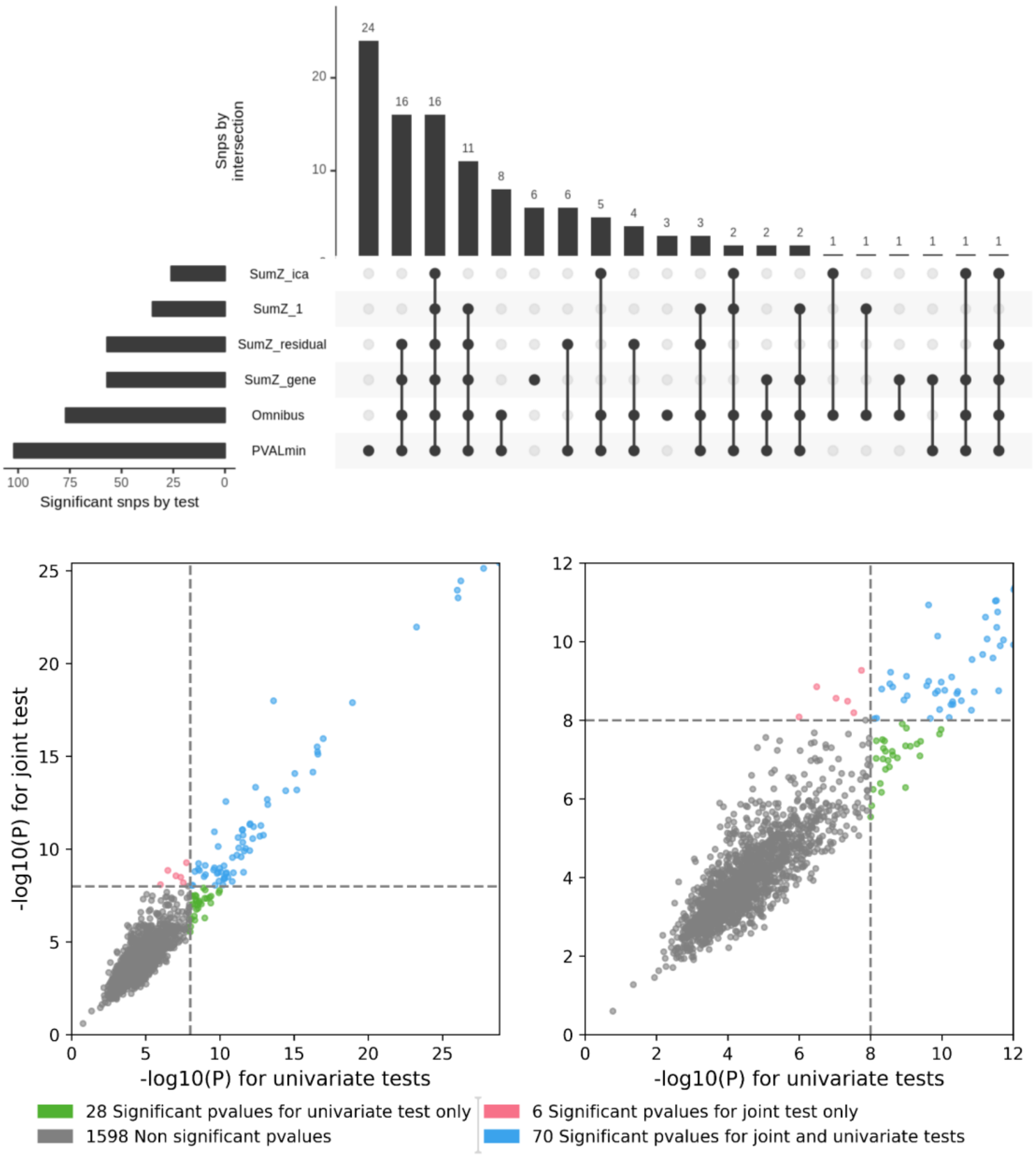

#### Figure S21. Proportion of tissue type enriched by phenotype group.

The union of enriched tissue by phenotype set were mapped to their larger anatomical category (Tissue type) in GTEx. To simplify the visualization, anatomical categories that did not explain at least 5% of any of the phenotype group were regrouped under the “Other” category. In each group, anatomical category representing less than 1% were approximated to 0. a) results obtained with variants detected with univariate test and b) results with variants detected with multivariate tests.

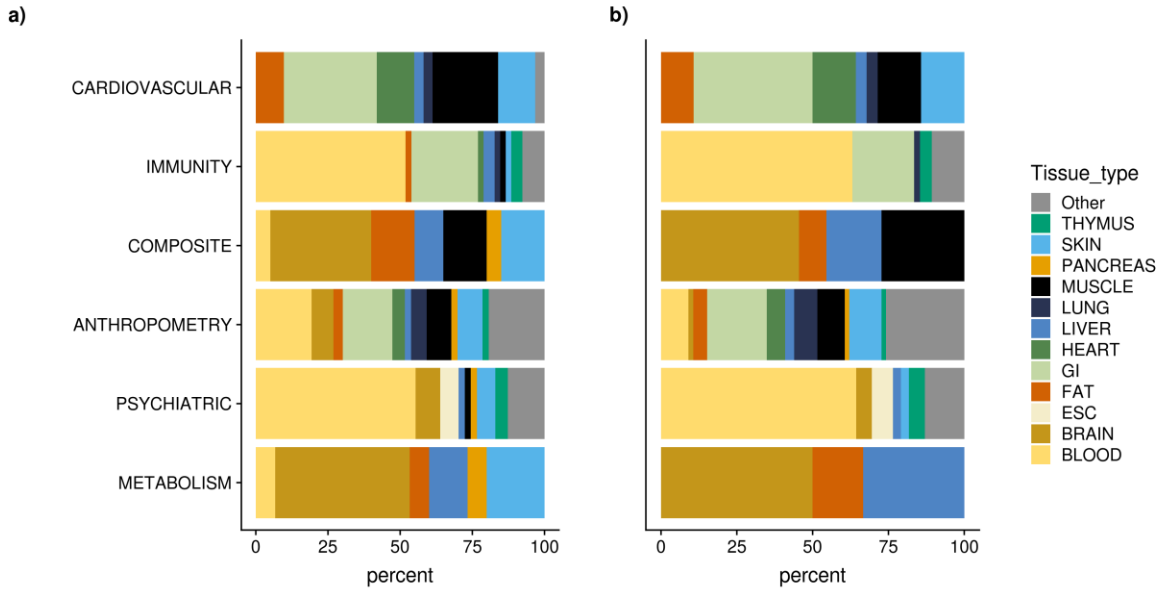

#### Figure S22. Clustering criterion by number of clusters and phenotype set.

In all panels, the x-axis is the number of clusters derived using the union of significant SNPs in the *Omnibus*, $SumZ_{genet}$ and univariate tests. On the left column, the y-axis represents the BIC (Bayesian Information Criteria). On the right column, the y-axis is the Silhouette criteria (see Methods). Each line corresponds to a different group of phenotypes.

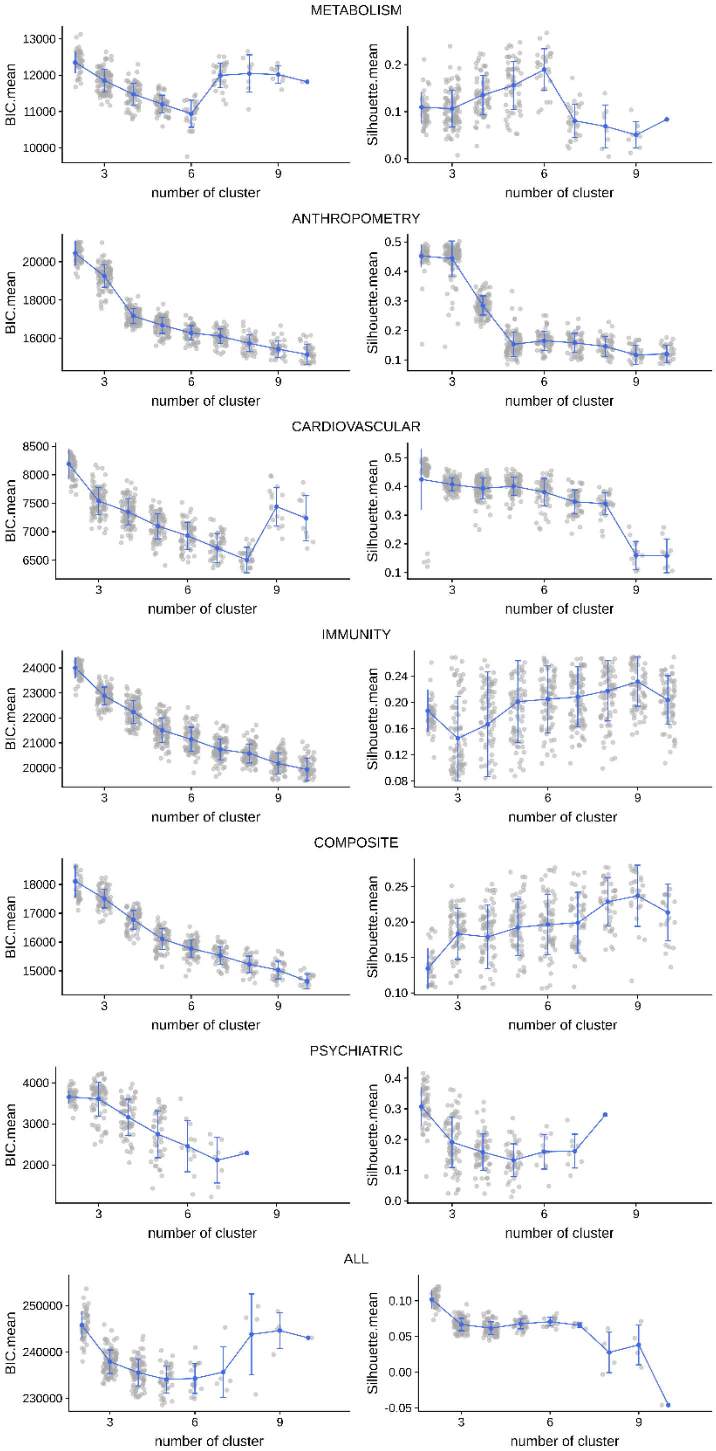

#### **Figure S23. Clustering for the *Metabolism* set using SNPs detected by univariate analysis only.**

The x-axis is the number of clusters derived using the significant SNPs univariate tests. On the left column, the y-axis represents the BIC (Bayesian Information Criteria). On the right column, the y-axis is the Silhouette criteria (see Methods).

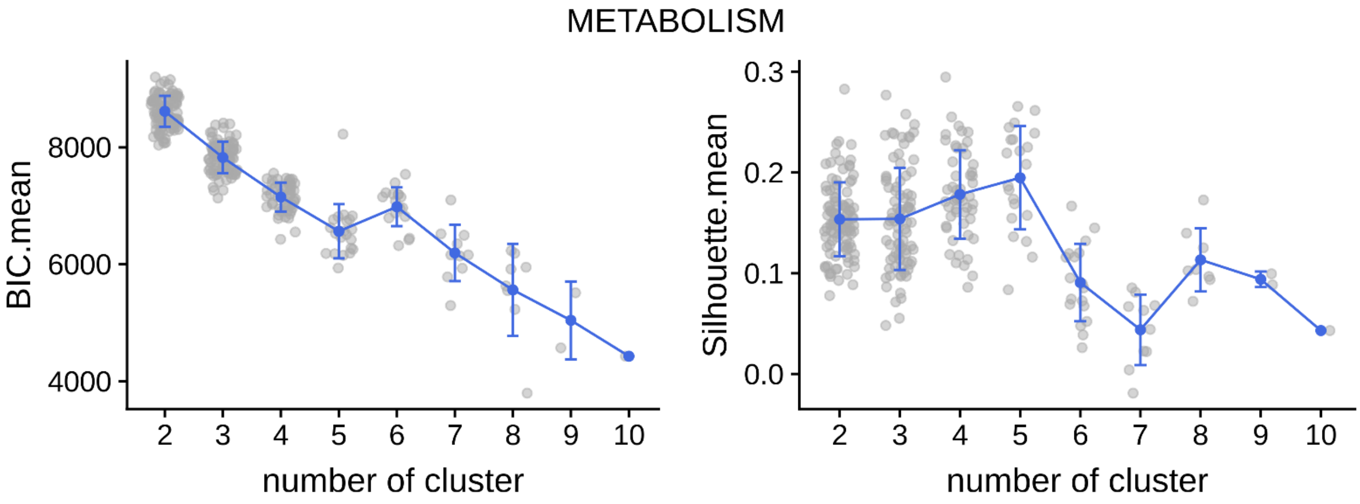

#### Figure S24. Alluvial plot and heatmaps for the Anthropometry GWAS set

On the left panel, the alluvial plot represents the re-assignment of SNPs from univariate analysis to clusters. To emphasize the relative genetic contribution to phenotypes, SNPs from each phenotype block were weighted by their variance explained to that phenotype. On the right panel, the heatmap represents the multi-trait signatures. Each line is a SNPs, each column, a trait. The gradient of color represents the strength of the Z-scores.

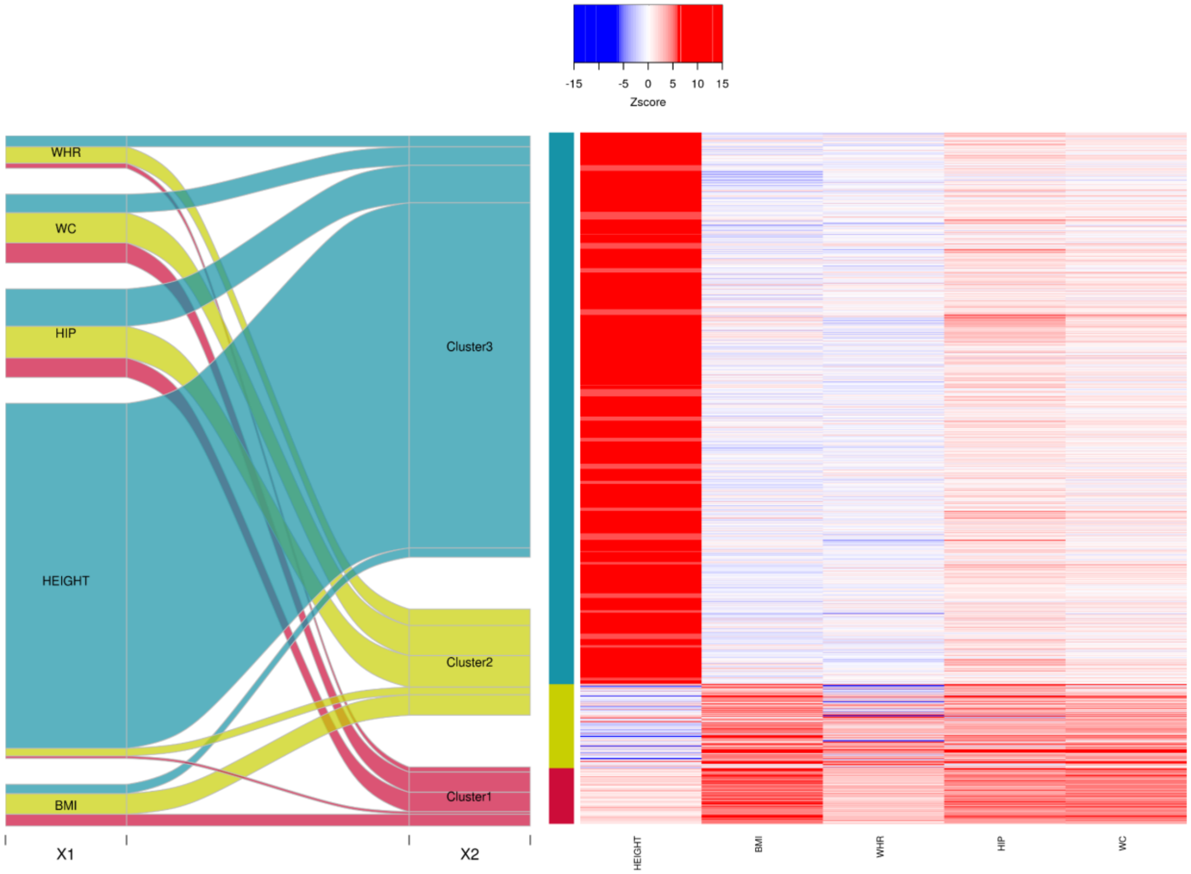

#### Figure S25. Alluvial plot and heatmaps for the Cardiovascular GWAS set

On the left panel, the alluvial plot represents the re-assignment of SNPs from univariate analysis to clusters. To emphasize the relative genetic contribution to phenotypes, SNPs from each phenotype block were weighted by their variance explained to that phenotype. On the right panel, the heatmap represents the multi-trait signatures. Each line is a SNP, each column, a trait. The gradient of color represents the strength of the Z-scores.

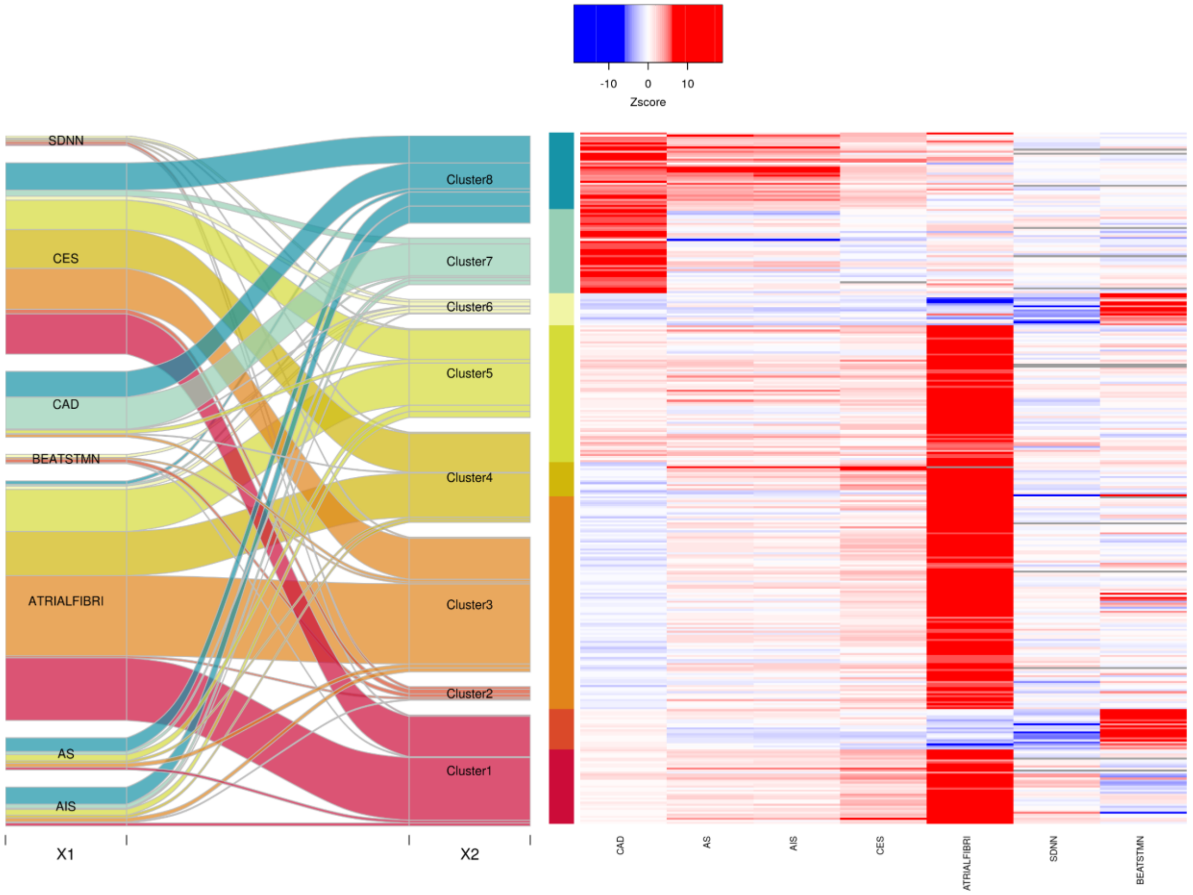

#### Figure S26. Alluvial plot and heatmaps for the Immunity GWAS set

On the left panel, the alluvial plot represents the re-assignment of SNPs from univariate analysis to clusters. To emphasize the relative genetic contribution to phenotypes, SNPs from each phenotype block were weighted by their variance explained to that phenotype. On the right panel, the heatmap represents the multi-trait signatures. Each line is a SNP, each column, a trait. The gradient of color represents the strength of the Z-scores.

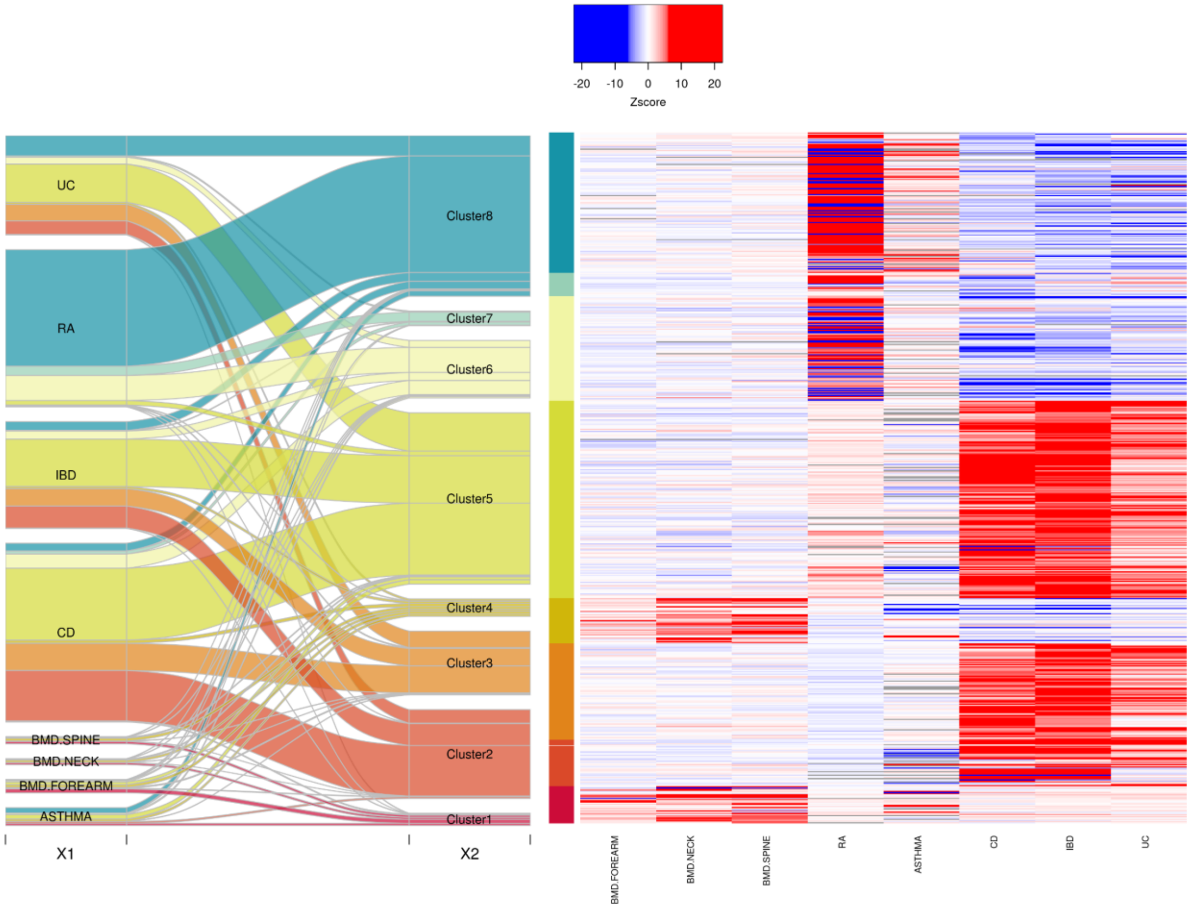

#### Figure S27. Alluvial plot and heatmaps for the Composite GWAS set

On the left panel, the alluvial plot represents the re-assignment of SNPs from univariate analysis to clusters. To emphasize the relative genetic contribution to phenotypes, SNPs from each phenotype block were weighted by their variance explained to that phenotype. On the right panel, the heatmap represents the multi-trait signatures. Each line is a SNP, each column, a trait. The gradient of color represents the strength of the Z-scores.

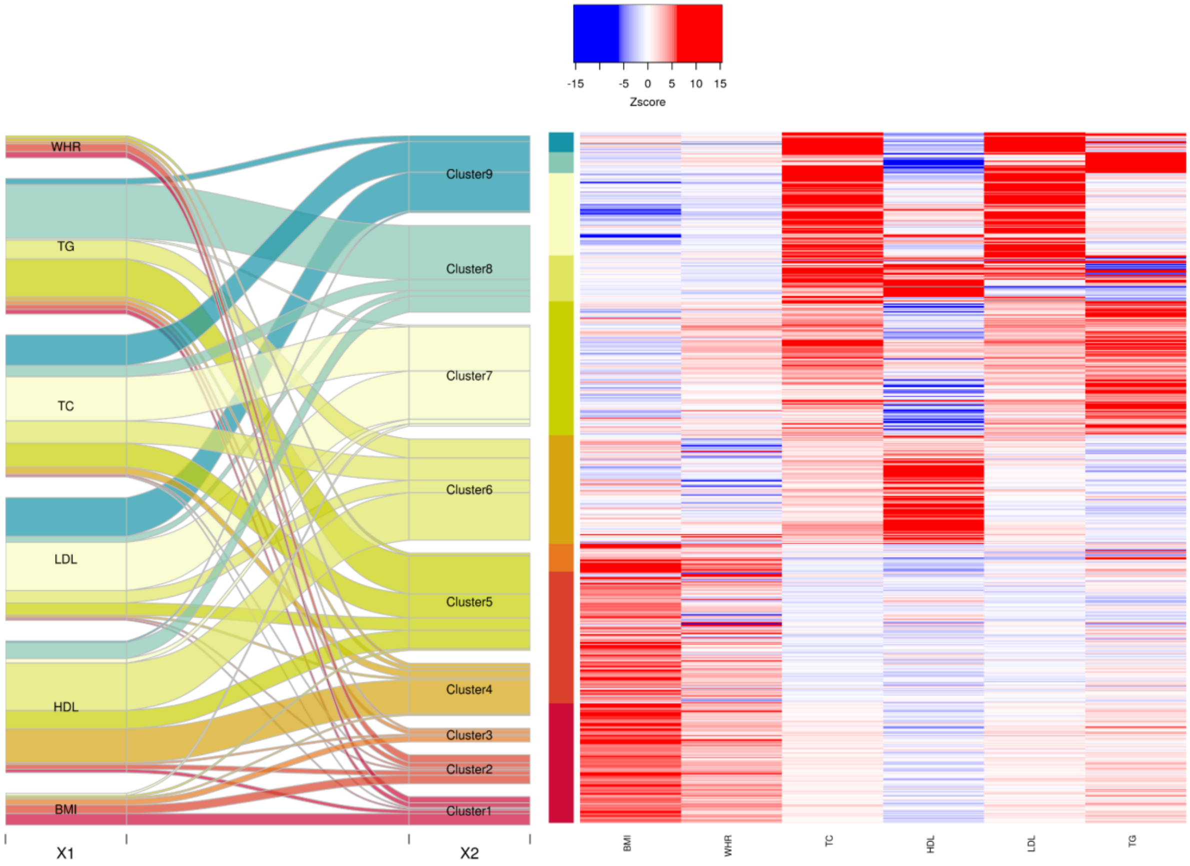

#### Figure S28. Alluvial plot and heatmaps for the Neurology GWAS set

On the left panel, the alluvial plot represents the re-assignment of SNPs from univariate analysis to clusters. To emphasize the relative genetic contribution to phenotypes, SNPs from each phenotype block were weighted by their variance explained to that phenotype. On the right panel, the heatmap represents the multi-trait signatures. Each line is a SNP, each column, a trait. The gradient of color represents the strength of the Z-scores.

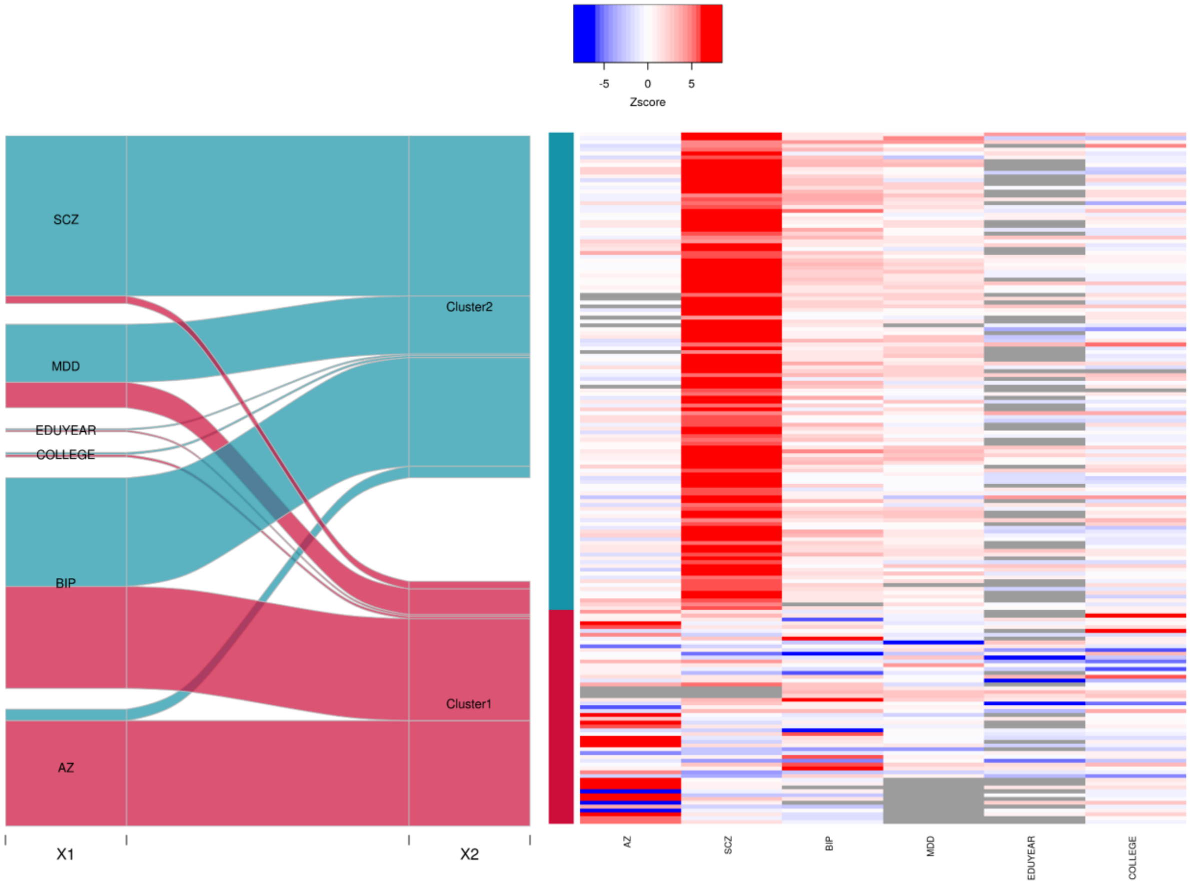

#### Figure S29. Alluvial plot and heatmaps for the Metabolism GWAS set

On the left panel, the alluvial plot represents the re-assignment of SNPs from univariate analysis to clusters. To emphasize the relative genetic contribution to phenotypes, SNPs from each phenotype block were weighted by their variance explained to that phenotype. On the right panel, the heatmap represents the multi-trait signatures. Each line is a SNP, each column, a trait. The gradient of color represents the strength of the Z-scores.

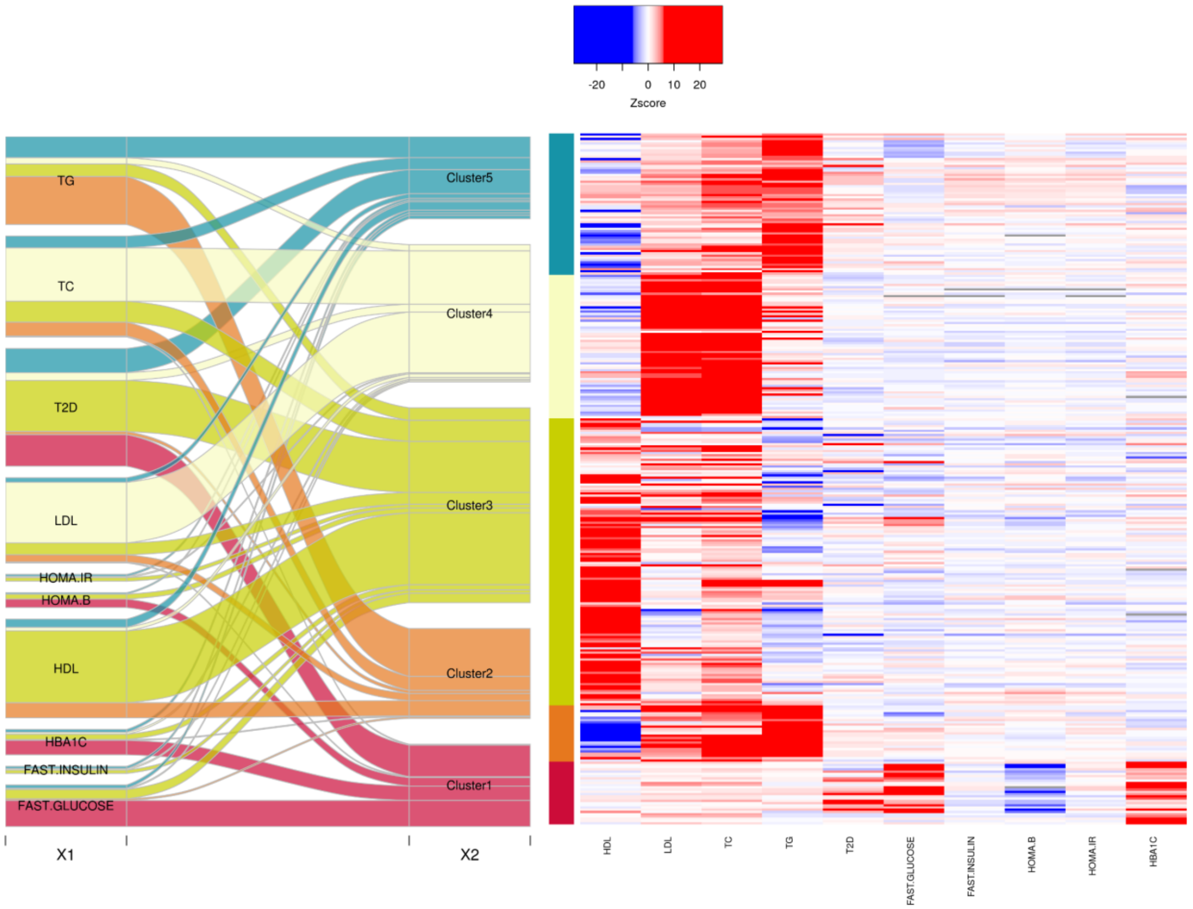

#### Figure S30. Alluvial plot and heatmaps for all GWAS combined

On the left panel, the alluvial plot represents the re-assignment of SNPs from univariate analysis to clusters. To emphasize the relative genetic contribution to phenotypes, SNPs from each phenotype block were weighted by their variance explained to that phenotype. On the right panel, the heatmap represents the multi-trait signatures. Each line is a SNP, each column, a trait. The gradient of color represents the strength of the Z-scores.

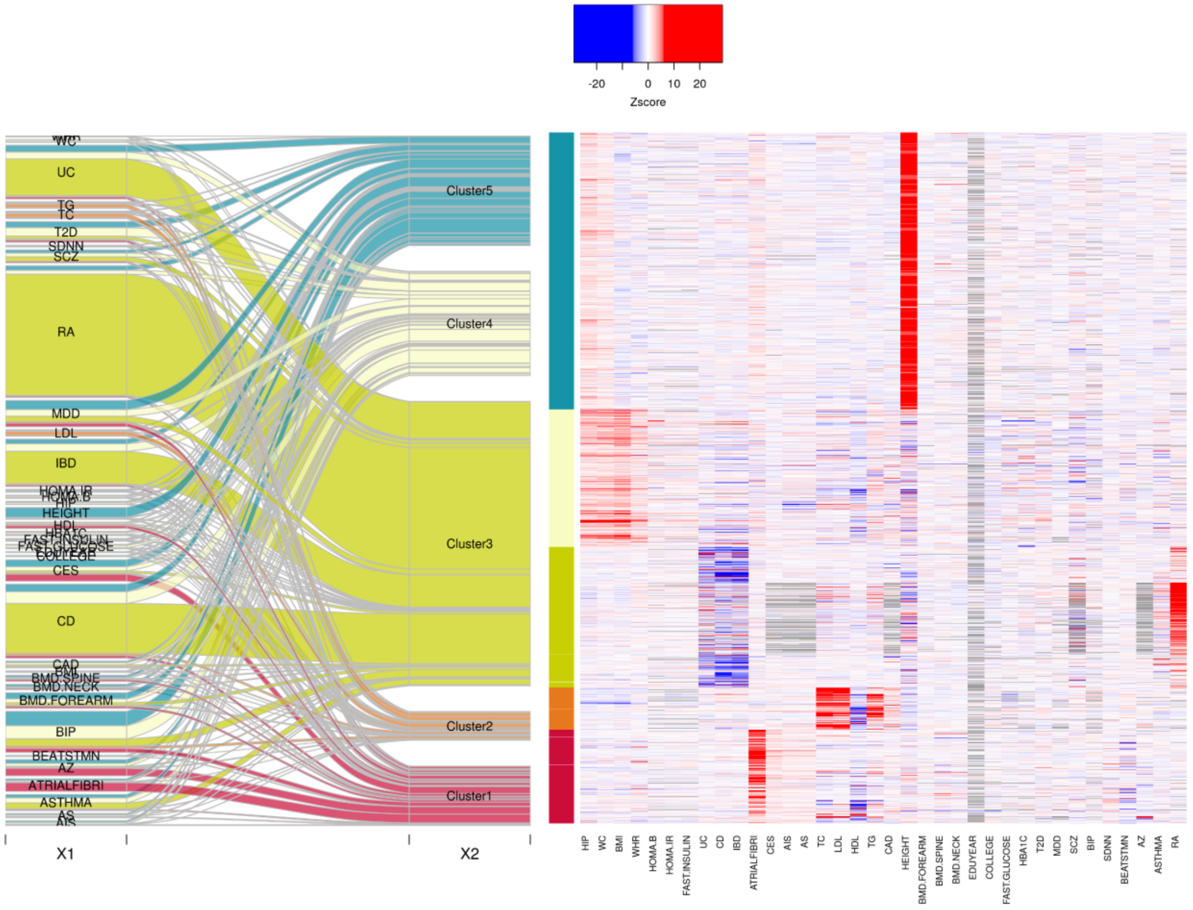

#### Figure S31. Heatmaps for the *Metabolism* SNPs for cardiovascular traits and Alzheimer disease

the heatmap represents the impact multi-trait signatures detected on the METABOLISM set on disease that have been linked to hyperlipidemia and diabetes. Each line is a SNP, each column, a trait. The gradient of color represents the strength of the Z-scores. Note that,

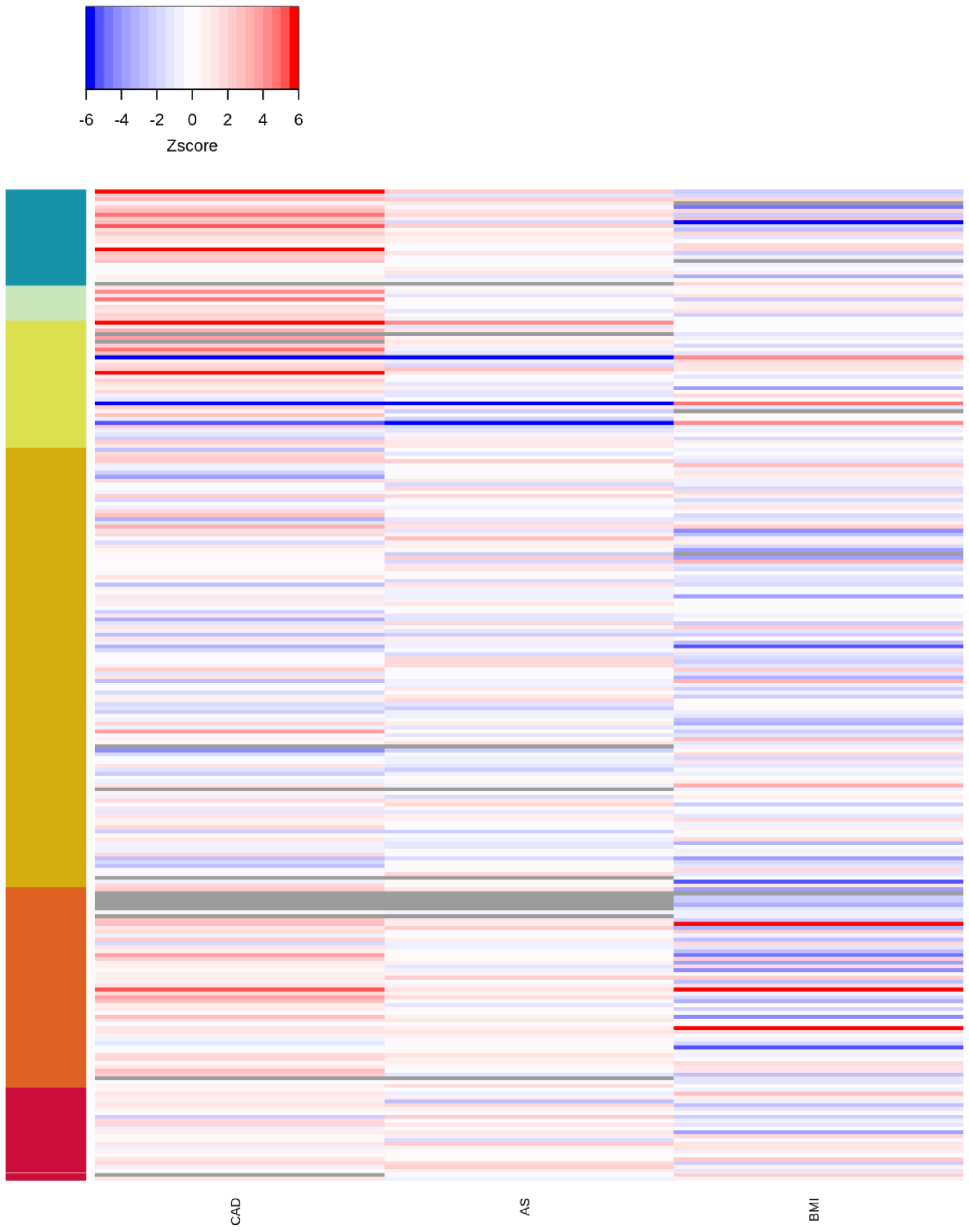
